# Neural correlates of motor sequence learning and enhanced offline consolidation in 7- to 11-year-old children

**DOI:** 10.1101/2025.08.15.670597

**Authors:** Anke Van Roy, Ainsley Temudo, Vincent Koppelmans, Kerstin Hödlmoser, Genevieve Albouy, Bradley R. King

## Abstract

**Intro:** Many daily activities involve series of interrelated movements and thus the capacity to learn new motor sequences is vital for everyday functioning. Although learning new skills is especially prominent throughout childhood, remarkably few studies have examined the neural underpinnings of motor sequence learning (MSL) and memory consolidation in children.

**Methods:** Twenty-two children (7-11 years) and 23 adults (18-30 years) underwent functional magnetic resonance imaging while completing two sessions of a MSL task, separated by a 5-hour offline period of wakefulness.

**Results:** Analyses of the behavioral data revealed comparable initial learning in children and adults. However, and consistent with previous research, children exhibited superior motor memory consolidation over the 5-hour offline epoch. Neuroimaging analyses revealed that children exhibited smaller modulations in brain activity between task and rest epochs in a widespread network, including the sensorimotor cortex, supplementary motor area, cerebellum, putamen and regions associated with the default mode network. Similar levels of activity during task and rest epochs in the hippocampus, dorsolateral prefrontal cortex and somatosensory cortex were associated with better motor memory consolidation in children.

**Conclusion:** Results potentially suggest that the continued engagement of the developing brain during interleaved rest contributes to the childhood advantage in motor memory consolidation.

## 1. Introduction

Many daily activities performed throughout the human lifespan rely on the successful execution of movement sequences (e.g., eating with utensils, tying shoelaces, typing a text message, etc.). Consequently, the capacity to acquire novel movement sequences is vital throughout one’s life and thus an understanding of the underlying neural processes is critical. Prior research has extensively examined the behavioral and neural correlates of motor sequence learning (MSL) in healthy young adults (∼18-35 years; see Albouy et al., 2013; Doyon et al., 2018; King et al., 2017; Krakauer et al., 2019 for reviews), an age group that is often considered the model of optimal functioning (e.g., Leversen et al., 2012). In brief, the initial learning of a motor sequence is characterized by rapid performance improvements and is supported by distinct neural networks that present different dynamical patterns of activity (Albouy, King, et al., 2013; Doyon, Bellec, et al., 2009; Hikosaka et al., 2002). An associative brain network that includes the hippocampus, associative territories of the striatum and cerebellum, as well as frontal and parietal areas is heavily recruited during early task practice and exhibits subsequent decreases as a function of learning. Conversely, activity in a sensorimotor network encompassing motor cortical areas, the sensorimotor striatum and cerebellum increases as learning progresses.

The “offline” (i.e., in the absence of active task practice) periods that follow initial learning afford the opportunity for the recently acquired memory trace to be consolidated into a stable, long-term form. Previous neuroimaging studies have revealed that changes in brain activity across an offline consolidation epoch often depend on whether this interval includes sleep. Specifically, activity in hippocampo-frontal and motor cortical areas increases following offline periods that include sleep (Albouy, King, et al., 2013; Debas et al., 2010; Walker et al., 2005), whereas awake consolidation appears to be supported by increases in primary motor and parietal cortices (Albouy et al., 2008; Robertson et al., 2005; Wang et al., 2024). Changes in striatal activity are less dependent on vigilance state, as increases in activity have been observed across offline periods that include sleep as well as only wakefulness (Albouy et al., 2013). The research reviewed above characterized the neural correlates associated with the traditional “slow” consolidation process that takes place on the “macro-offline” timescale (i.e., hours to days following initial training). More recent work has suggested that consolidation also occurs on a faster “micro-offline” timescale (i.e., the seconds to minutes that constitute the brief rest periods between practice blocks; Bönstrup et al., 2019; Buch et al., 2021; Du et al., 2017; Gann et al., 2023; Jacobacci et al., 2020). Offline processing on the micro-timescale has been linked to activity in the hippocampus and caudate nucleus (Buch et al., 2021; Chen et al., 2024; Gann et al., 2023; Jacobacci et al., 2020; Sjøgård et al., 2024).

In contrast to the abundance of prior research that has examined the neural underpinnings of motor sequence learning in young adults, remarkably few studies have done so in children and these have largely been limited to the initial learning session [e.g., (Hedenius & Persson, 2022; Thomas et al., 2004)]. Results revealed age-group differences in the activation of motor-related brain regions during initial learning, with children and adults exhibiting greater recruitment of subcortical (e.g., putamen) and cortical areas (e.g., premotor cortex), respectively. The limited number of neuroimaging studies in pediatric samples is especially striking given that childhood is a time when learning new skills is very prominent and the developing brain exhibits heightened plasticity (see Ismail et al., 2017 for a review) that may provide children with a unique capacity for enhanced skill development. For example, whereas previous research has produced mixed results with respect to differences in initial learning between children and adults (Adi-Japha et al., 2014; Fischer et al., 2007; Janacsek et al., 2012; Meulemans et al., 1998; Salehi et al., 2016; Wilhelm et al., 2008) research from our team and others have provided behavioral evidence suggesting that children (∼7-12 years) exhibit superior macro-offline consolidation following a period of post-learning wakefulness as compared to adults (Adi-Japha et al., 2014; Ashtamker & Karni, 2013; Dorfberger et al., 2007; Van Roy et al., 2024). There is also evidence to suggest that such a childhood advantage may not be limited to the macro-offline timescale. Young children (i.e., 6 year-olds) also showed greater performance gains across the micro-offline rest periods interspersed with blocks of active task practice during an initial learning session (Du et al., 2017). The neural correlates of these behavioral advantages in offline processing remain unknown.

The aim of the current research was to examine the neural substrates supporting motor sequence learning as well as the developmental advantage in awake offline consolidation processes in children. To do so, 7- to 11-year-old children and 18–to 30-year-old adults performed a MSL task during two learning sessions, separated by a 5-hour offline period of wakefulness. Using magnetic resonance imaging (MRI), functional brain images were acquired during both sessions, affording the assessment of age group differences in task-related brain activation during learning as well as changes in activity across the 5-hour macro-offline period. Similar to our previous behavioral study (Van Roy et al., 2024), we hypothesized that learning dynamics (i.e., block-to-block performance changes) during initial learning would be comparable in the two age groups; however, children were predicted to have smaller micro-online and larger micro-offline gains as compared to adults. During this initial acquisition phase, and in line with Thomas et al. (2004), children were expected to exhibit greater task-related activation of the striatum and hippocampus, whereas adults would show greater cortical brain activity. We predicted that these differences in task-related brain activity would be associated with the behavioral micro-online and -offline performance gains. Following initial learning, we hypothesized that – in line with our prior study (Van Roy et al., 2024) – children would exhibit a behavioral advantage in performance gains across the 5-hour offline period. At the neural level, we expected that both children and adults would show increases in activity in the hippocampus and striato-motor regions across the consolidation interval, with larger increases in children. These activity changes were anticipated to be related to the greater macro-offline performance changes in children.

## 2. Methods and materials

Data collection and analysis plans were registered in April 2024 via Open Science Framework and can be accessed at https://doi.org/10.17605/OSF.IO/SW6D8. Any deviations from the registered procedures are marked in this text with an asterisk and additional details for the deviations are provided in Appendix 1 as part of the Supplementary Material. Any analytic procedures that were not included in the registration are labeled as exploratory in this text. Note that prior to the registration, 27 (out of 45 included) datasets were acquired and 18 were processed using preliminary analytic scripts. These analyses served to assess data quality (i.e., keypresses and their timing were logged adequately, acquired images were of sufficient quality, and that saved and exported data were complete) and were also used as pilot data for funding applications. The data that support the findings of this study are openly available on Zenodo at https://doi.org/10.5281/zenodo.16891677. The study protocol was approved by the University of Utah Ethics Committee (IRB_00155118) and experimental procedures were conducted according to the declaration of Helsinki (2013).

### 2.1 Participants

A convenience sample of seemingly healthy children (operationally defined as 6-12 years) and adults (18-30 years) of all genders was recruited via advertisements posted in public spaces and on relevant websites. Eligibility was assessed based on self- or parental report via an online screening questionnaire. Exclusion criteria were: 1) left-handed; 2) history of medical, neurological, psychological, or psychiatric conditions, including depression and anxiety; 3) consumption of psychotropic or psychoactive medications; 4) self-reported indications of abnormal or irregular sleep; 5) mobility limitations of the fingers/hands that could prevent successful motor task completion; 6) extensive training in a musical instrument requiring dexterous finger movements (e.g., piano, guitar) or as a typist; 7) previous participation in a research experiment employing a similar motor sequence learning task; and, 8) presence of MRI contraindications (including claustrophobia). Adult participants and parents of child participants gave informed consent, and children 7 to 12 years of age gave informed assent (i.e., children of 6 years were not required to provide assent). Note that although 6- to 12-year-old children were recruited, our final sample of child participants was limited to 7-11-year-olds (see Appendix 1 for information on attrition).

Sample size estimation was conducted *a priori* and with the software G*Power (Faul et al., 2007). The effect size of interest was set to d = 0.76 and computed as the average effect size from the behavioral results of two published studies assessing motor memory consolidation over post-learning wake intervals in children and adults. Specifically, the comparison in 5hr offline performance changes between children and young adults in Van Roy et al., (2024) corresponded to Cohen’s d = 0.6. In Ashtamker and Karni (2013), Cohen’s d was estimated to be 0.92 based on the data depicted. To detect a childhood advantage in motor memory consolidation across a period of wakefulness using a one-tailed independent samples t-test with an effect size of d = 0.76, α = 0.05 and a desired power of 0.80, the desired sample size was 46 participants (23 children; 23 adults). A total of 61 participants (36 children, 25 adults) enrolled in the study. Sixteen of these participants (14 children) did not complete the experiment (see flow diagram for inclusion and exclusion in Appendix 1). Due to a rate of exclusion that was slightly higher than expected for children, our final sample included 22 children and 23 adults and thus was one child short of the registered sample size. Note that a subset of participants was excluded from specific analyses due to excessive head motion or deviations from the experimental protocol (see Appendix 1 for details). All figure and table captions in the main text and Supplementary Tables explicitly state the sample size included in the corresponding analysis.

Potential age group differences in participant characteristics, sleep, vigilance and protocol timing were assessed using independent samples t-test or Mann-Whitney U tests, depending on whether the data were normally distributed. These measures included morningness/eveningness (self-report), self-reported sleep quantity and quality of the night prior to the experimental day, time of testing of each session and the duration of the offline period between initial training and the 5-hour delayed retest. Furthermore, objective and subjective measures of vigilance at the time of testing (see Section 2.2) were compared between age groups and sessions using a group (2 levels: children vs. adults) by session (2 levels: training vs. retest session) mixed ANOVA.

### 2.2 Experimental procedure

Participation in this research consisted of an initial familiarization visit followed by two experimental sessions that took place on the same day (see Figure 1). Participants were asked to have a good night of sleep the night prior to each testing day. The first visit consisted of a short familiarization session (∼1 hour) that was scheduled based on participant availability but restricted to between 9 am and 6 pm. The purpose of this session was to familiarize participants to the research team, MRI environment and motor tasks. A detailed description of the familiarization procedure is provided in Appendix 1. At the end of the familiarization session, participants were given a sleep diary on which they were asked to record their sleep times starting three nights prior to the experimental visit. Ideally, the experimental session occurred at least two days following the familiarization session, but this was not always possible given the schedules of two participants. The duration between these two visits ranged from 0 (one adult) to 29 days (medians for children and adults were 6.5 and 4 days, respectively).

**Figure 1.**
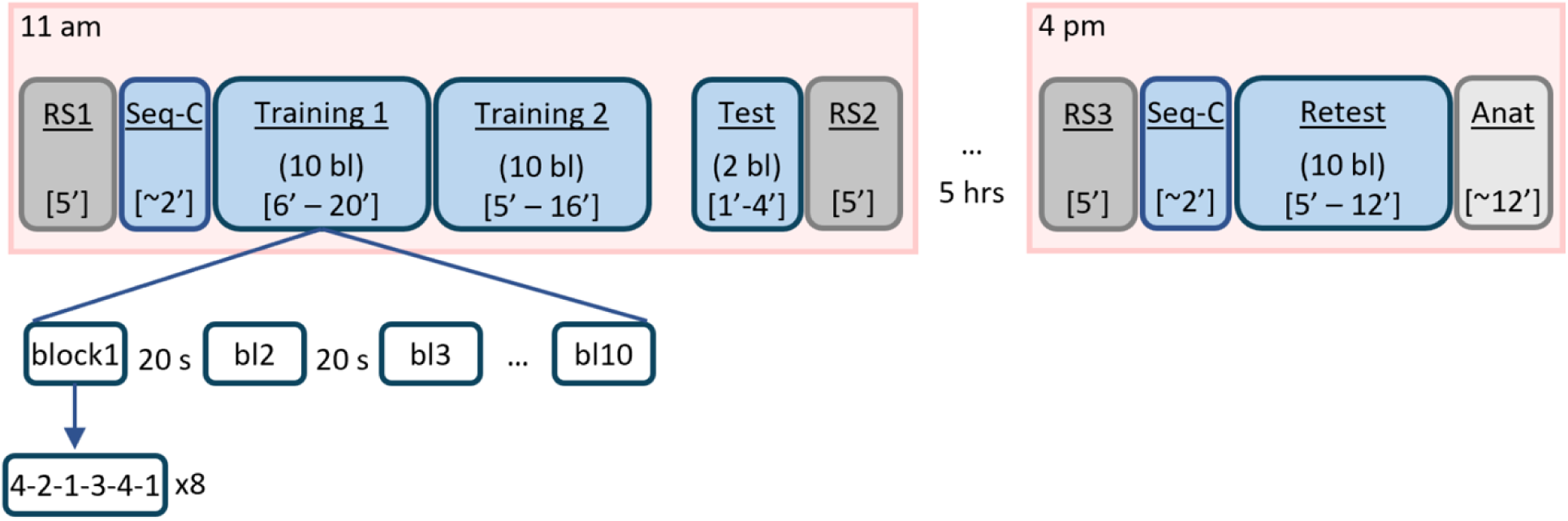
Design for the experimental sessions. Initial training on the finger tapping task (FTT) was divided into training 1, training 2 and test runs. The test run provided a behavioral measure of end-of-training performance after the dissipation of fatigue (Pan & Rickard, 2015). Resting-state (RS) scans were acquired prior to and following the FTT task runs (RS data not included in this manuscript). Following a 5-hour macro-offline consolidation interval, participants completed another RS scan followed by a retest of the motor task. In all FTT runs, blocks of task practice were alternated with 20s rest epochs, affording the assessment of micro-online and -offline learning. hrs = hours. Seq-C = sequence-check. Anat = anatomical T1- and T2-weighted images. Anatomical data were originally acquired at the end of the retest session (i.e., for the first 10 children and 15 adults). However, to check that participants were comfortable with the real scanning environment prior to the experimental visit, the acquisition of the structural images was moved to the end of the familiarization session for the last 12 children and 8 adults included in the analyses. Durations of runs are indicated between brackets. Note that ranges were provided for task runs since durations depended on individual performance. Pink shaded regions depict instances when participants were in the MRI scanner. BOLD images were acquired during RS and FTT task runs, except for the sequence-check runs.

The primary experimental day consisted of two sessions, herein referred to as training and retest, completed approximately 5 hours apart (see Table 1 for details on the duration of this between-session interval). The training session took place in the morning between 9 am and 1:30 pm. The 5-hour delayed retest session was thus scheduled between 2 pm and 6:30 pm, affording the assessment of motor memory consolidation after an interval of post-learning wakefulness.

**Table 1.** Participant demographics and sleep and vigilance scores for each age group.

| Variable |  | Children | Adults | p-value |
| --- | --- | --- | --- | --- |
| *n* |  | 22 | 23 |  |
| Female/Male/Non-Binary (*n*) |  | 14/7/1 | 16/6/1 |  |
| Age (years) |  | 9.5 (1.3) | 23.4 (3.3) |  |
| Race/Ethnicity | *Asian* | 1 | 2 |  |
|  | *Black/African American* | 0 | 1 |  |
|  | *White* | 18 | 17 |  |
|  | *Mixed Race* | 2 | 2 |  |
|  | *Hispanic / Latino* | 1 | 1 |  |
| M/E preference |  | 2.8 (0.7) | 2.7 (0.9) | 0.582 |
| Training session | *Time of testing* | 10:49 (49) | 10:53 (62) | 0.598 |
|  | *Sleep quality* | 5.2 (0.7) | 4.9 (0.8) | 0.173 |
|  | *Sleep duration (hrs.)* | 9.83 (0.83) | 8.3 (0.9) | < 0.001* |
|  | *SSS score* | 2.3 (0.8) | 2.0 (0.7) | 0.346 |
|  | *PVT score (ms)* | 393.7 (66.1) | 298.0 (24.6) | < 0.001 * |
| Retest session | *Time of testing* | 16:28 (45) | 16:19 (59) | 0.295 |
|  | *Offline period (hrs.)* | 5.1 (0.7) | 5.1 (0.4) | 0.574 |
|  | *SSS score* | 2.3 (0.8) | 2.4 (1.1) | 0.836 |
|  | *PVT score (ms)* | 393.0 (54.9) | 301.6 (29.7) | < 0.001* |

Values for *n*, gender and race/ethnicity represent number of participants. For other values, the numbers represent the mean, except for the PVT (median), with standard deviation (SD) in parentheses. Gender was determined by asking what gender the participant identified with the most. Morningness/Eveningness (M/E) was included as an exploratory variable (i.e., not included in registration) to assess age-related differences in circadian preferences and was defined on a 5-point Likert scale (i.e., 1 = extreme morning person, 5 = extreme evening person). The average time of testing reflects the start of FTT practice during the respective session, and is specified in 24-hour time notation, with the SD in minutes. The offline period represents the time period between the end and start of practice of the test and retest runs of the FTT, respectively. Sleep quality was defined on a 6-point Likert scale (i.e., 1 = very bad, 6 = very good). SSS = Stanford Sleepiness Scale (Maclean et al., 1992), with higher numbers indicative of increased sleepiness. The PVT score indicates the median simple RT across the 30 trials included in the analyses. The p-values resulted from independent samples t-tests or Mann-Whitney U-test assessing group differences, depending on whether the assumption of normality was met or violated, respectively. See Appendix 2 for full statistical output.

At the beginning of the training session, participants returned their completed sleep diary and re-verified that they did not present any MRI contra-indications. This was followed by answering brief questionnaires reporting on their sleep patterns in the previous 24 hours, their subjective feelings of alertness (Stanford Sleepiness Scale; Maclean et al. 1992), intake of medicine in the past 3 days, as well as any exercise and consumption of alcohol and/or recreational drugs in the last 24 hours (alcohol and drug intake for adult participants only). Subjects also completed a modified version of the Psychomotor Vigilance Task (PVT; Dinges & Powell, 1985) to obtain an objective assessment of vigilance/alertness at the time of testing. Briefly, participants were asked to fixate on the middle of the computer screen. Following varying delay intervals, a visual stimulus appeared, and participants were instructed to respond by pressing the space bar as fast as possible. A shortened version of the PVT was administered (validated in Basner et al., 2011 and Loh et al., 2004), consisting of 35 trials, the first 5 of which were considered practice trials and excluded from analyses. The response times (i.e., time between appearance of stimulus and the keypress; RT) were logged and used to assess vigilance (i.e., higher RTs indicate lower vigilance). Following the PVT, participants completed 5 blocks of the pseudorandom SRTT in the mock scanner to assess general motor execution without engaging sequence learning processes, as well as the FTT habituation procedures at a table outside the mock scanner. These tasks were administered to ensure participants were comfortable with completing the motor tasks and the simulated MR environment prior to entering the real MR environment and initiating the portion of the protocol used for data analyses. Accordingly, data from the SRTT and familiarization procedure were not analyzed.

Participants were subsequently positioned in the 3-Tesla MR scanner with an MR compatible keyboard on each hand. Following a pre-training resting-state run (RS1), participants were introduced to the experimental sequence (4-2-1-3-4-1, where 1 is the left middle finger, 2 is the left index finger, 3 is the right index finger and 4 is the right middle finger) during a short sequence-check. Specifically, participants were provided with this sequence of keypresses for the first time and were asked to perform the sequence slowly and as accurately as possible. This sequence-check ended when 3 correct sequences were performed consecutively and ensured that participants could perform the sequence prior to starting the training runs. Participants then completed the initial learning session of the FTT task (see section 2.3) while blood-oxygen-level-dependent (BOLD) images were acquired (see Section 2.5). This session was organized into 3 runs, two training runs of 10 blocks followed by a test phase of 2 blocks. We elected to decompose the training portion into 2 runs to provide participants, particularly young children, with the opportunity to have a brief rest and to communicate with the research team. The test run was administered approximately 1 minute after the second training run, affording the assessment of end-of-training performance following the dissipation of mental and physical fatigue (Pan & Rickard, 2015). The initial learning session concluded with a post-learning resting-state run (RS2). Participants were then permitted to leave the imaging facility and were instructed to avoid strenuous exercise as well as caffeine (i.e., coffee, tea, energy drinks, soda). Importantly, as our research question is focused on offline consolidation over wake intervals, participants were instructed to not sleep. Based on self-report, no participants slept during the offline consolidation period.

Participants returned to the imaging center approximately 5 hours after the end of the training session to complete the retest. Following vigilance assessments (i.e., PVT and SSS), participants were positioned back in the MR scanner. This retest session included a pre-retest RS run (RS3) and the sequence-check, followed by 10 blocks of the FTT.

Note that functional brain images were acquired during all FTT and RS runs, except for the sequence-check. Analyses of RS data are not included in this manuscript. Images acquired during the test run were not analyzed given the limited number of volumes. Anatomical data, consisting of T1- and T2-weighted images, was originally acquired at the end of the retest session (i.e., for the first 10 children and 15 adults). However, to check that participants were comfortable with the real scanning environment prior to the experimental visit, the acquisition of the structural images was moved to the end of the familiarization session for the last 12 children and 8 adults included in the analyses. Participants were allowed to watch a movie or listen to music during the structural scans.

### 2.3 Finger Tapping Task (FTT)

An explicit, bimanual finger tapping task (FTT) was employed, coded and implemented in the MATLAB Psychophysics Toolbox version 3 (Kleiner et al., 2006). The task was analogous to our earlier research (Dolfen et al., 2019), but was modified slightly for children (see Figure S2). Specifically, participants used the index and middle fingers of both hands to press four keys on specialized MR-compatible keypads (Celeritas fMRI Button Response System; Psychology Software Tools, Pittsburgh, PA, USA). The task consisted of alternating practice and 20-second rest blocks, indicated by green and red text, respectively. During each practice block, a green cross and a 6-element sequence of numbers corresponding to the fingers to press (i.e., 4-2-1-3-4-1) were displayed on the screen. Participants were asked to repeatedly tap the finger sequence on the keyboard as rapidly and accurately as possible. Each practice block consisted of 48 keypresses (i.e., ideally corresponding to 8 repetitions of the 6-element sequence) and ended once this number of keypresses was recorded. During the rest periods, the cross turned red, and the sequence of numbers was replaced by a fixed sequence of random symbols (i.e., @-!-$-%-&-#) displayed in red to avoid exposure to the sequence during rest. Participants were asked to focus on the red cross and to not move their fingers during these rest epochs. During performance of the FTT, the timing and number of each keypress were recorded.

### 2.4 Behavioral data analyses

For each practice block of the FTT, performance speed was computed as the median time to perform a transition between pairs of consecutive keypresses, and performance accuracy was computed as the percentage of correct transitions. A transition was classified as “correct” if two subsequent keys were pressed in the same order as presented in the sequence. Similar to our previous research (Van Roy et al., 2024) and to account for expected age group differences in movement speed and accuracy independent of sequence learning (i.e., baseline differences; Meulemans et al. 1998; Sugawara et al. 2014; Salehi et al. 2016), outcome variables from task runs were normalized relative to outcome variables from the first training block. Specifically, for each outcome measure and each block of the FTT runs, performance was divided by the median outcome measure of the first block of training 1. Per our registration, statistical analyses on these normalized measures are presented in the main text. For completeness, results based on exploratory analyses on the non-normalized outcome measures (i.e., transition time and % of correct transitions) and the variables computed using these measures (i.e., micro-online and -offline performance changes, macro-offline performance changes) are reported in Appendix 3.

All behavioral statistical testing described below was performed using R version 4.5.1 (Posit, PBC, Boston, Massachusetts, USA). Statistical tests were considered significant when p < 0.05. However, for completeness, we also highlight those effects when 0.05 < p < 0.1 as non-significant trends. For all mixed ANOVAs detailed below, Greenhouse-Geisser corrections were applied in the event of a violation of sphericity. Effect size measures include eta-squared, partial eta-squared or Hedge’s g, depending on the statistical test. Null hypothesis testing for all behavioral effects presented in the main text was supplemented by the computation of Bayes Factors (BF; BayesFactor R-package; see Appendix 1 for details), reflecting the likelihood of the observed data favoring an alternative model (e.g., evidence for differences among groups, blocks, etc.) relative to the null model (i.e., no differences).

To assess potential age group differences in initial encoding of the motor sequence (i.e., prior to the macro-offline period), learning dynamics (i.e., block-to-block changes) were compared between groups using separate mixed ANOVAs for performance speed and accuracy with *group* (i.e., children vs. adults) as between-subject factor, and *run* (i.e., training 1 vs. training 2) and *block* (i.e., 10 training blocks) as within-subject factors. Similarly, to assess age group differences in continued performance of the motor sequence following the 5h offline interval, performance dynamics during retest were compared between groups using separate mixed ANOVAs with *group* (i.e., children vs. adults) as the between-subject and *block* (10 retest blocks) as the within-subject factor.

To examine the consolidation process on a macro-timescale, an “offline gain” measure was computed for each outcome variable. Specifically, the difference in normalized performance from the end of training (average across the 2 post-learning test blocks) to the beginning of the retest (average across the first 2 blocks) was calculated. The resulting measure reflects the offline change in performance from the end of the initial training session to the retest session and thus, is used as a measure of memory consolidation across the post-learning period of wakefulness. Per our registration, one-tailed independent samples t-tests were used to assess age group differences in the offline gain measures.

To obtain an assessment of performance that captures both performance components (i.e., speed and accuracy) in a single measure, a general performance index (GPI) was computed as an exploratory outcome metric. GPI was computed at the block level, using equation 2.1 below (similar to Reverberi et al. 2023). Subsequently, offline gains were calculated for the GPI, as the difference in GPI from the end of training (average across the 2 post-learning test blocks) to the beginning of the retest (average across the first 2 blocks). The GPI across practice blocks as well as macro-offline changes in GPI are depicted in Appendix 3.

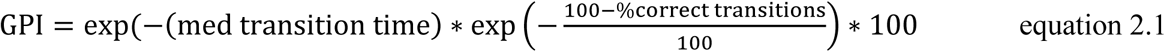

To assess performance changes on a micro-timescale, we distinguished between micro-online (i.e., within blocks of task practice) and micro-offline (i.e., between blocks of practice) performance improvements. As in our previous work (Van Roy et al., 2024), micro-*online* improvements were computed as the difference between the median normalized transition times of the first and last 6 transitions *within* each block. Micro-*offline* performance changes were computed as the difference between the median normalized transition times of the last 6 transitions of one block and the first 6 transitions of the subsequent block. The computations used six transitions to correspond to the five transitions between the six keypresses within a sequence, plus the transition between two consecutive sequences. To obtain a single micro-online and micro-offline value per training run, the micro-measures were averaged across task blocks or rest intervals, respectively, *within* training 1 and training 2 separately. Micro-online and -offline performance changes were subjected to separate 2 x 2 mixed ANOVAs with *group* (i.e., children vs. adults) as between-subject and training run (i.e., training 1 vs. training 2) as within-subject factor.

Registered analyses compared differences between age groups. Yet, to provide more fine-grained analyses of the development of motor memory consolidation behaviors, we also conducted exploratory analyses assessing age-related changes within our sample of child participants. Specifically, micro- and macro-offline performance changes were subjected to separate multiple regression analyses with age as a continuous independent variable. For both dependent measures, and consistent with our previous work (Van Roy et al., 2024), five potential fit options (i.e., single exponential, double exponential, linear, quadratic and power functions) were tested and the model with the lowest Akaike Information Criterion values was deemed as the best fitting model. Results from these regression analyses are provided in Appendix 3.

### 2.5 MRI data acquisition

MRI data were acquired on a Siemens MAGNETOM Vida 3.0T MRI system equipped with a 64-channel head coil. To minimize head motion, additional padding was placed around the head. For each participant, a high-resolution, T1-weighted structural image was acquired with a 3D MPRAGE sequence (axial; repetition time (TR) = 2500 ms; echo time (TE) = 2.98 ms; inversion time (TI) = 1070 ms; flip angle (FA) = 8°; 176 slices; field-of-view (FoV) = 256 x 256 x 176mm^3^; voxel size = 1.0 x 1.0 x 1.0 mm^3^; duration = 5:59 min). Parallel acquisition was conducted in the GRAPPA mode, with reference line phase encoding (PE = 32), and acceleration factors PE and 3D equal to 2 and 1, respectively. Analyses of the structural T2-weighted image are not included in this manuscript.

Resting-state (RS) and task-related fMRI data were acquired with an interleaved gradient EPI pulse sequence for T2*weighted images (SMS acceleration with multiband factor 5; TR = 797.0 ms; TE = 31.0 ms; flip angle = 59°; 55 transverse slices; voxel size = 2.5 x 2.5 x 2.5 mm^3^; field of view = 220 x 220 x 138 mm^3^). The task-related runs were set to a maximum of 1500 volumes, with a maximum duration of 20:03 minutes (including unsaved dummy pulses). These runs were terminated when participants completed all FTT practice blocks and thus the exact number of scans in each task run depended on each individual’s performance speed (median number of scans for children: 1096 for training 1, 803 for training 2, 690 for retest; adults: 588 for training 1, 508 for training 2, 486 for retest).

Gradient-recalled echo (GRE) fieldmaps were acquired prior to task performance during the training and retest sessions to correct for distortions due to magnetic field inhomogeneities. Specifically, two phase-encoded GRE images with different echo times (TR = 752 ms; TE_1_ = 5.19 ms, TE_2_ = 7.65 ms; flip angle = 90°; 55 transverse slices; voxel size = 2.5 × 2.5 × 2.5 mm^3^; field of view = 220 × 220 × 138 mm^3^) were collected. Phase-difference maps were calculated from the two GRE images using in-scanner processing.

### 2.6 fMRI data analysis

Prior to any processing, the DICOM images output by the SIEMENS scanner were converted to BIDS using the dcm2bids tool (Boré et al., 2023), implemented in Python 3.11.5. Additionally, the T1-weighted images were reoriented to the Montreal Neurological Institute (MNI) template to facilitate spatial normalization. The section below details the initial spatial preprocessing that was performed in fMRIPrep 24.1.0* (Esteban et al., 2019). This is followed by a description of the additional preprocessing and analyses performed in SPM12 (Wellcome Department of Imaging Neuroscience, London, UK) in Matlab R2023a.

#### 2.6.1 Spatial preprocessing in fMRIPrep

Initial spatial preprocessing was performed in fMRIPrep 24.1.0 (Esteban et al., 2019). Appendix 1 contains a detailed description of the spatial preprocessing, which was automatically generated by fMRIPrep and released under the CC0 license. Here, we provide a brief description of the preprocessing steps that are relevant for analyses presented in this manuscript. The T1-weighted (T1w) anatomical scan underwent correction for intensity inhomogeneities, followed by skull-stripping and tissue segmentation into gray matter (GM), white matter (WM), and cerebrospinal fluid (CSF). For each functional task run, head motion parameters were estimated (i.e., translation and rotation in the three motion planes; see Appendix 4 for information on the magnitude of head motion). Functional images were then distortion-corrected using the GRE fieldmaps and subsequently co-registered to the T1w image.

To reduce the detrimental effect of excessive head motion, images or entire runs where head motion exceeded the pre-defined threshold of 5mm (∼2 voxels) were removed. Details on head movement and data exclusion at the participant level are provided in Appendices 4 and 1, respectively. In brief, fMRI data from the training runs of 3 participants were truncated due to maximum head translation across any motion plane exceeding 5 mm. In the case of one child subject, head motion was large and consistent enough such that fMRI data from the entire Training 1 and Training 2 runs were excluded from analyses.

#### 2.6.2 Univariate fMRI analysis

Additional preprocessing of the MRI images and statistical testing were performed in SPM12 (Wellcome Department of Imaging Neuroscience, London, UK). A sample-based DARTEL (Diffeomorphic Anatomical Registration Through Exponentiated Lie algebra) template was built independently of the spatial preprocessing in fMRIPrep. Specifically, the reoriented T1-weighted image of each participant was segmented into 6 tissue types (i.e., white matter, gray matter, cerebrospinal fluid, bone, soft tissue, background) and an average sample-based template was created and registered to the MNI space using DARTEL (n = 45, 22 children, 23 adults). Each subject’s fMRIPrep-processed T1w image and functional images were then normalized to this template. Additionally, functional images were spatially smoothed using an isotropic 8mm full-width at half-maximum (FWHM) Gaussian kernel.

Activation analyses of task-based fMRI data were performed to assess age group differences in the activation of task-relevant regions during initial learning as well as changes in activity across the macro-offline consolidation interval. These analyses were conducted in two serial steps, accounting for intraindividual (fixed effects) and interindividual (random effects) variance. At the individual level, a general linear model (GLM) was built, estimating changes in regional brain signals in response to practice of the FTT and its linear modulation by performance speed (i.e., median transition time per block). The 20-second rest blocks interspersed between blocks of FTT practice served as the baseline condition, modeled implicitly in the block design. The regressor of interest (i.e., task practice) consists of box cars convolved with the canonical hemodynamic response function. Head motion estimates (i.e., translation and rotation in the three motion planes; see Appendix 4 for head motion values per group) output by spatial pre-processing and erroneous keypresses were included in the model as regressors of no interest in order to maximize the explained variance, and high-pass filtering with a cutoff period of 128s was implemented to remove slow drifts from the time series. Serial correlations in the fMRI signal were estimated using an autoregressive plus white noise model and a restricted maximum likelihood (ReML) algorithm.

Linear contrasts were generated at the individual level to test for (1) the effect of task practice (i.e., task-related activity) within each task run (i.e., training 1, training 2 and retest); (2) the linear modulation of task-related activity by performance speed within each task run (data not presented here); and, (3) the change of task-related activity between pairs of runs. For all linear contrasts, the resulting contrast images were further spatially smoothed using an isotropic 6mm FWHM Gaussian kernel and entered into a second-level analysis for statistical interference at the group level (i.e., independent samples t-test comparing children and adults). Such a group-level analysis corresponds to a random effects model that accounts for inter-subject variance. The set of voxel values resulting from each second-level analysis constitutes maps of the t-statistics [SPM(T)], thresholded at p < 0.001* (uncorrected for multiple comparisons). Significant brain clusters were visually inspected and were excluded when deemed to be fully located in white matter.

For results presented in the main text, statistical inferences were performed at a threshold of p < 0.05 after family-wise error (FWE) correction for multiple comparisons over small spherical volumes (small volume correction (SVC) approach; Poldrack, 2007; Poldrack et al., 2008). Specifically, for clusters within a priori-defined regions of interest (i.e., frontal, motor and parietal cortices, basal ganglia, thalamus, cerebellum and hippocampus, based on the AAL3 atlas of Rolls et al. 2020) that were significant at p(uncorrected)<0.001, coordinates in close proximity to the activation peak were extracted from published motor sequence learning and memory consolidation research (see Appendix 1). Spheres (10 mm radius) were centered on these coordinates and correction for multiple comparisons was applied across voxels included in each spherical volume. Note that activations in occipital and temporal regions were ignored, as these regions are thought to not play a significant role in motor sequence learning and memory consolidation processes. All results reported and discussed in the main text survived SVC. An additional correction for multiple volumes of interest within each contrast was performed using Holm-Bonferroni procedures (p < 0.05; Holm, 1979). Results that survived this additional Holm-Bonferroni correction are simply indicated with an asterisk in the corresponding table.

To assess the link between brain activation and behavioral performance, the individual’s contrast images from the activation-based analyses outlined above were regressed against the individual’s behavioral performance measures. Specifically, we assessed the relationship between (1) task-related brain activity during training 1 and the respective micro-online and -offline performance changes; (2) task-related brain activity during training 2 and the respective micro-online and - offline performance changes; (3) task-related brain activity during training 1 and the macro-offline performance changes; (4) changes in brain activity from training 2 to the retest session (i.e., across the 5-hour offline period) and the macro-offline performance changes. Additionally, an exploratory analysis was conducted with the average (non-normalized) transition time in training 1 as a covariate of no interest. This allowed the determination of whether age group differences in task-related brain activity during training 1 remained after accounting for differences in performance speed. Results are provided in Appendix 5.

The analyses outlined above compared differences between the two age *groups*. To provide more fine-grained analyses of the development of task-induced activity, we also performed exploratory analyses assessing age-related changes in task-induced activity and changes in brain activity from initial training to the retest across childhood. Specifically, the previously described univariate fMRI analyses were run with age as a covariate in a regression analysis. Results of these analyses can be found in Appendix 5.

## 3. Results

### 3.1 Participant characteristics, sleep and vigilance

Table 1 provides group means of participant characteristics, self-reported sleep quality and duration for the night preceding the experimental day, subjective (SSS) and objective (PVT) measures of vigilance at time of FTT completion, as well as details on the timing of the experimental protocol. Results from the corresponding statistical analyses can be found in Appendix 2. In brief, significant age group differences were observed for sleep duration and performance on the psychomotor vigilance task (PVT). Specifically, children reported a longer sleep duration than adults for the night prior to the training session, reflecting the known decrease in total sleep time with age (Ohayon et al., 2004). Additionally, children performed slower on the PVT as compared to adults, which is in line with prior research (Iida et al., 2010). The main effect of session and the group by session interaction on the PVT data were not significant. This demonstrates that vigilance remained similar across the two sessions for the two groups and suggests that any observed behavioral and imaging results reported below cannot be attributed to changes in vigilance across the testing day.

### 3.2 Behavioral results

The full sample (n = 22 children, 23 adults) was included in the behavioral analyses. Two children were excluded from statistical contrasts including training 2 (see Appendix 1 for details).

#### 3.2.1 Learning dynamics

Normalized performance measures from all FTT runs are depicted in Figure 2 and the full statistical output can be found in Appendix 3. Here, we provide a review of the key results with a focus on normalized performance speed, as performance accuracy remained relatively stable across practice blocks and largely comparable between age groups (see Figure 2B). Normalized transition time across training runs showed a non-significant trend towards a group x training run x block interaction effect (F_(5.27,221.14)_ = 2.14, p = 0.058, ƞ^2^ = 0.009, BF_10_ = 0.020). Follow-up ANOVAs within each training run indicated significant main effects of group (run 1: F_(1,43)_ = 8.85, p = 0.005, ƞ^2^ = 0.107, BF_10_ = 6.661; run 2: F_(1,42)_ = 4.10, p = 0.049, ƞ^2^ = 0.064, BF_10_ = 0.929) and block in both runs (run 1: F_(5.80, 249.41)_ = 26.08, p < 0.001, ƞ^2^ = 0.202, BF_10_ = 1.414; run 2: F_(4.27,179.43)_ = 3.43, p = 0.008, ƞ^2^ = 0.024, BF_10_ = 8685.588). There was no significant group x block interaction effect in either run (both p > 0.14; see Appendix 3 for details). These results indicate that performance speed improved across practice blocks within each training run, with slower performance in children but similar performance changes across blocks in the two age groups.

**Figure 2.**
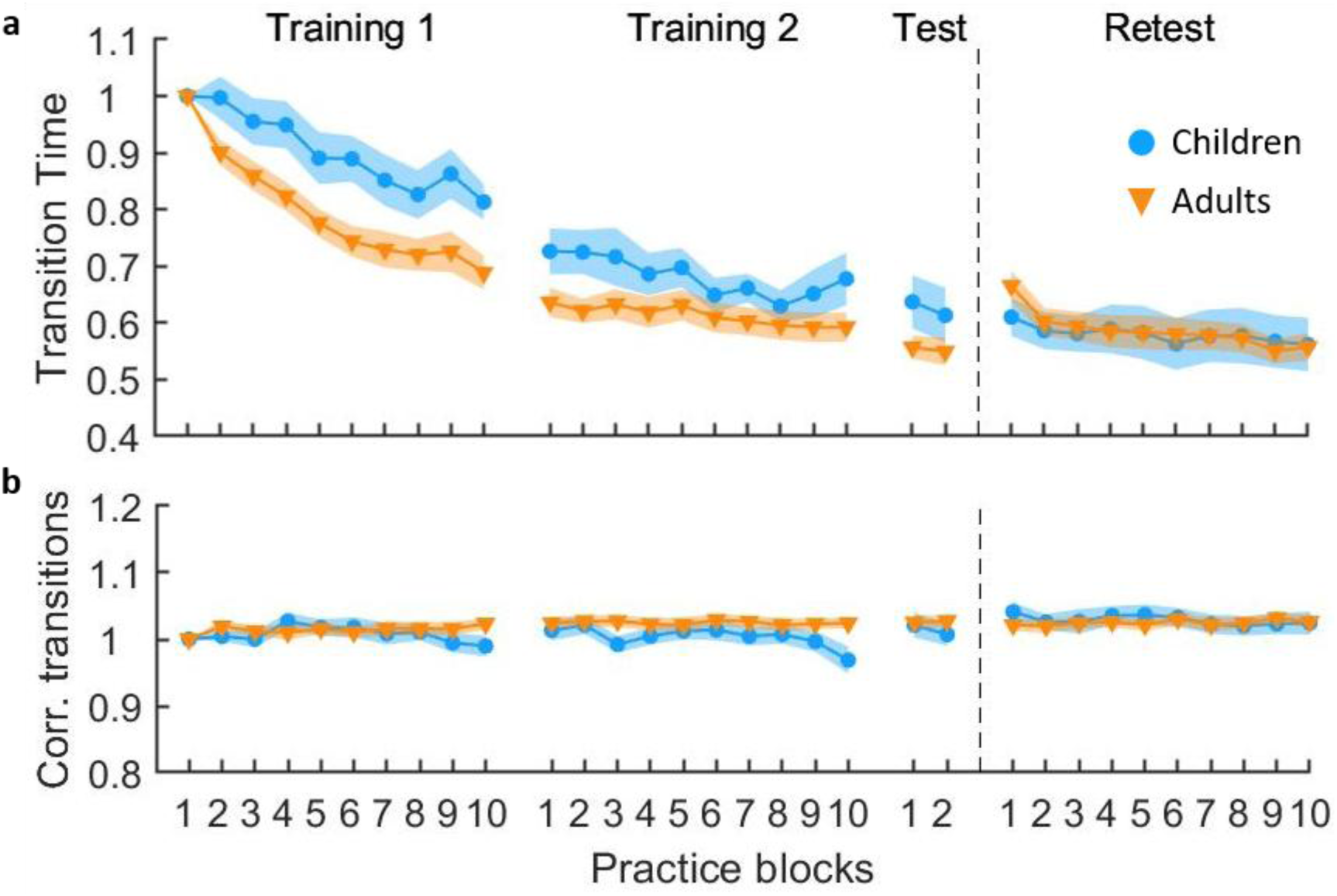
Normalized motor performance across FTT task runs. Measures of movement speed (median normalized transition time; panel **a**) and accuracy (normalized percentage of correct transitions; panel **b**) for the training, test and retest runs. Shaded regions represent standard errors of the mean. n = 22 for children and n = 23 for adults for all task runs, except training 2 (n = 21 children). The main text provides primary statistical information of interest but see Appendix 3 for complete statistical output.

During the post-learning test, normalized speed showed no credible evidence for effects of group, block or a group x block interaction effect (all p > 0.13; full output provided in Appendix 3). These results indicate that children and adults showed comparable performance at the end of training and prior to assessing macro-offline consolidation.

During the retest, normalized performance speed showed a significant main effect of block (F_(2.48,106.72)_ = 4.91, p = 0.005, ƞ^2^ = 0.017, BF_10_ = 5171.154), but no group (F_(1,43)_ = 0.03, p = 0.873, ƞ^2^ < 0.001, BF_10_ = 0.493) or group x block interaction (F_(2.48,106.72)_ = 0.97, p = 0.398, ƞ^2^ = 0.003, BF_10_ = 0.037). These results collectively suggest that children and adults demonstrated comparable improvements in performance speed as a function of practice during the retest.

#### 3.2.2 Micro-learning across training

Micro-online and -offline performance changes are depicted in Figure 3. Panels a and b show the cumulative sums of these performance changes as a function of practice blocks (i.e., values of block n show the sums of the gain values from block 1 through block n). Age group by training run ANOVAs were conducted on the average micro-online and -offline performance changes across task blocks or rest intervals, respectively (Panels c and d in Figure 3). For micro-online performance changes, the group x training run interaction was not significant (F_(1,43)_ < 0.01, p = 0.971, ƞ^2^ < 0.001, BF_10_ = 0.287). There was, however, a significant main effect of group, with greater overall micro-online performance changes for adults as compared to children (F_(1,43)_ = 4.46, p = 0.040, ƞ^2^ = 0.058, BF_10_ = 1.416). Additionally, training run 2 demonstrated greater (i.e., less negative) micro-online performance changes than training run 1 (F_(1,43)_ = 4.80, p = 0.034, ƞ^2^ = 0.044, BF_10_ = 2.039). The assessment of micro-offline learning showed no group x training run interaction (F_(1,43)_ = 0.00, p = 0.963, ƞ^2^ < 0.001, BF_10_ = 0.300). While the magnitudes of micro-offline performance changes were larger in children as compared to adults, this effect was a non-significant trend (F_(1,43)_ = 3.70, p = 0.061, ƞ^2^ = 0.044, BF_10_ = 0.869). A main effect of training run was present, with micro-offline performance changes being larger in training run 1 as compared to training run 2 (F_(1,43)_ = 7.99, p = 0.007, ƞ^2^ = 0.080, BF_10_ = 12.154). Although children showed quantitatively larger and smaller micro-online and micro-offline gains, respectively, these results should be interpreted with caution. Between group differences in micro-offline learning did not reach statistical significance (i.e., p = 0.061) and the reported Bayes Factors for age group differences in both micro-online and -offline performance changes can be considered small (i.e., < 1.5).

**Figure 3.**
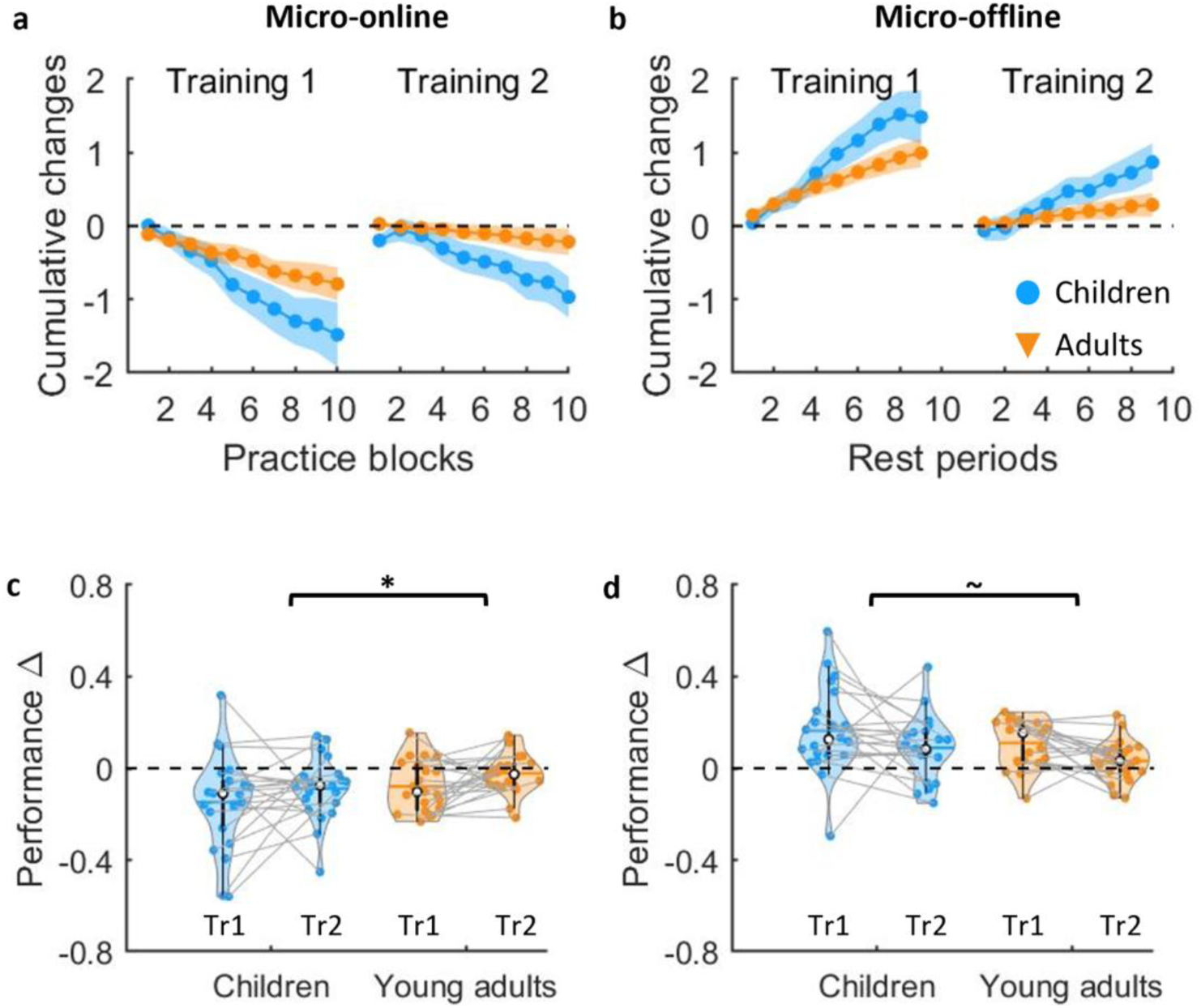
Micro-online (**left**) and micro-offline (**right**) performance changes. The top panels depict the cumulative sums of the micro-online (**a**) and micro-offline (**b**) performance changes across practice/rest epochs for children and adults. Shaded regions represent standard errors of the mean. The bottom panels contain violin plots of micro-online (**c**) and -offline (**d**) gains averaged across blocks for Training Runs 1 and 2 (Tr1 and Tr2, respectively). Positive values are indicative of performance improvements. Shaded regions represent the kernel density estimates of the data, colored circles depict individual data, open circles represent group median, and the horizontal lines depict group means (Bechtold et al., 2021). * p < 0.05 and ∼ p < 0.1 for group differences across the two training runs. n = 22 during training 1 and n = 21 during training 2 for children. n = 23 for adults.

#### 3.2.2 Macro-offline performance changes

Age group differences in macro-offline performance changes are depicted in Figure 4. Analyses revealed that children, as compared to adults, exhibited larger macro-offline changes in both performance speed (t_43_ = 2.059, p = 0.023, G = 0.60, BF_10_ = 1.573) and accuracy (t_28.685_ = 2.527, p = 0.009, G = 0.74, BF_10_ = 3.812). For completeness, we conducted an exploratory analysis on the magnitude of offline gains using an often-used dependent measure (Dan et al. 2015; King et al. 2016) that combines speed and accuracy measures into a single metric (global performance index – GPI). Children demonstrated greater offline gains in GPI as compared to adults (depicted Appendix 3; t_43_ = 2.583, p = 0.007, G = 0.76, BF_10_ = 4.233). Collectively, these results demonstrate that children exhibited greater macro-offline performance changes than adults, a finding that is in line with our previous work (Van Roy et al., 2024).

**Figure 4.**
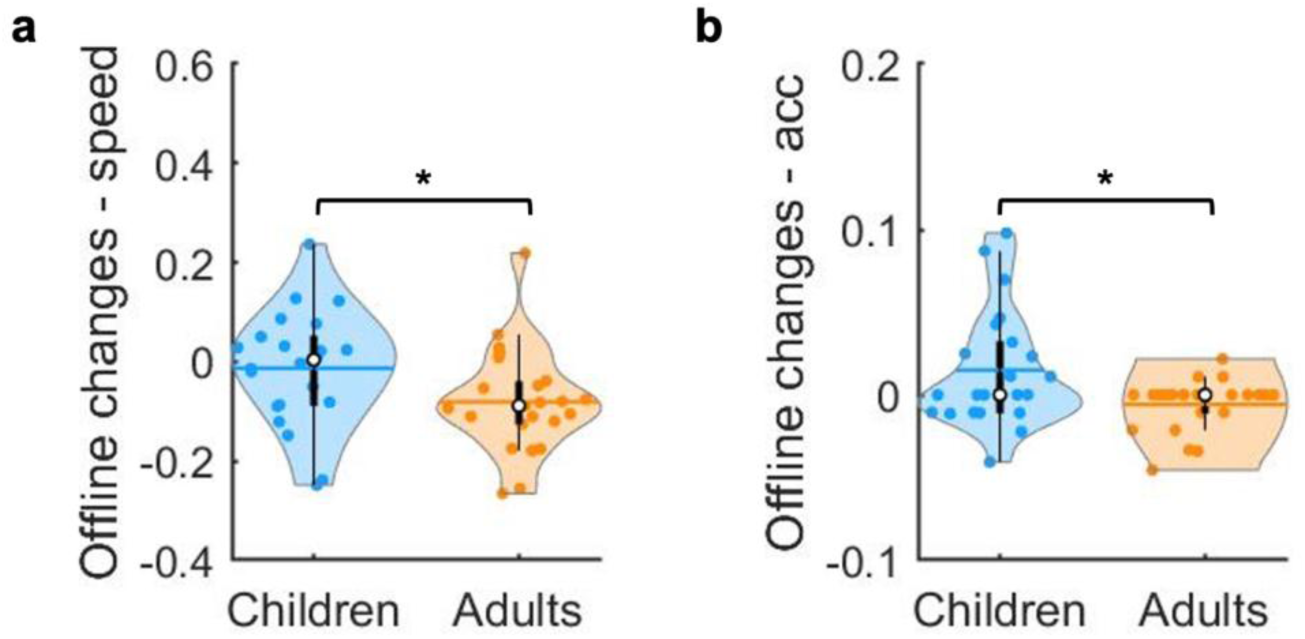
Macro-offline performance changes. Offline changes in normalized speed (**panel a**) and accuracy (**panel b**) across the 5-hour offline period for children and adults. Positive values are indicative of performance improvements from end of training to the 5-hour delayed retest (i.e., larger values reflect enhanced offline consolidation). Shaded regions represent the kernel density estimates of the data, colored circles depict individual data, open circles represent group medians, and the horizontal lines depict group means (Bechtold et al., 2021). * p < 0.05 for group differences. n = 22 for children and n = 23 for adults.

### 3.3 Functional neuroimaging results

The main text focuses on differences between age groups in brain activation during initial learning (i.e., training 1) and changes in activity from training 2 to retest (i.e., across the macro-offline period), as well as their relationship with behavioral performance measures. Results pertaining to brain activity during training 2 and retest, and activity changes from training 1 to training 2 are presented in Appendix 5.

#### 3.3.1 Brain activity during early learning

Age group differences in the recruitment of brain regions during initial learning (i.e., training 1) were assessed by comparing children and adults in the main effect of task practice (interleaved rest as the implicit baseline; see Figure 5; corresponding results presented in Table 2a). Results revealed that adults exhibited greater activity in a large cluster with an activation peak in the right superior parietal cortex. This cluster extended into the primary motor cortex (M1), postcentral gyrus, supplementary motor area, and cerebellum. Additionally, adults also demonstrated greater activity in a cluster with an activation peak in the right thalamus that extended into the putamen and globus pallidus. Visual inspection of the extracted beta estimates (Figure 5a) as well as the results from the within-group, follow-up analyses (see Appendix 5) revealed that the primary motor cortex, postcentral gyrus, supplementary motor area and cerebellum exhibited significant activations in both age groups (although greater in magnitude in adults). The superior parietal gyrus, thalamus, putamen and globus pallidus showed significant task activity in adults, whereas extracted beta estimates approximated 0 in children (i.e., no differences between task and rest epochs). And lastly, the parahippocampus was significantly deactivated (rest > task) in both age groups, with greater deactivation in children relative as compared to young adults.

**Figure 5.**
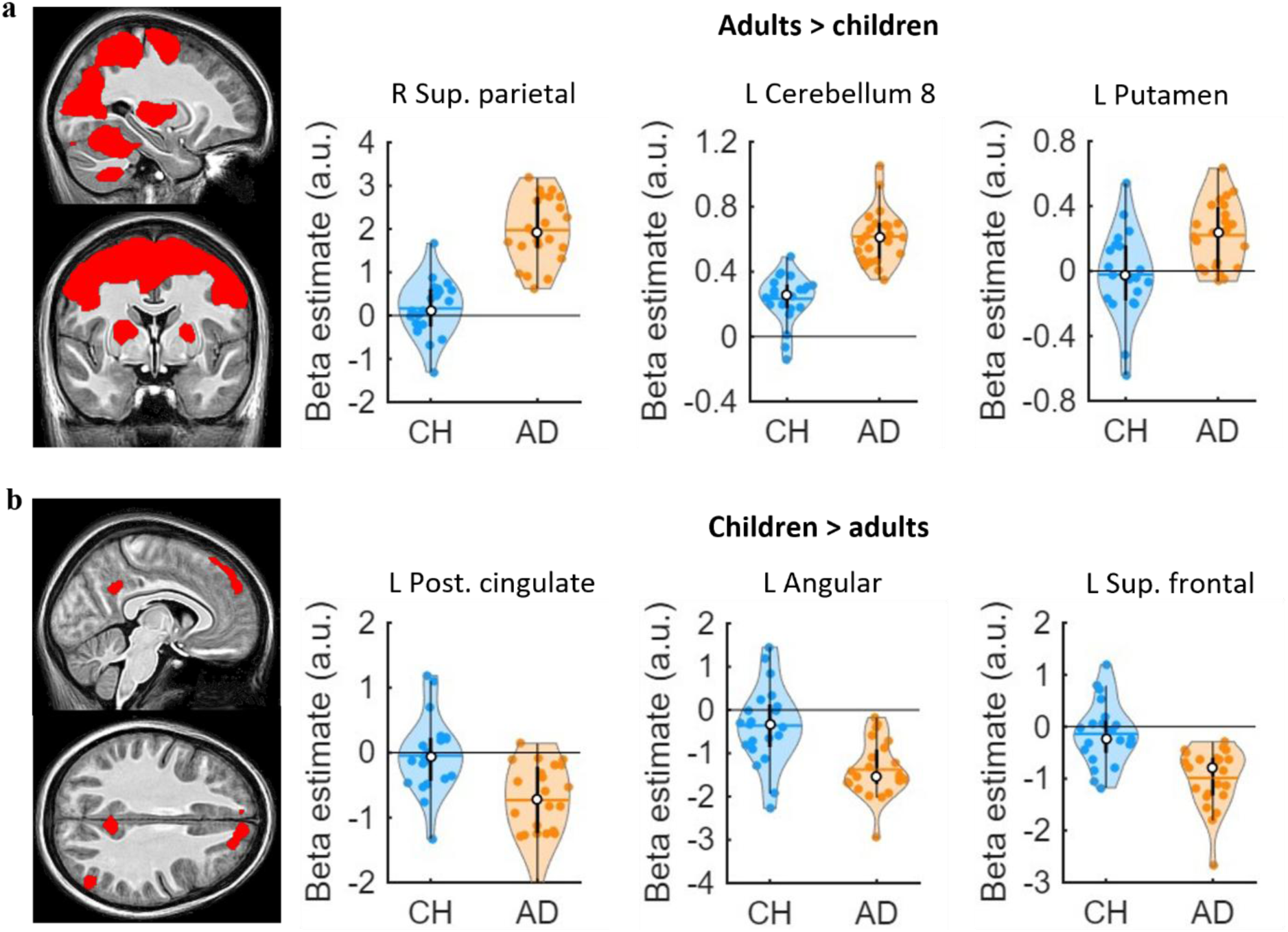
Brain regions showing a significant difference in task-related brain activity between children and adults. **(a)** Adults (AD) > children (CH) contrast. Left images are centered on the L putamen [-26 -1 6]. R sup. parietal = [18 -62 55], L cerebellum 8 = [-20 -60 -48]. **(b)** CH > AD contrast. Left images are centered on the L posterior cingulate peak [-5 50 32]. L angular = [-50 - 68 32], L sup. frontal = [-20 42 40]. Statistical images are thresholded at p_uncorrected_ < 0.001. x, y and z coordinates are specified in MNI space. Violin plots depict extracted parameter (beta) estimates for selected regions of interest. Shaded regions represent the kernel density estimates of the data, colored circles depict individual data, open circles represent group medians, and the horizontal lines depict group means (Bechtold et al., 2021).

**Table 2.** Results of task-related activity during early learning (i.e., Training 1).

| Area | x | y | z | T | P<sub>FWE-SVC</sub> |
| --- | --- | --- | --- | --- | --- |
| **a. [Training 1]** |  |  |  |  |  |
| <u>a1. Children - adults</u> |  |  |  |  |  |
| L Superior frontal | -20 | 42 | 40 | 4.47 | 0.001* |
|  | -12 | 32 | 55 | 4.1 | 0.003* |
| L Angular | -50 | -68 | 32 | 4.04 | 0.003* |
|  | -45 | -75 | 40 | 3.64 | 0.008* |
| L Posterior cingulate | -5 | -50 | 32 | 3.84 | 0.005* |
| L Middle frontal | -38 | 20 | 50 | 3.73 | 0.007* |
| <u>a2. Adults - children</u> |  |  |  |  |  |
| R Superior parietal | 18 | -62 | 55 | 8.29 | < 0.001* |
| R Cerebellum 6 | 22 | -55 | -20 | 8.02 | < 0.001* |
| L Cerebellum 8 | -20 | -60 | -48 | 7.96 | < 0.001* |
| R Thalamus | 22 | -18 | -2 | 6.30 | < 0.001* |
| R Parahippocampus | 32 | -20 | -25 | 4.32 | 0.001* |
| **b. [Training 1 x macro-offline performance changes]** |  |  |  |  |  |
| <u>b1. Children - adults</u> |  |  |  |  |  |
| R Middle frontal | 52 | 45 | 5 | 6.17 | < 0.001* |
|  | 48 | 52 | 8 | 5.32 | < 0.001* |
| L Parahippocampus | -18 | -30 | -18 | 5.05 | < 0.001* |
| L Postcentral | -25 | -38 | 58 | 5.20 | < 0.001* |
| Extends into R hippocampus |  |  |  |  |  |
| R Cerebellum 9 | 15 | -42 | -42 | 4.42 | 0.001* |
| L Precentral | -50 | 5 | 38 | 3.84 | 0.006 |
| L Middle frontal | -50 | 12 | 45 | 3.60 | 0.010 |
| L Inferior frontal, triangular | -55 | 18 | 25 | 3.63 | 0.010 |
| L Amygdala | -25 | 0 | -18 | 3.71 | 0.008 |
| R Supramarginal | 52 | -35 | 32 | 3.61 | 0.010 |
|  | 55 | -42 | 38 | 3.48 | 0.014 |
| R Superior frontal | 30 | 52 | 20 | 3.61 | 0.010 |
|  | 25 | 48 | 30 | 3.46 | 0.014 |
| L Supplementary motor area | -5 | 20 | 55 | 3.45 | 0.015 |
| R Cerebellum crus 1 | 42 | -68 | -35 | 3.53 | 0.012 |
| <u>b2. Adults - children</u> |  |  |  |  |  |
| No suprathreshold clusters |  |  |  |  |  |

Table presents clusters at $p_{FWE-SVC} < 0.05$ after family-wise error (FWE), small volume correction (SVC, 10 mm radius) for multiple comparisons. Voxels that were located outside of grey matter and that did not survive SVC are not reported. \* $p < 0.05$ , Holm-Bonferroni corrected. Minimum cluster size = 5 voxels. x, y and z coordinates are specified in MNI space. Results presented in this table are based on $n = 21$ children and 23 adults. Results of the within- and across-group contrasts are provided in Appendix 5 for completeness.

In contrast to the results described above, children exhibited greater activity as compared to adults in the left middle and superior, left angular, and left posterior cingulate gyri (Figure 5B). Visual inspection of the beta estimates indicated that these regions were deactivated in adults (i.e., more activity during the rest as compared to task periods), whereas children showed parameter estimates approximating zero.

We then conducted regression analyses assessing the link between brain activation during early training (i.e., training 1; see Appendix 5 for regressions with training 2 activity) and the behavioral performance measures, and whether this differed between age groups. Results revealed no significant age group differences in the relationship between activity during early learning and *micro*-online and -offline performance changes (see Appendix 5).

For *macro*-offline performance changes, the regression analyses showed significant group differences in a large cluster with an activation peak in the left parahippocampus (Figure 6, Table 2b). This cluster extended into the hippocampus, thalamus, caudate nucleus, bilateral primary motor cortices, postcentral gyrus, right supplementary motor area, bilateral middle and posterior cingulate gyri and paracentral lobule. Additionally, group differences were observed in the middle frontal gyrus, left inferior and right superior frontal gyri, left M1, right supramarginal gyrus, left amygdala, and cerebellum. The extracted parameter (beta) estimates revealed that task-related activity in most of these regions was positively related to macro-offline performance changes *within* children, whereas adults either showed the opposite pattern (i.e., negative relationship) or no association.

**Figure 6.**
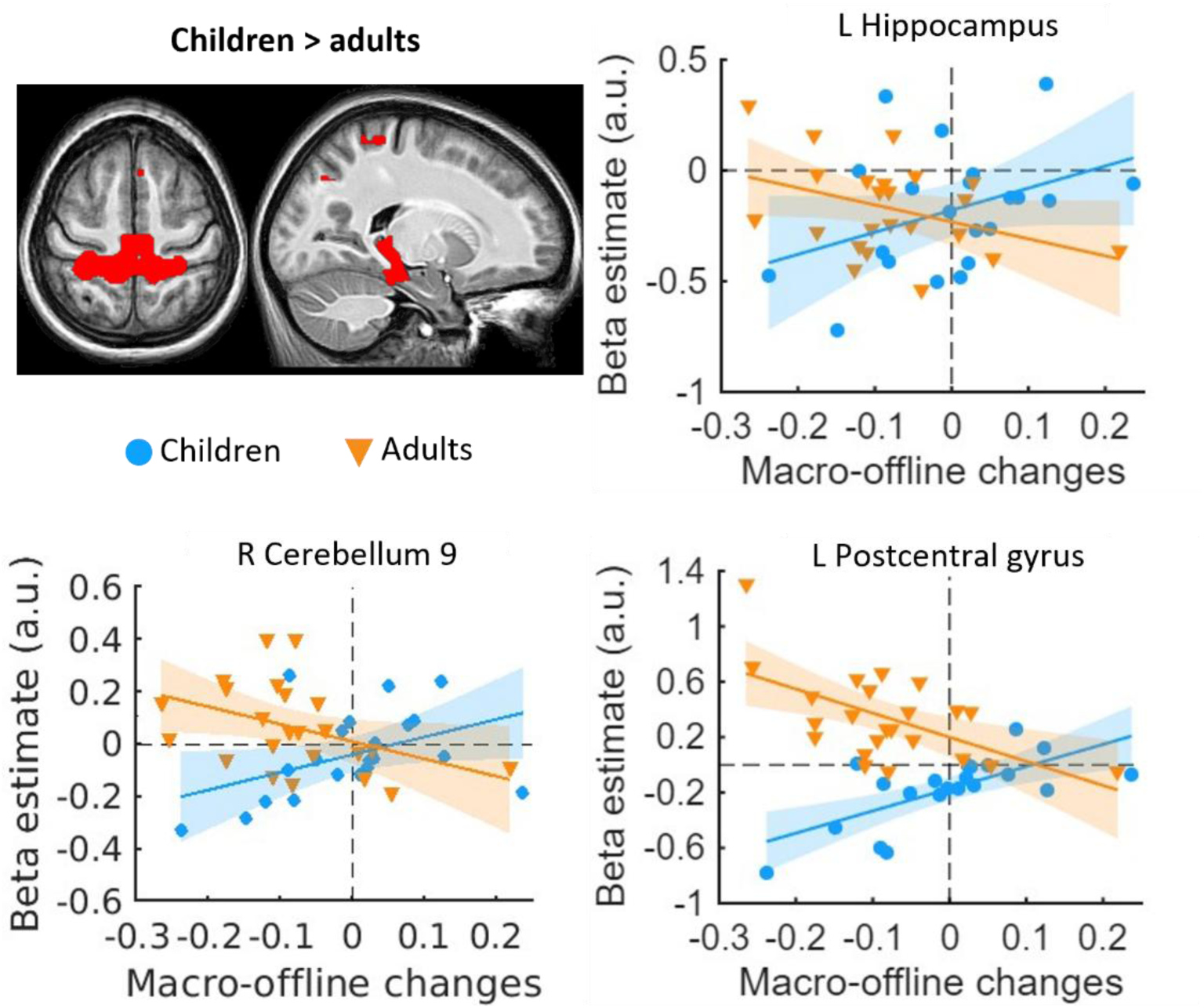
Brain regions showing significant age group differences in the relationship between task-related brain activity during training 1 and macro-offline performance changes. Children > adults contrast. Left brain image is centered on the L postcentral peak [ -25 -38 58]. Right brain image is centered on the L hippocampus [ -19 -27 -11]. R cerebellum 9 = [15 -42 -42]. Statistical images are thresholded at p_uncorrected_ < 0.001. Extracted parameter (beta) estimates for selected regions of interest are plotted as a function of macro-offline performance changes. x, y and z coordinates are specified in MNI space. Shaded regions represent the 95% confidence intervals.

Altogether, results concerning training 1 revealed that children exhibited less activation as compared to adults across a widespread set of brain regions that included cortical (i.e., superior parietal cortex, M1, postcentral gyrus, SMA) and subcortical (i.e., thalamus, putamen and globus pallidus) motor regions as well as the cerebellum. Furthermore, activations during early learning in the dorsolateral prefrontal cortex, sensorimotor cortex, supramarginal gyrus, cingulate gyrus, amygdala, caudate, (para)hippocampus, thalamus and cerebellum were positively related to macro-offline performance improvements in children.

#### 3.3.2 Changes in brain activity from training 2 to retest

As outlined in our registration, changes in brain activity across the 5-hour macro-offline period were examined by contrasting training 2 and retest (Figure 7, Table 3a). Results revealed significant group differences in inter-session changes in activity in the bilateral cerebellum (i.e., left cerebellum crus I, bilateral cerebellum 4-5, right cerebellum 9), right M1 and left hippocampus. Inspection of the beta estimates indicated that these group by session interactions were largely due to a decrease in activity in the left hippocampus in children from training 2 to retest, whereas adults demonstrated inter-session increases in the right M1 and cerebellum. Children also exhibited a slight decrease in activity in cerebellum crus I.

**Figure 7.**
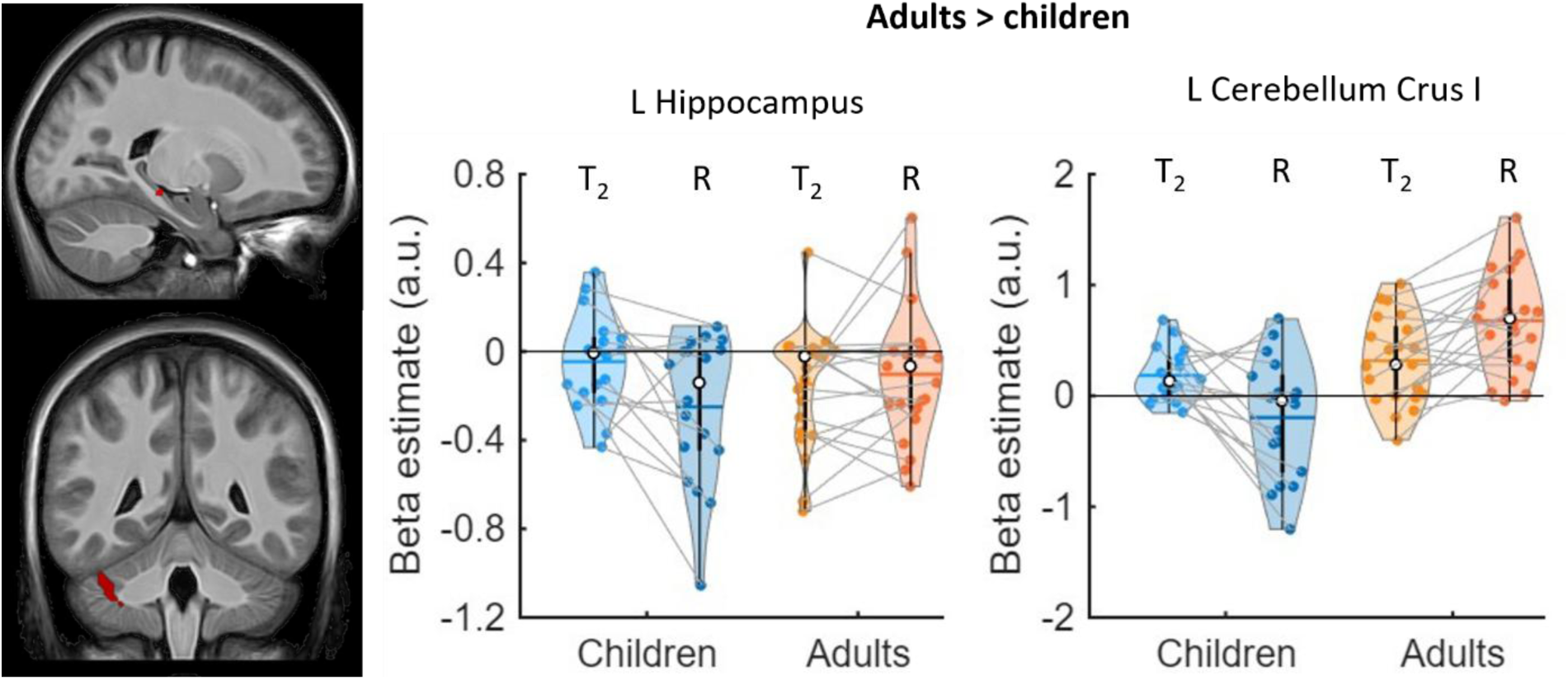
Brain regions showing significant age group differences in the inter-session changes in activity from training 2 to retest. Adults > children contrast. Left images are centered on the L hippocampus peak [-20 -28 -12] (top) and L cerebellum crus I peak [-40 -45 -35] (bottom). Statistical images are thresholded at p_uncorrected_ < 0.001. x, y and z coordinates are specified in MNI space. Violin plots depict extracted parameter (beta) estimates for selected regions of interest. Shaded regions represent the kernel density estimates of the data, colored circles depict individual data, open circles represent group medians, and the horizontal lines depict group means (Bechtold et al., 2021). T_2_ = Training 2. R = Retest.

**Table 3.**
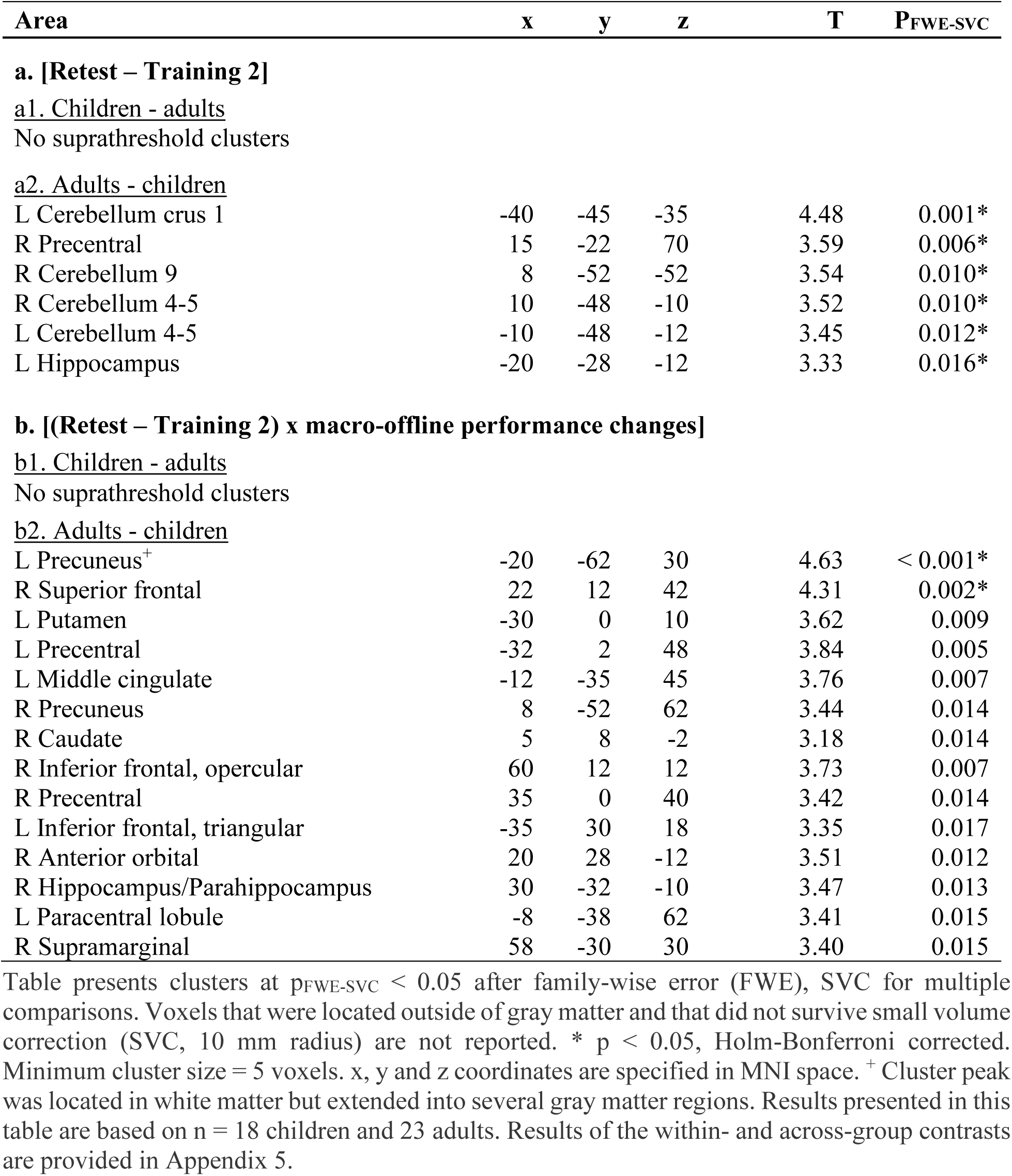
Changes in task activity from training 2 to retest [Retest – Training 2].

| Area | x | y | z | T | P<sub>FWE-SVC</sub> |
| --- | --- | --- | --- | --- | --- |
| **a. [Retest – Training 2]** |  |  |  |  |  |
| <u>a1. Children - adults</u> |  |  |  |  |  |
| No suprathreshold clusters |  |  |  |  |  |
| <u>a2. Adults - children</u> |  |  |  |  |  |
| L Cerebellum crus 1 | -40 | -45 | -35 | 4.48 | 0.001* |
| R Precentral | 15 | -22 | 70 | 3.59 | 0.006* |
| R Cerebellum 9 | 8 | -52 | -52 | 3.54 | 0.010* |
| R Cerebellum 4-5 | 10 | -48 | -10 | 3.52 | 0.010* |
| L Cerebellum 4-5 | -10 | -48 | -12 | 3.45 | 0.012* |
| L Hippocampus | -20 | -28 | -12 | 3.33 | 0.016* |
| **b. [(Retest – Training 2) x macro-offline performance changes]** |  |  |  |  |  |
| <u>b1. Children - adults</u> |  |  |  |  |  |
| No suprathreshold clusters |  |  |  |  |  |
| <u>b2. Adults - children</u> |  |  |  |  |  |
| L Precuneus<sup>+</sup> | -20 | -62 | 30 | 4.63 | < 0.001* |
| R Superior frontal | 22 | 12 | 42 | 4.31 | 0.002* |
| L Putamen | -30 | 0 | 10 | 3.62 | 0.009 |
| L Precentral | -32 | 2 | 48 | 3.84 | 0.005 |
| L Middle cingulate | -12 | -35 | 45 | 3.76 | 0.007 |
| R Precuneus | 8 | -52 | 62 | 3.44 | 0.014 |
| R Caudate | 5 | 8 | -2 | 3.18 | 0.014 |
| R Inferior frontal, opercular | 60 | 12 | 12 | 3.73 | 0.007 |
| R Precentral | 35 | 0 | 40 | 3.42 | 0.014 |
| L Inferior frontal, triangular | -35 | 30 | 18 | 3.35 | 0.017 |
| R Anterior orbital | 20 | 28 | -12 | 3.51 | 0.012 |
| R Hippocampus/Parahippocampus | 30 | -32 | -10 | 3.47 | 0.013 |
| L Paracentral lobule | -8 | -38 | 62 | 3.41 | 0.015 |
| R Supramarginal | 58 | -30 | 30 | 3.40 | 0.015 |

Table presents clusters at $p_{\text{FWE-SVC}} < 0.05$ after family-wise error (FWE), SVC for multiple comparisons. Voxels that were located outside of gray matter and that did not survive small volume correction (SVC, 10 mm radius) are not reported. \* $p < 0.05$ , Holm-Bonferroni corrected. Minimum cluster size = 5 voxels. x, y and z coordinates are specified in MNI space. <sup>+</sup> Cluster peak was located in white matter but extended into several gray matter regions. Results presented in this table are based on $n = 18$ children and 23 adults. Results of the within- and across-group contrasts are provided in Appendix 5.

We then assessed whether changes in brain responses from training 2 to the 5-hour retest were related to the macro-offline performance changes. Results revealed that inter-session changes in activity in the bilateral inferior and right superior frontal gyri, left middle cingulate gyrus, bilateral M1, left paracentral lobule, right supramarginal gyrus, bilateral precuneus, left putamen, right caudate nucleus, and right hippocampus/parahippocampus were differentially related to the macro-offline performance changes between the two groups (Figure 8, Table 3b). Inspection of the beta estimates was conducted to decompose these effects and revealed that greater decreases in activity from training 2 to retest in the left inferior and right superior frontal gyrus, left middle cingulate gyrus, left paracentral lobule, precuneus, left putamen, right caudate and right hippocampus were related to greater macro-offline performance changes *within* children, whereas no relationships were observed within adults.

**Figure 8.**
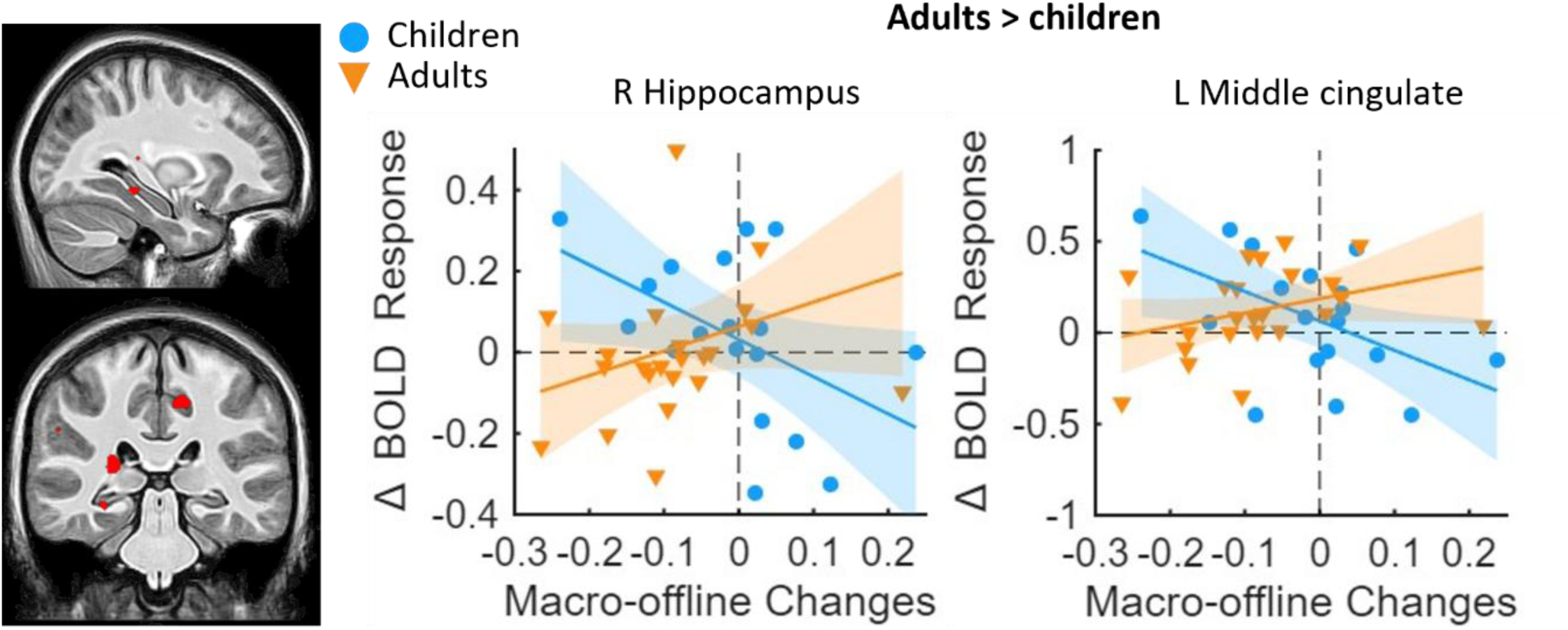
Brain regions showing significant age group differences in the relationship between inter-session changes (retest – training 2) in activity and macro-offline performance changes. Adults > children contrast. Left image is centered on the R hippocampus peak [30 -32 -10]. L middle cingulate = [-12 -35 45]. Statistical images are thresholded at p_uncorrected_ < 0.001. Extracted parameter (beta) estimates that represent inter-session changes in activity for selected regions of interest are plotted as a function of macro-offline performance changes. x, y and z coordinates are specified in MNI space. Shaded regions represent the 95% confidence intervals.

Taken together, children exhibited decreases in activity in the left hippocampus from training 2 to retest on the motor sequence, whereas adults showed inter-session increases in the right M1 and cerebellum. Moreover, greater inter-session decreases in activity in prefrontal cortex, hippocampus and the putamen were associated with larger macro-offline performance changes in children.

## 4. Discussion

The current study employed task-related fMRI to identify the neural correlates of MSL in children, including the developmental advantage in wake-related offline motor memory consolidation. The behavioral results reproduced our previously shown childhood benefit in awake macro-offline motor memory consolidation (Van Roy et al. 2024). Neuroimaging results revealed that: (1) children exhibited less activation as compared to adults during early training across a widespread set of brain regions that included cortical (i.e., superior parietal cortex, M1, postcentral gyrus, SMA) and subcortical (i.e., thalamus, putamen and globus pallidus) regions as well as the cerebellum; (2) there was a positive relationship between macro-offline performance improvements in children and activity during early learning in the dorsolateral prefrontal cortex, sensorimotor cortex, supramarginal gyrus, cingulate gyrus, amygdala, caudate, (para)hippocampus, thalamus and cerebellum; (3) children exhibited decreases in activity in the hippocampus from late initial learning to the retest, whereas adults showed inter-session increases in the M1 and cerebellum; and, (4) greater inter-session decreases in activity in prefrontal cortex, hippocampus and the putamen were associated with larger macro-offline performance changes in children.

The discussion below focuses on the interpretation of the primary differences between children and adults. Appendix 6 includes supplementary discussions on the behavioral results assessing initial learning dynamics (section 6.1), micro-offline performance changes (section 6.2), the relationship between brain activity during initial learning and micro-offline performance (section 6.4), and the inter-session changes in brain activity in adults (section 6.5).

### 4.1 Children exhibit a behavioral advantage in awake memory consolidation

Behavioral results of the current research revealed that macro-offline consolidation was enhanced in children, as evidenced by their larger performance changes in both speed and accuracy across the 5-hour macro-offline period. While adults exhibited a deterioration in performance (i.e., offline performance changes were significantly smaller than zero; see Appendix 3), children maintained their performance level over the 5-hour interval. This childhood advantage is consistent with the larger 5-hour performance changes in speed that we previously observed in children (Van Roy et al., 2024) as well as the enhanced offline consolidation in children reported by other researchers (Ashtamker & Karni, 2013; Beck et al., 2024; Voisin et al., 2024). Accordingly, our results add to the body of evidence demonstrating that children exhibit superior offline consolidation of acquired motor sequences across a post-learning period of wakefulness.

### 4.2 Children exhibit smaller modulations in brain activity between task and rest than adults

Analyses assessing age group differences during initial training (i.e., training 1) revealed that adults exhibited greater activation as compared to children across a widespread set of brain regions that included cerebellar, cortical and subcortical areas. Follow-up, within-group contrasts and visual inspection of the extracted beta estimates revealed that these between-group differences could be decomposed into different patterns of results. First, both children and adults showed significant activations of cerebellar and motor regions (i.e., superior parietal cortex, M1, postcentral gyrus, SMA) known to be involved in motor task performance, yet the magnitude of these activations was smaller in children. This result is partially consistent with the findings of Thomas et al. (2004), which observed greater cortical activation in adults as compared to children. One could argue that the reduced cortical activation in children may reflect the ongoing development of the cerebral cortex (i.e., cortical thinning and myelination; Kwon et al., 2020; Muftuler et al., 2011; Østby et al., 2009), as prior research in 8- to 18-year-old children has found positive relationships between the maturation of white matter and brain activation in task-relevant regions during a working memory task (Olesen et al., 2003). An alternative explanation is that the reduced activation of these areas during task practice in children may stem from greater engagement during the interleaved rest periods that serve as the baseline condition in the univariate fMRI block design. In other words, the involvement of these regions *during active task practice* may be relatively comparable between the two age groups, but the engagement of these regions *during interleaved rest* is greater in children. These two alternatives cannot be disentangled with the employed analytic approaches.

The second pattern that contributed to the overall greater activation in adults as compared to children was predominantly located in subcortical regions. Specifically, adults showed significant activation of the thalamus, putamen and globus pallidus, whereas children exhibited parameter estimates that approximated zero (i.e., the magnitude of activity was similar in task and rest). The lack of significant activations in children could simply be the result of these regions not being recruited during task practice (nor in interleaved rest). Alternatively, it could be that these regions are indeed recruited, but equivalently across task and rest epochs. Regardless of these competing explanations, these data are not in line with those presented in Thomas et al. (2004), which observed greater activation of subcortical regions in children relative to adults. Similarly, other developmental studies also demonstrated significant recruitment of the putamen and thalamus during a motor sequence learning task in children (Hedenius & Persson, 2022; Sharer et al., 2015).

Analyses of age group differences during early training also revealed group differences in the activation of a subset of brain regions commonly associated with the default mode network (DMN), including the middle and superior frontal, angular, and posterior cingulate gyri. Follow-up, within-group contrasts and visual inspection of the extracted beta estimates indicated that these between-group differences were the result of greater deactivation in adults as compared to children. These results in adults align with the well-established decreased DMN activity during task practice as compared to rest (Raichle, 2015). Furthermore, the differences between children and adults are consistent with previous research that observed developmental increases in brain deactivation during a repetitive button-press task (Morita et al., 2019). As the DMN has been associated with mind-wandering and other inward-directed cognition in adults (Buckner & DiNicola, 2019; Christoff et al., 2009), one could speculate that children disengage from the task less during the short rest periods and thus, show less deactivation of the DMN during task as compared to rest.

Altogether, results suggest that children exhibit smaller modulations in brain activity between task and rest epochs. Based on this finding, we speculate that the child’s brain may stay more engaged during the short rest breaks relative to adults. Several potential explanations could contribute to this continued engagement of the developing brain during interleaved rest, including active consolidation processes (Buch et al., 2021; Gann et al., 2023), the increased need of executive functions to lay stay still in the scanner (Camacho et al., 2020), or cognitive preparation for the upcoming practice block (Camacho et al., 2020). Appendix 6 (section 6.3) provides a more extensive discussion of these possibilities. Although the current experimental design and analytic approach do not allow us to distinguish between these various possible explanations, our results do suggest that brain activity during the interleaved rest epochs – as opposed to solely during task practice - may contribute to the observed age-related differences.

### 4.3 The relationship between functional activity during training 1 and offline performance changes

As the current study aimed to identify the functional neural correlates underlying the childhood advantage in offline processing, we conducted regression analyses that assessed the relationship between brain activation during initial training (i.e., training 1) and the behavioral macro-offline performance changes. Group differences were observed between children and young adults, which were largely driven by a positive relationship between macro-offline performance changes and functional activity during early training in the dorsolateral prefrontal cortex, sensorimotor cortex, supramarginal gyrus, cingulate gyrus, amygdala, caudate, (para)hippocampus, thalamus and cerebellum in children.

The sensorimotor cortex has been shown to be involved in the acquisition of a motor sequence in both pediatric (Hedenius & Persson, 2022; Mostofsky et al., 2006) and adult (Albouy et al., 2015; Dolfen et al., 2021; Nicolas et al., 2025) populations, a result that was replicated in the current study. Visual inspection of the positive brain-behavior relationships revealed that greater activity in these regions during task practice relative to rest was associated with greater macro-offline performance changes.

Activity in the DLPFC, caudate nucleus, (para)hippocampus, supramarginal gyrus and cerebellum crus I were also positively related to the macro-offline performance changes in children. Prior research in adults has demonstrated that the prefrontal and parietal regions as well as the caudate nucleus, (para)hippocampus and cerebellum crus I are recruited during the early phase of initial practice that requires high cognitive control (Albouy et al., 2013; Doyon et al., 2002; Hikosaka et al., 2002). Activity in these regions typically decreases as a function of practice, when the participant becomes more proficient at performing the motor sequence. This suggests that enhanced macro-offline consolidation in children may be linked to higher activation of brain regions that are engaged early in the learning process and under high cognitive control. Moreover, the early engagement of the hippocampus has been thought to play a key role in forming the abstract representation of a motor sequence, which is encoded in spatial coordinates (i.e., organization of the stimuli in space outside of the individual’s body) and is effector-independent (i.e., irrespective of the effector involved in the execution; Albouy et al., 2015; Doyon, Korman, et al., 2009). This hippocampal activity has previously been shown to predict subsequent consolidation across a period of sleep in adults (Albouy et al., 2008). It is thus tempting to speculate that greater macro-offline performance changes in children are associated with the development of an abstract representation of the motor sequence during early training.

Interestingly, visual inspection of the positive relationships in the hippocampus, supramarginal gyrus and DLPFC revealed that smaller and larger macro-offline performance changes were linked to beta estimates that were negative or approximated zero, respectively. Accordingly, significant deactivation of these regions during active task practice was detrimental for offline consolidation in children. Conversely, similar activations during task practice and rest (i.e., beta estimates approximating 0) were beneficial for consolidation. This suggests that similar levels of activity across task and rest epochs - potentially reflecting continued engagement of these regions during interleaved rest - facilitated the childhood advantage in macro-offline motor memory consolidation.

### 4.4 Inter-session changes in neural activity

The assessment of changes in brain activity across the 5-hour offline period (i.e., from training 2 to retest) revealed age group differences in the primary motor cortex, hippocampus and cerebellum (i.e., cerebellum lobule 4-5, lobule 9 and crus I). Specifically, children exhibited significant inter-session decreases in hippocampal activity, whereas adults showed significant increases in activation of M1 and the cerebellum (see Appendix 6 for a discussion of the changes in adults). Moreover, inter-session decreases in activity in the prefrontal cortex, hippocampus and putamen supported larger macro-offline performance changes in children, whereas adults failed to show such relationships.

The opposing inter-session changes in hippocampal activity between age groups, along with the differing relationships between macro-offline consolidation and inter-session changes in hippocampal and striatal activity, may reflect age-related differences in awake offline motor memory consolidation processes. Specifically, children demonstrated greater decreases in hippocampal activity from training 2 to retest as compared to adults. Furthermore, greater inter-session *decreases* in activity in the hippocampus and striatum were linked to larger performance changes across the macro-offline period in children, suggesting that these decreases at least partially support the enhanced consolidation in children. Whereas Hedenius & Persson, (2022) also observed a negative relationship between hippocampal activity and offline performance changes on a sequential task in children, research in adults has observed the opposite pattern (Albouy et al., 2008; Albouy, Sterpenich, et al., 2013; Walker et al., 2005). Specifically, in adults, the hippocampus and striatum have been shown to increase in activity across offline periods that include sleep. These sleep-dependent increases in activity are usually observed in parallel to the emergence of significant offline performance changes from training to retest. Additionally, striatal activity has also been shown to increase across offline periods of wakefulness, which are accompanied by a maintenance of performance (Albouy et al., 2008; Albouy, Sterpenich, et al., 2013). Based on these findings, we hypothesized that – similar to adults – the enhanced macro-offline performance changes in children would be linked to increases in striatal and hippocampal activity from training 2 to retest. Our observation of the opposite pattern in children suggests that the childhood benefit in macro-offline consolidation is supported by inter-session changes in activity that are distinct from those in adults. Further research is needed to clarify the underlying mechanisms contributing to this uniqueness in the developing brain.

### 4.6 Limitations and future directions

The current study has limitations that should be considered. *First*, there was a relatively high, although largely expected, exclusion rate of young children (i.e., 6- and 7-year-olds; see flow diagram for inclusion and exclusion in Appendix 1). As a result, we did not acquire task-related imaging data from any 6-year-olds, and 7-year-olds were relatively under-sampled in our experiment. Future studies could consider adjusting the motor task (e.g., spatially cued task) or implementing certain measures (e.g., additional familiarization in a mock scanner) to increase the number of young children that successfully complete the experiment.

*Second,* head motion can impact neuroimaging results. As children exhibited more head movement during the scanning (see Appendix 4), it is plausible that the deleterious influence of head motion on the BOLD signal is magnified in this age group (Greene et al., 2018). Although we adopted multiple approaches to minimize the negative impact of head motion (e.g., realignment of BOLD images, exclusion of runs where maximum linear movement exceeded 5 mm and inputting head motion parameters as nuisance regressors), we cannot eliminate the possibility that head motion affected the fMRI results more in children than in adults.

*Third*, the task design employed in this research does not allow the assessment of whether performance improvements were specific to sequence learning or general improvements in motor performance (e.g., mapping the numbers to the keys, getting better at pressing the keys). Sequence-specific learning is typically assessed by incorporating random trials in the motor task and comparing performance between sequential and random trials (see Van Roy et al., 2024 for an example). Nevertheless, results of our previous study indicated that children indeed exhibited sequence-specific learning (Van Roy et al., 2024), albeit with a smaller magnitude than in adults. Furthermore, the childhood advantage in macro-offline consolidation reported in this earlier work was shown to be specific to strengthening the sequential memory rather than a general improvement. Accordingly, it is likely that the developmental advantage reported in the current research was also specific to the acquired motor sequence.

*Fourth,* the sample size calculation used in our study registration was based on expected behavioral effect sizes rather than neural outcomes. This choice was largely motivated by the scarcity of pediatric neuroimaging studies that have examined motor learning and memory consolidation processes. Nonetheless, it is worth emphasizing that the magnitudes of task-based fMRI effects may be smaller than behavioral outcomes and thus it is possible that our neuroimaging analyses are under-powered.

*Fifth*, it is worth explicitly stating that only a subset of the brain regions for which activity linked to the magnitudes of offline gains (i.e., regression analyses reported in Tables 2b and 3b) survived the additional step of Holm-Bonferroni correction performed for the number of clusters. Accordingly, results pertaining to the regions that did not survive this correction should be interpreted with extra caution.

*Sixth,* there remain several open questions regarding the interpretation of the observed neural activity in children. Future research should systematically manipulate the experimental task or design to determine the specific learning mechanisms that the developing brain prioritizes and supports. For instance, future studies could differentiate between sequence-specific and general motor performance brain activity by incorporating a random task in which the order of the sequence numbers on the screen changes every trial. Additionally, we speculated that children prioritize the encoding of an abstract, spatial representation of the motor sequence. And experimental design like in Albouy et al. (2015) could be implemented in future research to determine whether the developing brain builds separate spatial and motoric representations of a motor sequence, and then identify the respective underlying neural correlates.

## 5. Conclusions

The present study provided a comprehensive examination of the behavioral and neural correlates of motor sequence learning and memory consolidation in 7- to 11-year-old children. Behavioral results added to the growing body of evidence suggesting that children exhibit enhanced offline consolidation over epochs of post-learning wakefulness. Results from our task-related fMRI analyses revealed that children exhibited less neural activation during initial motor sequence learning – as compared to interleaved rest intervals – across a widespread network of brain regions, including the striatum, cerebellum and cortical motor areas. We speculate that this smaller activation in children may be – at least partially – attributed to greater engagement of these areas during the interleaved rest periods in children as compared to adults. The enhanced performance improvements across the 5-hour offline period in children were associated with task-related activations during initial sequence learning in frontoparietal, cerebellar, sensorimotor and subcortical regions, as well as inter-session decreases in prefrontal, sensorimotor and subcortical activity. Further investigations into the specific learning mechanisms underlying the neural correlates of motor sequence learning and memory consolidation in children are warranted.

## Acknowledgements

This work was supported by internal funds at the University of Utah and a S10 Instrumentation Program Grant awarded to the University of Utah (S10OD026788).

## Notes

### Competing Interest Statement

The authors have declared no competing interest.

### Summary of Updates

Revisions were made as part of the peer review process. There were no major changes in the results.

