## Supplemental material for "Neural correlates of motor sequence learning and enhanced offline consolidation in 7- to 11-year-old children"

**Supplementary information**

**Authors**

Anke Van Roy^1^, Ainsley Temudo^1^, Vincent Koppelmans^2,3^, Kerstin Hödlmoser^4,5^, Genevieve Albouy^1^, Bradley R. King^1*^

**Affiliations**

^1^ Department of Health and Kinesiology, College of Health, University of Utah, Salt Lake City,

UT 84112, USA

^2^ Department of Psychiatry, University of Utah, Salt Lake City, UT 84112, USA

^3^ Huntsman Mental Health Institute, University of Utah, Salt Lake City, UT 84112, USA

^4^ Department of Psychology, University of Salzburg, 5020 Salzburg, Austria

^5^ Centre for Cognitive Neuroscience Salzburg, University of Salzburg, 5020 Salzburg, Austria

**^*^ Corresponding author**

Bradley King, PhD

Department of Health & Kinesiology, College of Health, University of Utah

250 S 1850 E

Salt Lake City, Utah 84112

Table of contents

APPENDIX 1: Supplementary methods	5

1.1 Registration deviations	5

Table S1. List of the deviations from the registered analyses followed by their justification.	5

1.2 Subject-specific deviations	6

Table S2. Deviations from the standard protocol.	6

1.3 Flow diagram of participant exclusions	8

Figure S1. Flow diagram of participant exclusions throughout the experiment.	8

1.4 Familiarization session	9

1.5 Serial Reaction Time Task (SRTT)	9

Figure S2. Motor sequence learning tasks	11

1.6 Finger Tapping Task (FTT) habituation	11

Figure S3. Finger Tapping Task (FTT) habituation.	13

1.7 Computation of Bayes Factors	14

1.8 fMRIPrep pre-processing details	14

1.9 Coordinates for small volume correction (SVC)	17

Table S3. Coordinates used for spherical small volume corrections on the results presented in the main text.	17

APPENDIX 2: Participant characteristics, sleep and vigilance	22

Table S4. Statistical output of age group comparisons in participant characteristics.	22

APPENDIX 3: Supplementary behavioral results	23

Table S5. Full statistical output for normalized task performance across all FTT task runs.	23

Figure S4. Non-normalized motor performance.	24

Table S6. Statistics output for absolute (non-normalized) task performance of all FTT task runs.	25

Figure S5. Average micro-online (a) and micro-offline (b) performance changes based on non-normalized transition times.	26

Figure S6. Macro-offline performance changes based on non-normalized performance.	27

Table S7. Statistics of one-sample t-tests assessing offline performance changes.	28

Figure S7. Normalized General Performance Index (GPI) across practice blocks for all FTT runs.	29

Table S8. Statistics output of block-to-block changes in the general performance index (GPI).	30

Figure S8. Macro-offline changes in the General Performance Index (GPI).	31

Table S9. Age-related changes in offline performance changes during childhood.	32

Figure S9. Age-related changes in macro-offline changes in speed.	32

APPENDIX 4: Head motion	33

Table S10. Head motion parameters across task runs.	33

Table S11. Statistics of group x task run ANOVAs on head motion parameters.	33

APPENDIX 5: Supplementary fMRI results	34

Table S12. Results of task-related activity during Training 1 across both age groups [task practice vs. interleaved rest].	34

Figure S10. Brain regions showing significant task-related brain activation in both age groups.	34

Table S13. Within-group results of task-related activity during Training 1 [task practice vs. interleaved rest]	35

Table S14. Uncorrected results of task-related activity during early learning (task practice vs. rest), with average transition time across practice blocks as a covariate.	37

Table S15. Regression analyses between brain responses during training 1 and micro-online performance changes [micro-online changes x training 1].	39

Table S16. Regression analyses between brain responses during training 1 and micro-offline performance changes [micro-offline changes x training 1].	40

Table S17. Additional results of the regression analysis of task activity during training 1 with the macro-offline performance changes.	41

Table S18. Task-related activity during training 2 [task practice vs. interleaved rest].	43

Table S19. Regression analyses between brain responses during training 2 and micro-online performance changes [micro-online changes x training 2].	45

Table S20. Regression analyses between brain responses during training 2 and micro-offline performance changes [micro-offline changes x training 2].	46

Table S21. Regression analyses between brain responses during training 2 and macro-offline performance changes [macro-offline changes x training 2].	47

Table S22. Changes in activity from training 1 to training 2 [Training 2 – Training 1].	48

Table S23. Regression analyses between activity changes from training 1 to training 2 and macro-offline performance changes [macro-offline changes x (training 2 – training 1)].	49

Table S24. Task-related activity during Retest [task practice vs. interleaved rest].	51

Table S25. Regression analyses between brain responses during retest and macro-offline performance changes [macro-offline changes x retest].	53

Table S26. Additional results of the inter-session changes in brain activity [Retest – Training 2].	54

Table S27. Additional results of the regression analysis inter-session changes with the macro-offline performance changes [(retest – training 2) x macro-offline performance changes].	56

Table S28. Age-related changes in brain responses in the child sample [task practice vs. interleaved rest].	58

APPENDIX 6: Supplementary discussion	59

6.1 Behavioral initial learning dynamics	59

6.2 Micro-offline performance changes	60

6.3 Potential explanations for the smaller modulations in brain activity between task and rest epochs in children	61

6.4 The relationship between brain activity during initial learning and micro-offline performance changes	62

6.5 Inter-session changes in brain activity in young adults	63

### APPENDIX 1: Supplementary methods

#### 1.1 Registration deviations

##### **Table S1**. List of the deviations from the registered analyses followed by their justification.

Deviations are marked with an asterisk in the main text.

|  | **Registered** | **Deviation** |
| --- | --- | --- |
| 1 | Spatial pre-processing was performed in **SPM 12 in Matlab**. Task-based and resting-state functional volumes of each participant are realigned to the first image of each MRI run and subsequently, realigned to the mean functional image computed across MRI runs of the respective participant using rigid body transformations. The mean image is co-registered to the pre-processed high-resolution T1-weighted structural image using a rigid body transformation optimized to maximize the normalized mutual information between the two images. The resulting co-registration parameters are then applied to the realigned functional images | Initial spatial preprocessing was performed in **fmriprep 24.1.0**, which overall performed similar pre-processing steps. However, the way that they were performed was slightly different and we implemented additional pre-processing such as **correction for inhomogeneities in the magnetic field using fieldmaps**. The main text contains a description of the pre-processing pipeline that was automatically generated by fmriprep. |
|  | **Justification**: Pre-processing was performed in fMRIPrep 24.1.0 instead of SPM12 to facilitate the implementation of correction for inhomogeneities in the magnetic field with the acquired field maps. Although SPM12 provides this option, prior researchers in our team experienced unforeseen image distortions in a small subset of task runs when implementing field maps using this software. | |
| 2 | The set of voxel values resulting from each second-level analysis described above (activation and regression analyses) will constitute maps of the t-statistics [SPM(T)], thresholded at **p < 0.005** (uncorrected for multiple comparisons). | The set of voxel values resulting from each second-level analysis constitutes maps of the t-statistics [SPM(T)], thresholded at **p < 0.001** (uncorrected for multiple comparisons) |
|  | **Justification:** The results were thresholded at a more restrictive p-value to reduce the risk of false positive results. | |

#### 1.2 Subject-specific deviations

##### **Table S2**. Deviations from the standard protocol.

There were occasional issues that arose during data acquisition, particularly in young children, that resulted in deviations from the standard protocol. A list of these subject-specific deviations is provided.

| **Subject** | **Age (yrs)** | **Deviation** |
| --- | --- | --- |
| E4 | 7.92 | The participant completed all 10 blocks of training 1; however, functional images were only acquired for the first 9 blocks since the pre-defined maximum number of scans was exceeded due to the slow performance of the participant. Only 9 practice blocks were performed for training 2 because the participant terminated this run.  Accordingly, there were missing behavioral data from this participant for block 10 in Training 2. This participant was thus excluded from the behavioral analyses assessing learning dynamics across the training runs.  There were missing neuroimaging data from blocks 10 in both Training runs 1 and 2. The onset file for our imaging analyses thus reflected this deviation. |
| E31 | 9.58 | The participant exhibited accuracy of less than 40% and a decrease in the time between transitions of over 150% percent during the post-learning test. Accordingly, the participant was no longer following task instructions (i.e., to perform the task as accurately and quickly as possible). As this behavior was only evident during the post-learning test phase (and not during training 1, training 2 or retest which were used in the fMRI analyses), we elected to include this participant in analyses but alter the computation of macro-offline performance changes. This metric was computed as the change in performance from the last two blocks of training 2 (in contrast to the average of the two test blocks) to the first two blocks of retest. Note that this change in the computation resulted in a substantially smaller macro-offline gain and thus was in the direction opposite to our hypothesized (and reported) results.  If we maintained the original computation (i.e., based on the average of the two test blocks), the significant group differences in macro-offline changes in performance speed and accuracy were as follows: Speed: t_43_ = 2.602, p = 0.006, G = 0.76, BF_10_ = 4.085; accuracy: t_43_ = 2.311, p = 0.015, G = 0.67, BF_10_ = 2.580.  Due to excessive head motion (i.e., maximum translation exceeding 5 mm), volumes 1 to 82 of Training 2 were included in the univariate GLM as a regressor of no interest. |
| E38 | 7.92 | The key on the keyboard that corresponds to #2 in the motor sequence (i.e., the left index finger) stopped working during training run 2 and thus, behavioral and neuroimaging data from this training run was discarded for this participant and thus not included in statistical analyses.  The retest run was restarted after one practice block due to excessive participant motion. Functional images as well as performance during this additional retest block were ignored. If computed based on the excluded block, the macro-offline performance changes would be as follows: speed: 0.248 instead of 0.087; accuracy: 0.032 instead of 0.023. |
| E39 | 27.33 | Training run 1 was restarted during practice block 1 because the keyboard was not connected to the computer. As these keypresses were not logged, functional images and performance during these trials were ignored. |
| E45 | 8.17 | The retest run was restarted at the start of the first practice block due to excessive motion. Functional images as well as performance during this additional retest block were ignored. |
| E49 | 10 | Due to excessive head motion (i.e., maximum translation exceeding 5 mm), volumes 719 to 914 of Training 2 were included in the univariate GLM as a regressor of no interest. |
| E103 | 10.83 | Participant lost the keys at the beginning of block 6 of training run 1. Scanning was stopped to reposition the fingers and the participant completed 5 additional blocks of training run 1. Functional images and performance during this portion of block 6 of the first part of training 1 were ignored. The participant’s general linear model (GLM) was built by combining the BOLD signal of the 5 remaining blocks of the two parts of training run 1, with equal weights. |
| E107 | 8.67 | Training run 1 was restarted after 2 blocks due to excessive motion. Functional images and performance during these two blocks were ignored.  Due to excessive head motion (i.e., maximum translation exceeding 5 mm), data from training runs 1 and 2 were not included in the univariate analyses. |
| E113 | 7.25 | Due to excessive head motion (i.e., maximum translation exceeding 5 mm), volumes 1008 to 1293 of Training 1 were included in the univariate GLM as a regressor of no interest. |

#### 1.3 Flow diagram of participant exclusions

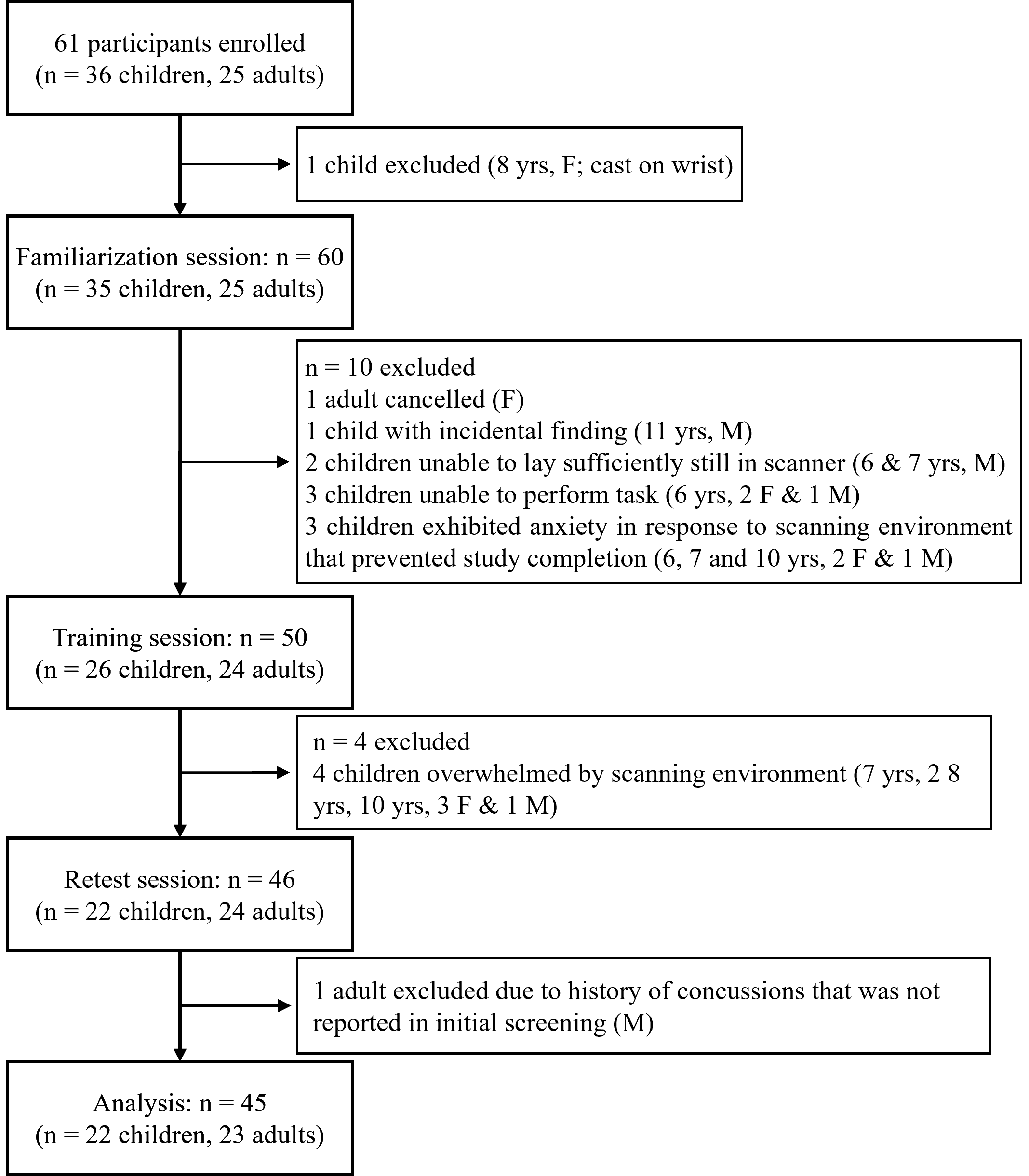

##### **Figure S1.** Flow diagram of participant exclusions throughout the experiment.

F = female, M = male. Note that most subjects were excluded due to the inability to lay sufficiently still in the scanner, correctly perform the motor task or becoming overwhelmed by the scanning environment. This primarily included younger children (i.e., 6- and 7-year-olds).

#### 1.4 Familiarization session

Following the informed consent process, verification that participants did not present any MRI contra-indications, and providing a general introduction to the study procedures, participants completed a series of tasks to become familiar with our experimental procedures. While seated in front of a laptop on a desk, participants first completed 5 blocks of the pseudorandom SRTT, followed by the FTT habituation procedure (see below for details on both the SRTT and FTT habituation). Next, participants were positioned in a mock scanner and were asked to lay as still as possible for a 5-minute mock resting scan during which EPI sounds were played, affording an opportunity for participants to become introduced to a simulated MR environment. Additionally, head motion was monitored with a custom-made motion detection system made up of a small camera and a commercial motion sensor, and auditory feedback was provided when this system detected bodily movement. Following the simulated resting state scan, participants completed another 5 blocks of the pseudorandom SRTT in the mock scanner with EPI sounds and the head motion detection system, further familiarizing them with the set-up of the real scanning environment (i.e., hands and keypads positioned at the thighs, use of a mirror on a head coil to view the rear-positioned monitor, etc.).

#### 1.5 Serial Reaction Time Task (SRTT)

To familiarize the participants with the use of two separate keyboards and the specific four fingers (2 fingers in each hand) used for the bimanual finger tapping task described in the main text, participants performed a modified version of the serial reaction time task (SRTT; see Figure S2; Reverberi et al., 2023). The task consisted of a 4-choice reaction time task in which participants were instructed to react to cues shown on the computer screen. Four squares, spatially corresponding to the 4 fingers used to perform the task (i.e., index and middle fingers of both hands) were presented on the screen. The color of the outline of the squares alternated between red and green, indicating blocks of rest or practice, respectively. Practice blocks consisted of 48 key presses and were separated by 20-second rest periods. During practice blocks, a visual stimulus appeared in one of the squares, and participants were instructed to press the corresponding key with the corresponding finger as quickly and as accurately as possible. The stimuli appeared in an order that pseudo-randomly changed every 4 elements (i.e., the stimulus appeared in each location once every 4 elements and never appeared in the same location consecutively). Pseudorandom sequences (48 elements) were generated ahead of time and the order in which sequences were presented was randomized each random run. Thus, all participants performed the same sequences of keypresses that appeared in a pseudorandom order, but the order of these sequences was randomized. Given the pseudorandom order of the stimuli, the general motor execution of keypresses on a keyboard was assessed without engaging any sequence learning processes.

During performance of the SRTT, keypresses and their specific timing were recorded. Thus, for each stimulus (i.e., trial), response time (i.e., time between each stimulus and corresponding press; RT) and accuracy (i.e., correct or incorrect) were measured. However, the sole purpose of this task was to familiarize participants with the use of the keyboards prior to completing the finger tapping task (FTT) employed to assess motor sequence learning and memory consolidation processes. Accordingly, data from the SRTT were not analyzed.

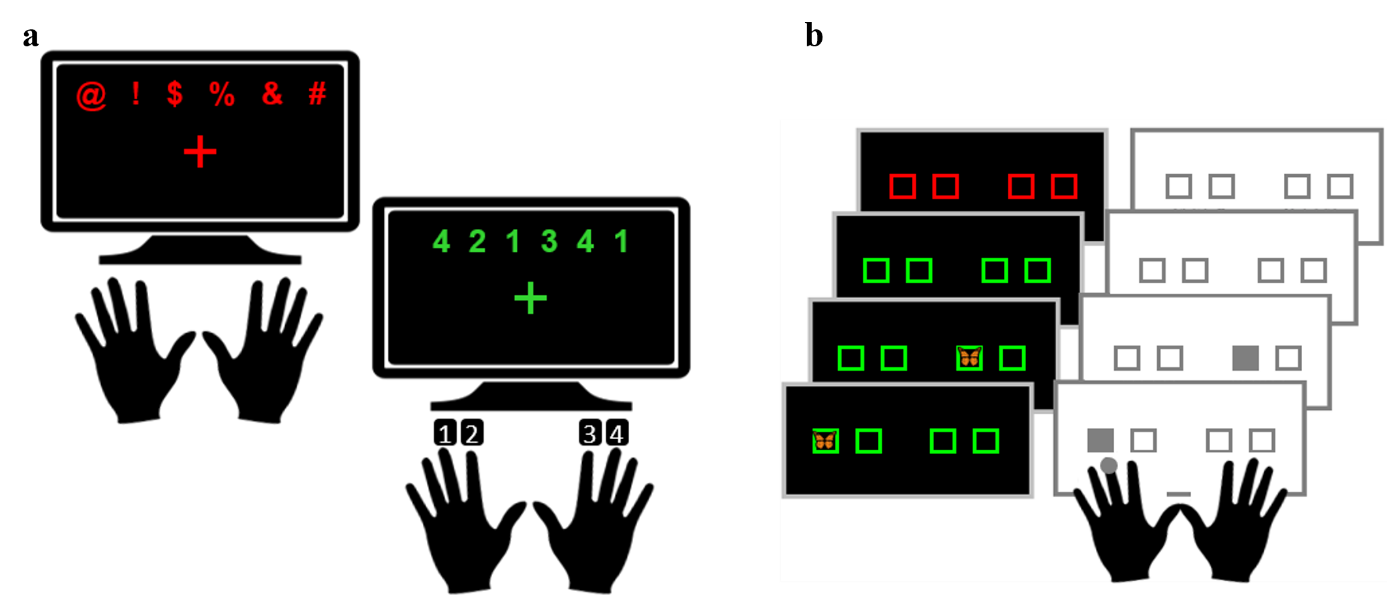

**Figure S2**. Motor sequence learning tasks

(**a**) Finger tapping task (FTT). When the cross turned green, participants repeatedly pressed the keys in the order of the numbers presented on the screen. Participants stopped when the cross turned red, indicting a rest epoch. (**b**) Serial Reaction Time Task (SRTT). A stimulus appears in 1 of 4 spatial locations and the participant responds with the corresponding key/finger as fast as possible. To increase motivation and attention the task was built around a story in which participants were asked to catch butterflies. Butterfly images were taken from [https://openclipart.org](https://openclipart.org/detail/224784/monarch-butterfly) and fall under the Creative Commons (CC) Zero 1.0 Public Domain License. Hand and computer screen images were taken from https://thenounproject.com and fall under the CC 3.0 Unported License.

#### 1.6 Finger Tapping Task (FTT) habituation

To ensure participants comprehended the instructions and were comfortable with the FTT prior to completing the experimental sessions in the MRI scanner, we designed a series of practice runs that incrementally introduced the FTT (Figure S3 below).

1. Participants were asked to complete the simple sequence 1-2-3-4, where the 1=left middle finger, 2=left index finger, 3=right index finger and 4= right middle finger), slowly and accurately until instructed to stop. This trial started with a screen that provided the sequence to-be-performed as well as an image that depicted the fingers to be used and their corresponding numbers. The text was presented in red, indicating that participants were not supposed to move (see Panel A in Figure S3 below). After a delay of 20 seconds, the text changed to green (Panel B) and participants performed the sequence slowly and accurately. Unbeknownst to the participant, this task run ended after the sequence was performed correctly 3 times in a row.
2. Participants were again asked to perform the same sequence as described above (i.e., 1-2-3-4), with the sole difference that the image showing the fingers to be used and their corresponding numbers was not shown (Panel C in Figure S3). Participants thus needed to know the finger-to-number mapping (i.e., 1 is the left middle finger, 2 is the left index finger, etc.).
3. Participants then completed steps (1) and (2) above, but with a different, albeit equally simple, sequence (i.e., 4-3-2-1).
4. Participants were then provided with a 6-element sequence – analogous to what was performed during the real experimental sessions inside the MR scanner. This practice sequence was 3-1-2-4-3-2 and was specifically chosen to not share any transitions with the real sequence 4-2-1-3-4-1. First, and similar to above, this sequence was presented on the screen with the image depicting the fingers to be used and their corresponding numbers immediately below. The text was initially in red and thus the participants were asked not to move (Panel D in Figure S3). Following a delay of 20 seconds, the text changed to green (Panel E) and participants were instructed to complete this sequence slowly and accurately until the color changed back to red. Subsequently, this was done without (Panel F) the image showing the fingers to be used and their corresponding numbers. Both task runs concluded after the sequence was performed correctly three times in a row.
5. Participants were instructed to complete the same 6-element sequence a little faster in order to resemble how the FTT would be performed during the imaging sessions. The image showing the fingers to be used and their corresponding numbers were not shown, and the task ended after the sequence was performed correctly five times in a row.

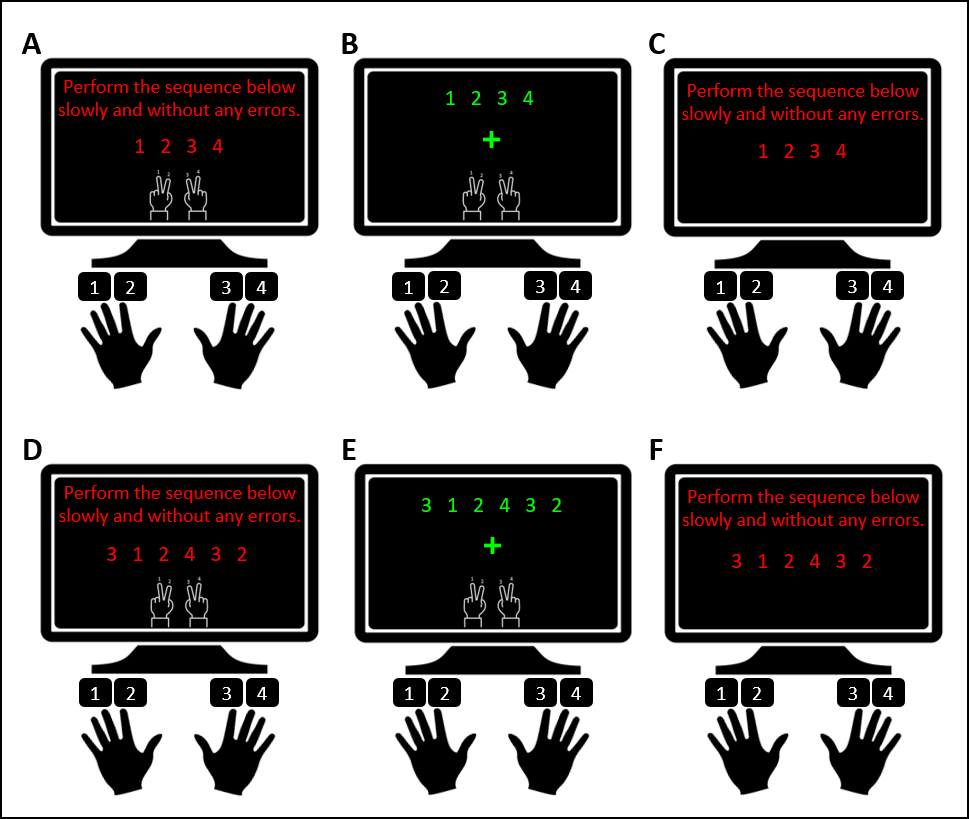

**Figure S3**. Finger Tapping Task (FTT) habituation.

Participants use the index and middle fingers of both hands to perform the shown sequence slowly and accurately until instructed to stop. Each trial started with a screen that provided the sequence to perform and an image depicting the fingers to be used and their corresponding numbers. The text was presented in red, indicating that participants were not supposed to move (Panels A & D). After a delay of 20 seconds, the text changed to green and participants performed the sequence slowly and accurately (Panels B & E). Participants were then asked to perform the same sequence again, with the sole difference that the image depicting the fingers to be used and their corresponding numbers was not shown (Panels C & F). Each task run ended after the sequence was performed correctly 3 times in a row. Participants practiced two simple 4-element sequences (one of them shown in Panels A-C), followed by a 6-element sequence (Panels D-F). Hand and computer screen images were taken from https://thenounproject.com and fall under the CC 3.0 Unported License.

#### 1.7 Computation of Bayes Factors

Null hypothesis testing for all behavioral effects presented in the main text was supplemented by the computation of Bayes Factors (BF; BayesFactor R-package), reflecting the likelihood of the observed data favoring an alternative model (e.g., evidence for differences among groups, blocks, etc.) relative to the null model (i.e., no differences). The null models for the main effects of the t-tests, one-way ANOVAs and simple linear regressions, and mixed model ANOVAs were specified as the difference between means equals zero, the intercept term, and the random effect of subjects, respectively. For the interaction effects, the reported BFs reflect the comparison of two models with and without the interaction term of interest. Whereas most parameters were set to the default options, the “whichModels” parameter was set to “all” for mixed model ANOVAs and multiple regressions to compute the appropriate BFs. In our results, we report BF10 values, with larger values indicating a greater likelihood that the observed data favors the alternative as compared to the null hypotheses.

#### 1.8 fMRIPrep pre-processing details

The text below was automatically generated by fMRIPrep (Esteban et al., 2019) in each participant’s visual report and is released under the CC0 license. Additions or clarifications to this automated text made by the authors of this manuscript are explicitly noted below.

A total of 2 fieldmaps were found available within the input BIDS structure for this particular subject. A *B_0_* nonuniformity map (or *fieldmap*) was estimated from the phase-drift map(s) measure with two consecutive GRE (gradient-recalled echo) acquisitions. The corresponding phase-map(s) were phase-unwrapped with prelude (FSL None).

A total of 2 T1-weighted (T1w) images were found within the input BIDS dataset [Author note: The acquired T1w image was duplicated, creating 2 T1w images to have 1 within the directory for each testing session (i.e., initial training vs. 5hr retest)]. Each T1w image was corrected for intensity non-uniformity (INU) with N4BiasFieldCorrection (Tustison et al., 2010), distributed with ANTs 2.5.3 (Avants et al., 2008; RRID:SCR_004757). The T1w-reference was then skull-stripped with a *Nipype* implementation of the antsBrainExtraction.sh workflow (from ANTs), using OASIS30ANTs as target template. Brain tissue segmentation of cerebrospinal fluid (CSF), white-matter (WM) and gray-matter (GM) was performed on the brain-extracted T1w using fast (FSL (version unknown), RRID:SCR_002823, Zhang et al., 2001). An anatomical T1w-reference map was computed after registration of 2 <module ‘nipype.interfaces.image’ from ‘/opt/conda/envs/fmriprep/lib/python3.11/site-packages/nipype/interfaces/image.py’> images (after INU-correction) using mri_robust_template (FreeSurfer 7.3.2, Reuter et al., 2010). Volume-based spatial normalization to one standard space (MNI152NLin2009cAsym) was performed through nonlinear registration with antsRegistration (ANTs 2.5.3), using brain-extracted versions of both T1w reference and the T1w template. The following template was selected for spatial normalization and accessed with *TemplateFlow* (24.2.0, Ciric et al., 2022): *ICBM 152 Nonlinear Asymmetrical template version 2009c* [Fonov et al., 2009, RRID:SCR_008796; TemplateFlow ID: MNI152NLin2009cAsym]. [Author note: The spatial normalization to MNI space described above was not used for the analyses presented in this research. Rather, we used the fMRIPrep-processed images in native space and normalized these to a sample-specific template using DARTEL in SPM 12 (described in Section 2.6.2)].

For each of the 7 BOLD runs found per subject (across all tasks and sessions), the following preprocessing was performed. [Author note: RS runs were included in the preprocessing to facilitate other analyses that included these runs but are not included in this paper]. First, a reference volume was generated, using a custom methodology of *fMRIPrep*, for use in head motion correction. Head-motion parameters with respect to the BOLD reference (transformation matrices, and six corresponding rotation and translation parameters) are estimated before any spatiotemporal filtering using mcflirt (FSL, Jenkinson et al., 2002). The estimated *fieldmap* was then aligned with rigid-registration to the target EPI (echo-planar imaging) reference run. The field coefficients were mapped on to the reference EPI using the transform. The BOLD reference was then co-registered to the T1w reference using mri_coreg (FreeSurfer) followed by flirt (FSL, Jenkinson & Smith, 2001) with the boundary-based registration (Greve & Fischl, 2009) cost-function. Co-registration was configured with six degrees of freedom.

Several confounding time-series were calculated based on the *preprocessed BOLD*. [Author note: The confounds file produced by fMRIPrep contained numerous metrics not used in these analyses and thus omitted from the text below.] The head-motion estimates calculated in the correction step were also placed within the corresponding confounds file. All resamplings can be performed with *a single interpolation step* by composing all the pertinent transformations (i.e. head-motion transform matrices, susceptibility distortion correction when available, and co-registrations to anatomical and output spaces). Gridded (volumetric) resamplings were performed using nitransforms, configured with cubic B-spline interpolation.

#### 1.9 Coordinates for small volume correction (SVC)

##### **Table S3.** Coordinates used for spherical small volume corrections on the results presented in the main text.

| **Area** | **x** | **y** | **z** |  |
| --- | --- | --- | --- | --- |
| Anterior orbital | -24 | 34 | -12 | (Dandolo & Schwabe, 2019) |
| Lateral orbital | -48 | 50 | -10 | (Dolfen et al., 2021) |
| Posterior orbital | -36 | 24 | -12 | (Albouy et al., 2015) |
| Inferior frontal | -54 | 22 | 12 | (Dolfen et al., 2021) |
|  | -54 | 36 | 12 | (Dandolo & Schwabe, 2019) |
|  | -46 | 30 | 12 | (Penhune & Doyon, 2005) |
|  | 30 | 48 | -16 | (Veldman et al., 2023) |
|  | 56 | 12 | 8 | (Albouy et al., 2008) |
|  | 52 | 14 | 14 | (Albouy et al., 2008) |
|  | 52 | 46 | 2 | (Dolfen et al., 2021) |
|  | -46 | 18 | 32 | (Dolfen et al., 2021) |
|  | -42 | -2 | 32 | (Cousins et al., 2016) |
|  | -56 | 30 | 0 | (Albouy et al., 2015) |
|  | -40 | 48 | -2 | (King et al., 2020) |
|  | -54 | 22 | 12 | (Dolfen et al., 2021) |
|  | -24 | 30 | 10 | (Fogel et al., 2014) |
|  | 36 | 46 | -4 | (Albouy et al., 2008) |
| Middle frontal | -32 | 14 | 50 | (Penhune & Doyon, 2005) |
|  | 44 | 40 | 0 | (Penhune & Doyon, 2005) |
|  | 46 | 46 | 26 | (Veldman et al., 2023) |
|  | 54 | 6 | 44 | (Albouy et al., 2008) |
|  | -36 | 32 | 42 | (Veldman et al., 2023) |
|  | 44 | 40 | 36 | (Veldman et al., 2023) |
|  | -36 | 28 | 48 | (Fogel et al., 2014) |
|  | -22 | 62 | 22 | (Dandolo & Schwabe, 2019) |
|  | -30 | 46 | 22 | (Dolfen et al., 2021) |
|  | -28 | 16 | 40 | (Dolfen et al., 2021) |
|  | 40 | 56 | 16 | (Dolfen et al., 2021) |
|  | 54 | 6 | 44 | (Albouy et al., 2008) |
| Medial frontal | 10 | 48 | -8 | (Albouy et al., 2015) |
| Superior frontal | -16 | 42 | 48 | (Hedenius & Persson, 2022) |
|  | -24 | -14 | 52 | (Orban et al., 2010) |
|  | -32 | 54 | 20 | (Dolfen et al., 2021) |
|  | 30 | 18 | 40 | (Dolfen et al., 2021) |
|  | -24 | 68 | 4 | (Dolfen et al., 2021) |
|  | -26 | 48 | 34 | (Albouy et al., 2008) |
|  | 28 | 40 | 30 | (Dolfen et al., 2021) |
| Medial superior frontal | -2 | 62 | -8 | (Hedenius & Persson, 2022) |
|  | 10 | 68 | 20 | (Dolfen et al., 2021) |
|  | -8 | 66 | 8 | (Albouy et al., 2013) |
|  | -6 | 56 | 40 | (Dolfen et al., 2021) |
|  | -6 | 60 | 28 | (King et al., 2020) |
|  | -2 | 36 | 32 | (Albouy et al., 2015) |
|  | -2 | 40 | 56 | (Dandolo & Schwabe, 2019) |
|  | -14 | -48 | 44 | (Hedenius & Persson, 2022) |
|  | -14 | 54 | 36 | (King et al., 2020) |
|  | 14 | -40 | 40 | (Albouy et al., 2015) |
| Supplementary motor area | -2 | 4 | 62 | (Dolfen et al., 2021) |
|  | 2 | -2 | 70 | (Penhune & Doyon, 2005) |
|  | -2 | 10 | 56 | (Penhune & Doyon, 2005) |
|  | -8 | -12 | 56 | (Nicolas et al., 2024) |
|  | 8 | -4 | 50 | (Nicolas et al., 2024) |
| Precentral | 36 | -18 | 62 | (Albouy et al., 2013) |
|  | -54 | 4 | 36 | (King et al., 2020) |
|  | 54 | 6 | 44 | (Albouy et al., 2008) |
|  | -26 | -22 | 68 | (Albouy et al., 2015) |
|  | 22 | -14 | 72 | (Nicolas et al., 2024) |
|  | -50 | -8 | 24 | (Albouy et al., 2015) |
|  | 42 | -4 | 44 | (Nicolas et al., 2024) |
|  | -50 | -8 | 38 | (Nicolas et al., 2024) |
|  | 48 | 8 | 30 | (Albouy et al., 2008) |
|  | -30 | -6 | 52 | (Nicolas et al., 2024) |
| Postcentral | 48 | -10 | 30 | (Veldman et al., 2023) |
|  | 18 | -42 | 70 | (Penhune & Doyon, 2005) |
|  | -26 | -36 | 50 | (Dolfen et al., 2021) |
|  | -44 | -36 | 62 | (Veldman et al., 2023) |
|  | 32 | -38 | 72 | (Dolfen et al., 2021) |
|  | -50 | -40 | 60 | (Albouy et al., 2008) |
|  | 68 | -26 | 34 | (Albouy et al., 2008) |
| Anterior cingulate | -4 | 6 | 40 | (Albouy et al., 2008) |
|  | -20 | 40 | 14 | Cousins et al., 2016) |
| Middle cingulate | -6 | -44 | 38 | (King et al., 2020) |
|  | -14 | -14 | 34 | (Cousins et al., 2016) |
|  | -8 | -42 | 46 | (Albouy et al., 2013) |
|  | 14 | -40 | 40 | (Albouy et al., 2015) |
| Posterior cingulate | -4 | -54 | 28 | (Hedenius & Persson, 2022) |
|  | 2 | -32 | 30 | (Dolfen et al., 2021) |
|  | 6 | -38 | 14 | (Fogel et al., 2014) |
| Paracentral lobule | -12 | -18 | 74 | (Penhune & Doyon, 2005) |
|  | -12 | -32 | 60 | (Nicolas et al., 2024) |
|  | -12 | -22 | 80 | (Dandolo & Schwabe, 2019) |
|  | 2 | -32 | 30 | (Dolfen et al., 2021) |
| Inferior parietal | 60 | -36 | 46 | (King et al., 2016) |
|  | -42 | -40 | 36 | (Dolfen et al., 2021) |
|  | -44 | -48 | 38 | (Dolfen et al., 2021) |
| Superior parietal | 24 | -62 | 50 | (Nicolas et al., 2024) |
|  | -14 | -54 | 72 | (Albouy et al., 2008) |
|  | 28 | -52 | 68 | (King et al., 2020) |
| Supramarginal | 68 | -34 | 30 | (Dolfen et al., 2021) |
|  | 44 | -36 | 56 | (King et al., 2016) |
|  | 50 | -42 | 34 | (Dolfen et al., 2021) |
|  | 52 | -40 | 26 | (Dolfen et al., 2021) |
|  | 50 | -40 | 26 | (King et al., 2020) |
|  | 58 | -18 | 28 | (Nicolas et al., 2024) |
|  | -54 | -22 | 18 | (Byczynski et al., 2025) |
|  | -56 | -32 | 32 | (Nicolas et al., 2024) |
|  | -58 | -24 | 34 | (Albouy et al., 2008) |
|  | 64 | -28 | 30 | (Albouy et al., 2008) |
| Angular | -48 | -72 | 34 | (King et al., 2020) |
|  | 52 | -66 | 32 | (King et al., 2020) |
|  | -30 | -62 | 42 | (Dolfen et al., 2021) |
| Precuneus | 18 | -56 | 54 | (King et al., 2016) |
|  | 16 | -74 | 26 | (Albouy et al., 2008) |
|  | 8 | -56 | 20 | (King et al., 2020) |
|  | 12 | -50 | 34 | (Albouy et al., 2015) |
|  | -18 | -62 | 32 | (King et al., 2016) |
| Thalamus | -20 | -10 | 0 | (King et al., 2016) |
|  | -20 | -12 | 18 | (Albouy et al., 2008) |
| Caudate nucleus | -18 | 8 | 22 | (Nicolas et al., 2024) |
|  | 6 | 18 | 2 | (Albouy et al., 2015) |
| Putamen | -36 | 0 | 2 | (Nicolas et al., 2024) |
| Hippocampus | 32 | -38 | -6 | (Nicolas et al., 2024) |
|  | -28 | -20 | -16 | (Nicolas et al., 2024) |
| Parahippocampus | 33 | -18 | -27 | (Schendan et al., 2003) |
|  | -16 | -38 | -8 | (Dandolo & Schwabe, 2019) |
| Fusiform | -34 | -46 | -10 | (King et al., 2020) |
|  | -34 | -36 | -26 | (Dandolo & Schwabe, 2019) |
|  | -32 | -16 | -36 | (King et al., 2020) |
| Amygdala | -28 | -6 | -14 | (Dandolo & Schwabe, 2019) |
|  | -28 | -6 | -16 | (Dandolo & Schwabe, 2019) |
| Cerebellum 4-5 | -4 | -58 | -12 | (Albouy et al., 2013) |
|  | 16 | -48 | -20 | (Albouy et al., 2008) |
|  | -12 | -54 | -18 | (King et al., 2020) |
| Cerebellum 6 | -22 | -64 | -26 | (Penhune & Doyon, 2005) |
|  | 0 | -74 | -14 | (Albouy et al., 2008) |
|  | 36 | -60 | -21 | (Debas et al., 2010) |
| Cerebellum 7 | -36 | -66 | -40 | (Dandolo & Schwabe, 2019) |
|  | -22 | -86 | -46 | (Albouy et al., 2008) |
|  | -18 | -72 | -46 | (Albouy et al., 2008) |
| Cerebellum 8 | -20 | -64 | -52 | (King et al., 2020) |
|  | 28 | -54 | -50 | (Penhune & Doyon, 2005) |
|  | 22 | -62 | -52 | (King et al., 2020) |
|  | 26 | -50 | -48 | (Fogel et al., 2014) |
| Cerebellum 9 | 14 | -56 | -60 | (Veldman et al., 2023) |
|  | 16 | 50 | -44 | (Dandolo & Schwabe, 2019) |
|  | -18 | -50 | -26 | (Albouy et al., 2013) |
| Cerebellum crus I | 36 | -66 | -40 | (Dandolo & Schwabe, 2019) |
|  | 36 | -60 | -21 | (Orban et al., 2010) |
|  | 38 | -40 | -32 | (Penhune & Doyon, 2005) |
|  | 40 | -76 | -20 | (Dolfen et al., 2021) |
| Cerebellum crus II | 30 | -72 | -34 | (Dolfen et al., 2021) |
|  | 26 | -80 | -38 | (Albouy et al., 2015) |
| Vermis 4-5 | 6 | -52 | -6 | (Penhune & Doyon, 2005) |
| Vermis 6 | -10 | -74 | -18 | (Albouy et al., 2008) |
|  | 2 | -64 | -18 | (Dolfen et al., 2021) |

x, y and z coordinates are specified in MNI space. A subset of these coordinates was applied to results from the contralateral hemisphere.

### APPENDIX 2: Participant characteristics, sleep and vigilance

#### Table S4. Statistical output of age group comparisons in participant characteristics.

| **Variable** |  | **df** | **F/t/W** | **p-value** | **ƞ^2^/G** |
| --- | --- | --- | --- | --- | --- |
| M/E preference |  | / | 276 | 0.582 | / |
| Time of testing | Training | 43 | -0.249 | 0.598 | -0.07 |
|  | Retest | 43 | 0.542 | 0.295 | 0.16 |
| Offline period |  | 43 | -0.186 | 0.574 | -0.05 |
| Sleep quality |  | / | 307.5 | 0.173 | / |
| Sleep duration |  | 43 | 5.856 | < 0.001* | 1.72 |
| SSS score | Group | 1,43 | 0.03 | 0.864 | < 0.001 |
|  | Session | 1,43 | 1.38 | 0.246 | 0.013 |
|  | Group x session | 1,43 | 1.38 | 0.246 | 0.013 |
| PVT score | Group | 1,42 | 49.92 | < 0.001* | 0.508 |
|  | Session | 1,42 | 0.08 | 0.780 | < 0.001 |
|  | Group x session | 1,42 | 0.18 | 0.674 | < 0.001 |

Results from statistical analyses of Morning/Eveningness (M/E) preference, time of testing of both practice sessions, offline period duration, sleep quality, and sleep duration, as well as subjective and objective vigilance measures. Group x session ANOVAs were performed for the SSS and PVT scores. Based on whether the data was normally distributed, Mann-Whitney U tests were conducted for M/E preference and sleep quality, whereas other measures were assessed with independent samples t-tests. Significant values are marked with an asterisk. Df = degrees of freedom; ƞ^2^ = eta squared; SSS = Stanford Sleepiness Scale (Maclean et al., 1992). PVT = psychomotor vigilance task (Dinges & Powell, 1985). Corresponding group means are provided in Table 1 of the main text. Most participant characteristics were comparable between the two age groups. Sleep duration was significantly longer in children as compared to adults, which is consistent with the known decrease in total sleep time with age (Giddens et al., 2022; Ohayon et al., 2004). Additionally, PVT reaction times were significantly slower in children than adults, which is also in line with previous research (Iida et al., 2010).

### APPENDIX 3: Supplementary behavioral results

#### Table S5. Full statistical output for normalized task performance across all FTT task runs.

|  | **Transition time** | | | | **% Correct transitions** | | | |
| --- | --- | --- | --- | --- | --- | --- | --- | --- |
| **Effect** | **df** | **F** | **p** | **ƞ^2^** | **df** | **F** | **p** | **ƞ^2^** |
| *A. Training* | | |  |  |  |  |  |  |
| Group | 1,42 | 7.85 | 0.008* | 0.091 | 1,42 | 1.22 | 0.276 | 0.016 |
| Run | 1,42 | 177.82 | <0.001* | 0.308 | 1,42 | 1.25 | 0.270 | 0.001 |
| Block | 4.87,204.55 | 22.43 | <0.001* | 0.092 | 5.18,217.70 | 1.82 | 0.107 | 0.009 |
| G x R | 1,42 | 1.41 | 0.242 | 0.004 | 1,42 | 4.42 | 0.041* | 0.004 |
| G x B | 4.87,204.55 | 0.68 | 0.637 | 0.003 | 5.18,217.70 | 2.75 | 0.018* | 0.014 |
| R x B | 5.27,221.14 | 8.54 | <0.001* | 0.033 | 5.27,221.29 | 1.25 | 0.286 | 0.005 |
| G x R x B | 5.27,221.14 | 2.14 | 0.058^+^ | 0.009 | 5.27,221.29 | 0.73 | 0.609 | 0.003 |
| *B. Test* | |  |  |  |  |  |  |  |
| Group | 1,43 | 2.01 | 0.164 | 0.043 | 1,43 | 0.42 | 0.522 | 0.009 |
| Block | 1,43 | 2.37 | 0.131 | 0.002 | 1,43 | 0.93 | 0.339 | 0.002 |
| G x B | 1,43 | 0.60 | 0.444 | <0.001 | 1,43 | 1.57 | 0.217 | 0.003 |
| *C. Retest* | | |  |  |  |  |  |  |
| Group | 1,43 | 0.03 | 0.873 | <0.001 | 1,43 | 0.07 | 0.796 | 0.001 |
| Block | 2.48,106.72 | 4.91 | 0.005* | 0.017 | 6,257.90 | 1.05 | 0.393 | 0.004 |
| G x B | 2.48,106.72 | 0.97 | 0.398 | 0.003 | 6,257.90 | 1.41 | 0.209 | 0.005 |

Results from statistical analyses of normalized transition time and % of correct transitions for all FTT task runs. Significant values are marked with an asterisk, non-significant trends are marked with a plus sign. Df = degrees of freedom; ƞ^2^ = eta squared; G x R = group x training run interaction; G x B = group x block interaction; R x B = training run x block interaction; G x R x B = group x training run x block interaction.

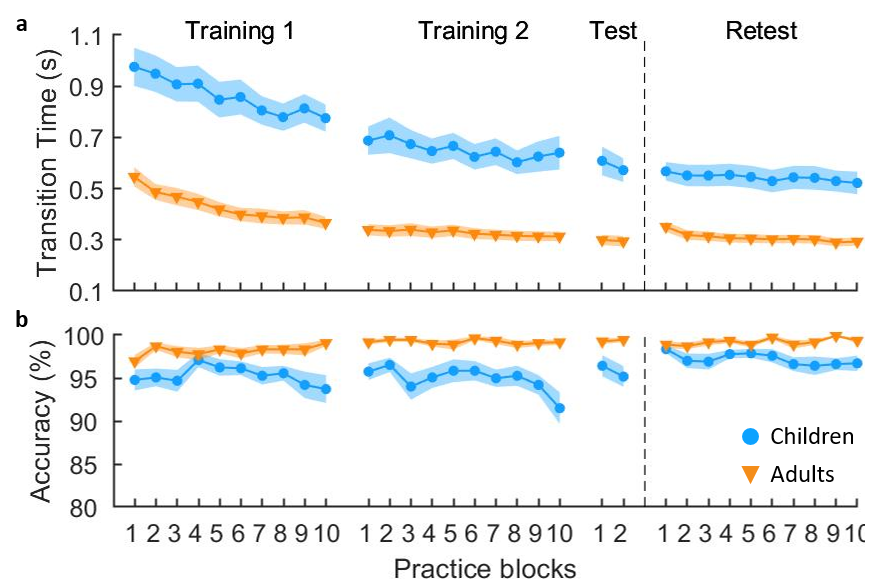

#### **Figure S4. Non-normalized motor performance.**

(**a**) Non-normalized transition time in seconds (s) for all FTT runs. (**b**) Percentage of correct transitions for all FTT runs. Shaded regions represent the standard errors of the mean. Results from the corresponding statistical analyses are shown in Supplementary Table S3.2. n = 22 children and 23 adults.

#### Table S6. Statistics output for absolute (non-normalized) task performance of all FTT task runs.

|  | **Transition time** | | | | **% Correct transitions** | | | |
| --- | --- | --- | --- | --- | --- | --- | --- | --- |
| **Effect** | **df** | **F** | **p** | **ƞ^2^** | **df** | **F** | **p** | **ƞ^2^** |
| *A. Training* | | |  |  |  |  |  |  |
| Group | 1,42 | 43.11 | <0.001* | 0.444 | 1,42 | 21.30 | <0.001* | 0.168 |
| Run | 1,42 | 111.86 | <0.001* | 0.119 | 1,42 | 1.16 | 0.288 | 0.002 |
| Block | 4.11,172.66 | 12.57 | <0.001* | 0.028 | 5.51,231.23 | 1.77 | 0.113 | 0.012 |
| G x R | 1,42 | 10.32 | 0.003* | 0.012 | 1,42 | 4.68 | 0.036* | 0.007 |
| G x B | 4.11,172.66 | 0.34 | 0.858 | <0.001 | 5.51,231.23 | 2.84 | 0.013* | 0.020 |
| R x B | 5.72,240.16 | 5.87 | <0.001* | 0.010 | 5.76,241.98 | 1.17 | 0.325 | 0.007 |
| G x R x B | 5.72,240.16 | 0.60 | 0.724 | 0.001 | 5.76,241.98 | 0.69 | 0.652 | 0.004 |
| *B. Test* | |  |  |  |  |  |  |  |
| Group | 1,43 | 30.57 | <0.001* | 0.403 | 1,43 | 10.86 | 0.002* | 0.163 |
| Block | 1,43 | 2.79 | 0.102 | 0.003 | 1,43 | 0.83 | 0.366 | 0.004 |
| G x B | 1,43 | 1.60 | 0.212 | 0.002 | 1,43 | 1.51 | 0.226 | 0.008 |
| *C. Retest* | | |  |  |  |  |  |  |
| Group | 1,43 | 29.80 | <0.001* | 0.384 | 1,43 | 15.58 | <0.001* | 0.122 |
| Block | 3.65,156.79 | 3.78 | 0.007* | 0.009 | 6.17,265.42 | 1.04 | 0.40 | 0.015 |
| G x B | 3.65,156.79 | 0.35 | 0.828 | <0.001 | 6.17,265.42 | 1.47 | 0.186 | 0.021 |

Results from statistical analyses of non-normalized transition time and % of correct transitions for all FTT task runs. Significant values are marked with an asterisk. Df = degrees of freedom; ƞ^2^ = eta squared; G x R = group x training run interaction; G x B = group x block interaction; R x B = training run x block interaction; G x R x B = group x training run x block interaction.

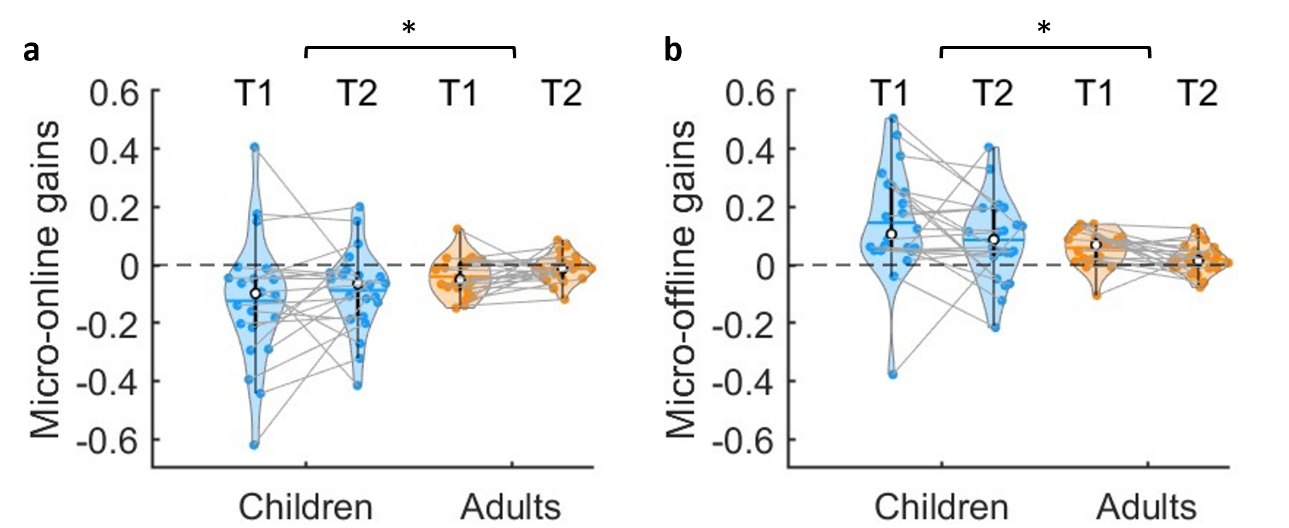

#### Figure S5. Average micro-online (a) and micro-offline (b) performance changes based on non-normalized transition times.

Positive values are indicative of performance improvements. Shaded regions represent the kernel density estimates of the data, colored circles depict individual data, open circles represent group medians, and the horizontal lines depict group means (Bechtold et al., 2021). n = 22 children and 23 adults. There was a significant age group main effect for micro-online performance changes (F_(1,43)_=5.81, p =0.020, ƞ^2^ = 0.082, BF_10_ = 2.724), with children exhibiting smaller (i.e., more negative) performance changes within practice blocks. Effects of training run (F_(1,43)_=1.97, p =0.17, ƞ^2^ = 0.015, BF_10_ = 0.53) and training run by group interaction (F_(1,43)_=0.03, p =0.87, ƞ^2^ < 0.001, BF_10_ = 0.20) were not significant. For micro-offline changes, the training run by group interaction was also not significant (F_(1,43)_=0.18, p =0.68, ƞ^2^ = 0.002, BF_10_ = 0.29). There were, however, significant main effects of both age group (F_(1,43)_=7.34, p =0.010, ƞ^2^ = 0.094, BF_10_ = 4.052) and training run (F_(1,43)_=4.72, p =0.035, ƞ^2^ = 0.041, BF_10_ = 1.842), demonstrating that children exhibited larger micro-offline performance gains as compared to adults and micro-offline gains were larger in training run 1 than training run 2.

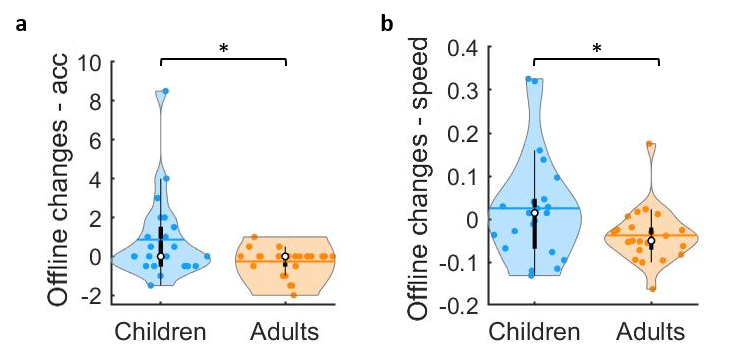

#### Figure S6. Macro-offline performance changes based on non-normalized performance.

Based on non-normalized accuracy (**a**) and transition times (**b)**. Positive values are indicative of performance improvements from end of training to the 5-hour delayed retest. Shaded regions represent the kernel density estimates of the data, colored circles depict individual data, open circles represent group medians, and the horizontal lines depict group means (Bechtold et al., 2021). n = 22 children and 23 adults. Results were the same as for the macro-offline measures based on normalized performance outcomes presented in the main text. There was a significant group effect for the macro-offline performance changes in speed (t_30.956_ = 2.128, p = 0.021, G = 0.62, BF_10_ = 1.841) and accuracy (t_25.243_ = 2.325, p = 0.014, G = 0.68, BF_10_ = 2.645), with children demonstrating greater improvements than adults.

#### Table S7. Statistics of one-sample t-tests assessing offline performance changes.

| **Variable** |  | **df** | **t** | **p** | **BF_10_** |
| --- | --- | --- | --- | --- | --- |
| *A. Micro-online changes* | | | | |  |
| Children | Training 1 | 21 | -3.373 | 0.003* | 14.328 |
|  | Training 2 | 21 | -3.115 | 0.005* | 8.583 |
| Adults | Training 1 | 22 | -3.468 | 0.002* | 17.951 |
|  | Training 2 | 22 | -1.193 | 0.246 | 0.411 |
| *B. Micro-offline changes* | | | | |  |
| Children | Training 1 | 21 | 4.097 | 0.001* | 63.972 |
|  | Training 2 | 21 | 3.078 | 0.006* | 7.989 |
| Adults | Training 1 | 22 | 5.149 | < 0.001* | 670.204 |
|  | Training 2 | 22 | 1.769 | 0.091^+^ | 0.834 |
| *C. Macro-offline changes in speed* | | | | |  |
| Children |  | 21 | -0.519 | 0.610 | 0.252 |
| Adults |  | 22 | -3.703 | 0.001* | 29.332 |
| *D. Macro-offline changes in accuracy* | | | |  |  |
| Children |  | 21 | 1.986 | 0.060^+^ | 1.161 |
| Adults |  | 22 | -1.772 | 0.090^+^ | 0.838 |

Results of one-sample t-tests to statistically assess micro-online, micro-offline and macro-offline performance changes. Test value = 0. Significant values are marked with an asterisk, non-significant trends are marked with a plus sign. Df = degrees of freedom; BF = Bayes factor.

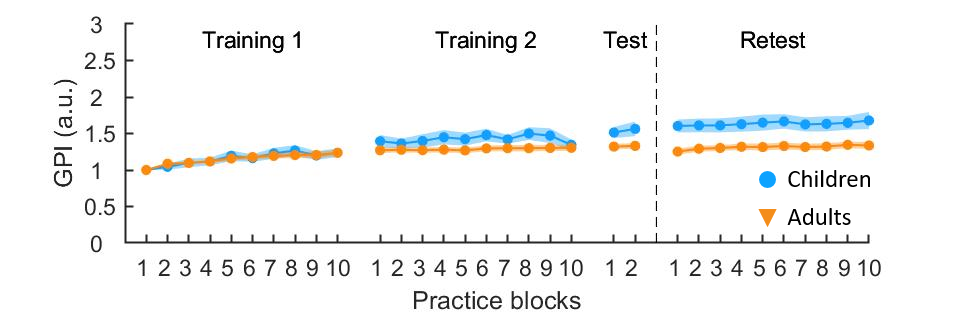

#### Figure S7. Normalized General Performance Index (GPI) across practice blocks for all FTT runs.

GPI is a single composite measure of performance that integrates performance speed and accuracy of each practice block, capturing both components of the task. Shaded regions represent the standard errors of the mean. n = 22 children and 23 adults. Statistical output on block-to-block changes in GPI is provided in Supplementary Table S8.

#### Table S8. Statistics output of block-to-block changes in the general performance index (GPI).

| **Effect** | **df** | **F** | **p** | **ƞ^2^** | **BF_10_** |
| --- | --- | --- | --- | --- | --- |
| *A. Training* |  |  |  |  |  |
| Group | 1, 42 | 0.43 | 0.518 | 0.006 | 0.511 |
| Run | 1, 42 | 110.03 | < 0.001* | 0.159 | 2.132*e^55^ |
| Block | 3.27, 137.27 | 7.40 | < 0.001* | 0.032 | 15588.04 |
| G x R | 1, 42 | 7.74 | 0.008* | 0.013 | 1243037 |
| G x B | 3.27, 137.27 | 0.42 | 0.754 | 0.002 | 0.002 |
| R x B | 5.71, 240.03 | 5.61 | < 0.001* | 0.015 | 23.993 |
| G x R x B | 5.71, 240.03 | 0.33 | 0.917 | < 0.001 | 0.006 |
| *B. Test* |  |  |  |  |  |
| Group | 1, 43 | 5.67 | 0.022* | 0.105 | 2.884 |
| Block | 1, 43 | 0.94 | 0.337 | 0.002 | 0.325 |
| G x B | 1, 43 | 0.42 | 0.522 | 0.001 | 0.366 |
| *C. Retest* |  |  |  |  |  |
| Group | 1, 43 | 9.41 | 0.004* | 0.171 | 6.634 |
| Block | 5.20, 223,69 | 3.01 | 0.011* | 0.004 | 7.882 |
| G x B | 5.20, 223.69 | 0.38 | 0.866 | < 0.001 | 0.013 |

Results from statistical analyses of normalized general performance index (GPI) for all FTT task runs. Significant values are marked with an asterisk. Df = degrees of freedom; ƞ^2^ = eta squared; G x R = group x training run interaction; G x B = group x block interaction; R x B = training run x block interaction; G x R x B = group x training run x block interaction.

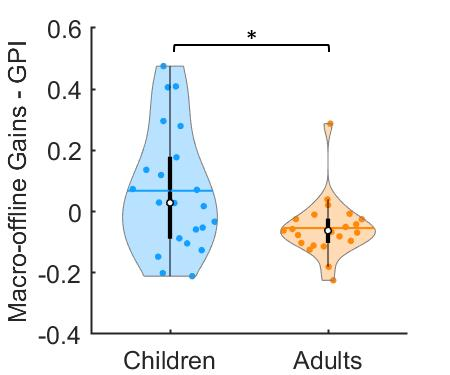

#### Figure S8. Macro-offline changes in the General Performance Index (GPI).

Positive values are indicative of performance improvements from end of training to the 5-hour delayed retest. Shaded regions represent the kernel density estimates of the data, colored circles depict individual data, open circles represent group medians, and the horizontal lines depict group means (Bechtold et al., 2021). n = 22 children and 23 adults. There was a significant group effect (t_43_ = 2.583, p = 0.007, G = 0.76, BF_10_ = 4.233), with children demonstrating greater offline performance changes in the GPI as compared to adults.

#### Table S9. Age-related changes in offline performance changes during childhood.

| **Variable** |  | **model** | **Intercept** | **C_1_** | **R^2^** | **F** | **p** |
| --- | --- | --- | --- | --- | --- | --- | --- |
| Micro-online | Training 1 | Linear | -0.322 | 1.576 | 0.039 | 0.804 | 0.381 |
|  | Training 2 | Linear | -0.255 | 1.527 | 0.058 | 1.233 | 0.280 |
| Micro-offline | Training 1 | Power | 0.219 | 0.852 | 0.010 | 0.207 | 0.654 |
| Micro-offline | Training 2 | Linear | 0.347 | -2.490 | 0.128 | 2.944 | 0.102 |
| Macro-offline | Speed | Power | 1.709 | -196.269 | 0.185 | 4.554 | 0.045* |
| Macro-offline | Accuracy | Linear | 0.009 | -0.063 | 0.052 | 1.095 | 0.308 |

Statistical output of the exploratory analyses assessing age-related changes in motor memory consolidation behaviors during childhood. Separate multiple regression analyses were conducted with age as a continuous independent variable and the respective behavioral measure as dependent measure. Five potential fit options (i.e., single exponential, double exponential, linear, quadratic and power functions) were tested. The model with the lowest Akaike Information Criterion value is listed in this table. C_n_ = parameter of regression equation; R^2^ = R-squared. Significant values are marked with an asterisk. Specifically, there was a significant, negative relationship between age and macro-offline changes in performance speed (see Figure S9 below).

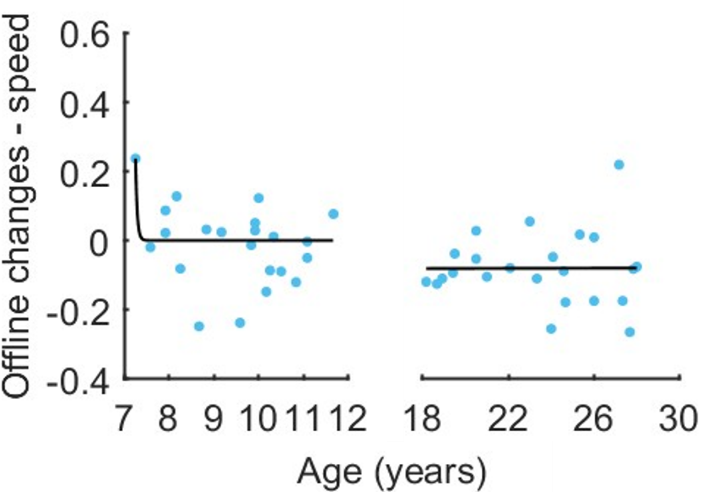

#### Figure S9. Age-related changes in macro-offline changes in speed.

Macro-offline performance changes are plotted as a function of age, separately for children and adults. Statistical parameters are provided in Table S9 above. As the significant result in children appears to be driven by the type of fit and the youngest child participant with the highest offline gain, we did not consider this result further.

### APPENDIX 4: Head motion

#### Table S10. Head motion parameters across task runs.

| **Epoch/region of interest** | | **Children** | **Adults** |
| --- | --- | --- | --- |
| Training 1 | FD (avg.) | 0.287 | 0.149 |
|  | Translation (max.) | 2.80 | 2.00 |
|  | Rotation (max.) | 0.143 | 0.023 |
| Training 2 | FD (avg.) | 0.310 | 0.140 |
|  | Translation (max) | 3.098 | 1.325 |
|  | Rotation (max.) | 0.10 | 0.021 |
| Retest | FD (avg.) | 0.244 | 0.151 |
|  | Translation (max) | 3.212 | 1.066 |
|  | Rotation (max.) | 0.063 | 0.016 |

Head motion parameters, including average framewise displacement (FD), maximum absolute value of the translation (mm) and rotations (rad) for each age group. Averages of translation and rotation maxima were computed across all three axes (x, y and z).

#### Table S11. Statistics of group x task run ANOVAs on head motion parameters.

| **Variable** |  | **df** | **F** | **p** | **ƞ^2^** | **BF_10_** |
| --- | --- | --- | --- | --- | --- | --- |
| *A. Average framewise displacement* | | | | | | |
| Group | | 1, 39 | 42.57 | < 0.001* | 0.430 | 2.64*e^5^ |
| Run | | 1.48, 57.88 | 3.64 | 0.045* | 0.028 | 0.334 |
| Group x Run | | 1.48, 57.88 | 6.11 | 0.008* | 0.046 | 4.560 |
| *B. Maximum translation* | | | | | | |
| Group | | 1, 39 | 24.43 | < 0.001* | 0.253 | 1506.065 |
| Run | | 1.61, 62.94 | 8.37 | 0.001* | 0.090 | 25.286 |
| Group x Run | | 1.61, 62.94 | 2.36 | 0.113 | 0.027 | 0.753 |
| *C. Maximum rotation* | | | | | | |
| Group | | 1, 39 | 27.70 | < 0.001* | 0.248 | 3535.543 |
| Run. | | 1.80, 70.38 | 3.65 | 0.035* | 0.048 | 0.307 |
| Group x Run | | 1.80, 70.38 | 2.61 | 0.086 | 0.035 | 0.392 |

Statistical results of the group by task run ANOVAs on head motion parameters, including framewise displacement, maximum translation and rotation. Significant values are marked with an asterisk.

### APPENDIX 5: Supplementary fMRI results

#### Table S12. Results of task-related activity during Training 1 across both age groups [task practice vs. interleaved rest].

| **Area** | **k** | **x** | **y** | **z** | **T** | **P_uncorr_** |
| --- | --- | --- | --- | --- | --- | --- |
| **a. ALL (task > rest)** |  |  |  |  |  |  |
| L Cerebellum 8 | 4857 | -20 | -60 | -48 | 16.65 | < 0.001 |
| R Cerebellum 4-5 |  | 15 | -52 | -25 | 15.01 | < 0.001 |
| L Cerebellum 4-5 |  | -5 | -55 | -15 | 14.85 | < 0.001 |
| L Supplementary Motor Area | 17606 | -5 | 2 | 58 | 15.04 | < 0.001 |
| R Postcentral |  | 40 | -22 | 52 | 14.93 | < 0.001 |
| L Precentral |  | -38 | -15 | 60 | 14.48 | < 0.001 |

Table presents clusters at p_uncorr_ < 0.001. ALL = across all participants, irrespective of age group. Voxels that were located outside of grey matter are not reported. k = cluster size, determined at p_uncorr_ < 0.001. Minimum cluster size = 5 voxels. x, y and z coordinates are specified in MNI space. Activation peaks were in the cerebellum and supplementary motor area; however, these clusters extended into pre and postcentral gyri, middle cingulate, prefrontal and parietal cortices as well as the putamen, caudate and thalamus. Results presented in this table are based on n = 21 children and 23 adults.

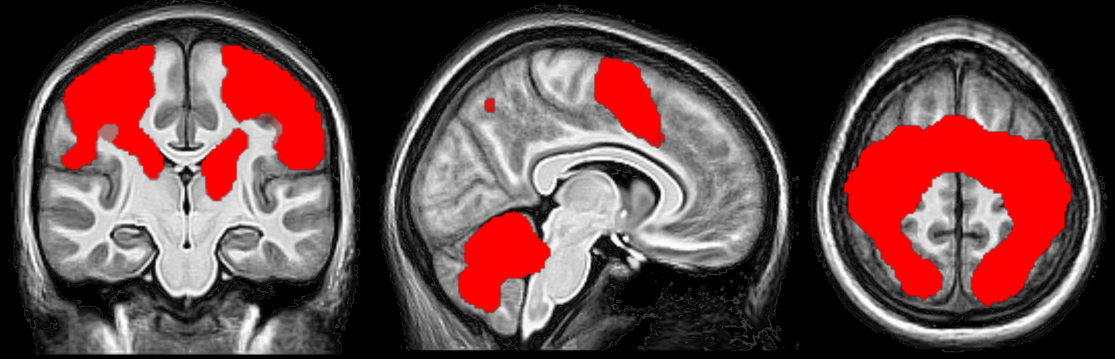

#### Figure S10. Brain regions showing significant task-related brain activation in both age groups.

Images are centered on the left supplementary motor area [-5 2 58]. Statistical images are thresholded at p_uncorrected_ < 0.001. x, y and z coordinates are specified in MNI space. In brief, and as expected, task practice recruited a large bilateral network of cortical (i.e., M1, postcentral gyrus, supplementary motor area, middle cingulate, prefrontal and parietal cortices) and subcortical (i.e., putamen, caudate, thalamus) regions, as well as the cerebellum. These results indicate that task performance elicited the expected activation in regions known to be involved in a motor sequence learning task.

#### Table S13. Within-group results of task-related activity during Training 1 [task practice vs. interleaved rest]

| **Area** | **k** | **x** | **y** | **z** | **T** | **P_uncorr_** |
| --- | --- | --- | --- | --- | --- | --- |
| **a. Adults (task > rest)** | |  |  |  |  |  |
| L Cerebellum 8 | 27690 | -20 | -60 | -48 | 19.06 | < 0.001 |
| R Cerebellum 4-5 |  | 18 | -52 | -25 | 16.36 | < 0.001 |
| **b. Children (task > rest)** |  |  |  |  |  |  |
| L Cerebellum 4-5 | 1411 | -5 | -55 | -15 | 7.11 | < 0.001 |
| R Cerebellum 4-5 |  | 10 | -52 | -22 | 6.02 | < 0.001 |
| L Cerebellum 8 |  | -18 | -60 | -45 | 6.02 | < 0.001 |
| L Supplementary motor area | 1687 | -8 | 2 | 58 | 6.75 | < 0.001 |
| L Precentral |  | -38 | -15 | 60 | 6.60 | < 0.001 |
| L Superior frontal |  | -22 | -12 | 48 | 5.38 | < 0.001 |
| R Precentral | 1105 | 38 | -22 | 55 | 6.45 | < 0.001 |
| R Cerebellum 8 | 32 | 18 | -65 | -45 | 4.24 | < 0.001 |
| **c. -Adults (rest > task)** | |  |  |  |  |  |
| R Fusiform | 20180 | 28 | -42 | -12 | 11.56 | < 0.001 |
| L Middle cingulate |  | -5 | -40 | 40 | 9.65 | < 0.001 |
| R Inferior frontal, orbital | 129 | 35 | 35 | -12 | 5.40 | < 0.001 |
| R Inferior frontal, triangular | 138 | 52 | 32 | 10 | 4.63 | < 0.001 |
| R Cerebellum crus 2 | 93 | 32 | -75 | -40 | 4.09 | < 0.001 |
| **d. -Children (rest > task)** | |  |  |  |  |  |
| R Fusiform | 20459 | 28 | -45 | -12 | 15.56 | < 0.001 |
| L Fusiform |  | -25 | -52 | -10 | 12.25 | < 0.001 |
|  |  | -28 | -32 | -22 | 10.22 | < 0.001 |
| L Medial superior frontal | 111 | -2 | 70 | 0 | 4.15 | < 0.001 |
|  |  | -5 | 70 | 15 | 3.77 | < 0.001 |
| L Superior frontal |  | -15 | 70 | -2 | 3.46 | < 0.001 |
| L Inferior frontal, triangular | 122 | -52 | 20 | 15 | 4.07 | < 0.001 |
|  |  | -50 | 30 | 12 | 3.82 | < 0.001 |
| L Anterior orbital | 57 | -28 | 35 | -15 | 4.02 | < 0.001 |
| R Inferior frontal, triangular | 45 | 52 | 42 | 12 | 3.90 | < 0.001 |
| R Supramarginal | 64 | 62 | -32 | 42 | 3.77 | < 0.001 |
| L Medial superior frontal | 16 | 0 | 38 | 30 | 3.55 | < 0.001 |
|  | 19 | 0 | 30 | 48 | 3.52 | 0.001 |
| R Supramarginal | 6 | 45 | -28 | 25 | 3.45 | 0.001 |
|  |  | 55 | -48 | 35 | 3.34 | 0.001 |
| L Postcentral | 6 | -60 | -5 | 30 | 3.38 | 0.001 |

Table presents clusters at p_uncorr_ < 0.001. Voxels that were located outside of grey matter are not reported. k = cluster size, determined at p_uncorr_ < 0.001. Minimum cluster size = 5 voxels. x, y and z coordinates are specified in MNI space. Results presented in this table are based on n = 21 children and 23 adults. Results of the between-group contrasts are depicted in Table 2a.

#### Table S14. Uncorrected results of task-related activity during early learning (task practice vs. rest), with average transition time across practice blocks as a covariate.

| **Area** | **k** | | **x** | **y** | **z** | **T** | **P_uncorr_** |
| --- | --- | --- | --- | --- | --- | --- | --- |
| **a. Children - adults** |  | |  |  |  |  |  |
| No suprathreshold voxels |  | |  |  |  |  |  |
| **a1. Children (task > rest)** |  | |  |  |  |  |  |
| Vermis 4-5 | 597 | | -2 | -60 | -15 | 6.04 | < 0.001 |
| L Precentral | 529 | | -38 | -18 | 60 | 6.01 | < 0.001 |
| R Precentral | 336 | | 35 | -22 | 55 | 5.29 | < 0.001 |
| L Cerebellum 8 | 27 | | -20 | -60 | -48 | 4.73 | < 0.001 |
| L SMA | 99 | | -8 | 2 | 58 | 4.55 | < 0.001 |
| R Cerebellum 8 | 9 | | 30 | -42 | -48 | 3.70 | < 0.001 |
| **a2. -Adults (rest > task)** |  |  | |  |  |  |  |
| R Fusiform | 369 | 28 | | -42 | -12 | 6.74 | < 0.001 |
| L Fusiform | 242 | -25 | | -48 | -10 | 4.74 | < 0.001 |
| L Middle cingulate | 475 | -8 | | -38 | 45 | 4.98 | < 0.001 |
| R Precuneus |  | 10 | | -38 | 52 | 4.29 | < 0.001 |
| L Superior anterior cingulate | 315 | 0 | | 28 | 18 | 4.67 | < 0.001 |
| L Inferior parietal | 365 | -55 | | -55 | 42 | 4.60 | < 0.001 |
| L Angular |  | -55 | | -68 | 32 | 4.47 | < 0.001 |
| L Supramarginal |  | -62 | | -42 | 40 | 4.08 | < 0.001 |
| L Medial superior frontal | 223 | -8 | | 32 | 58 | 4.49 | < 0.001 |
| L Superior frontal | 65 | -18 | | 45 | 42 | 3.81 | < 0.001 |
| L Amygdala | 58 | -30 | | 2 | -18 | 3.79 | < 0.001 |
| R Medial superior frontal | 13 | 5 | | 65 | 12 | 3.43 | 0.001 |
| R Middle cingulate | 6 | 5 | | -20 | 40 | 3.41 | 0.001 |
| **b. Adults - children** |  | |  |  |  |  |  |
| L Fusiform | 4572 | | -32 | -58 | -18 | 6.87 | < 0.001 |
| L SMA | 161 | | -12 | -12 | 72 | 6.07 | < 0.001 |
|  |  | | -15 | -2 | 65 | 4.08 | < 0.001 |
| L Middle frontal |  | | -25 | -2 | 55 | 3.84 | < 0.001 |
| R Postcentral | 3130 | | 52 | -15 | 40 | 5.81 | < 0.001 |
| R Superior parietal |  | | 18 | -62 | 50 | 5.46 | < 0.001 |
| R Inferior parietal |  | | 32 | -48 | 52 | 5.31 | < 0.001 |
| R Cerebellum 6 | 1446 | | 28 | -58 | -22 | 5.80 | < 0.001 |
| L Precentral | 351 | | -50 | 0 | 42 | 5.05 | < 0.001 |
|  |  | | -58 | 2 | 30 | 4.73 | < 0.001 |
| L Putamen | 248 | | -30 | -20 | -2 | 4.59 | < 0.001 |
| L Fusiform | 19 | | -25 | -32 | -25 | 4.41 | < 0.001 |
| R Cerebellum 8 | 59 | | 18 | -70 | -48 | 4.31 | < 0.001 |
| L Cerebellum 8 | 47 | | -18 | -60 | -48 | 4.23 | < 0.001 |
| L SMA | 213 | | 0 | 0 | 60 | 4.20 | < 0.001 |
| L Middle cingulate |  | | 0 | 5 | 42 | 3.43 | 0.001 |
| R Superior frontal | 15 | | 18 | -10 | 75 | 4.09 | < 0.001 |
| R Fusiform | 12 | | 32 | -32 | -25 | 3.87 | 0.001 |
| **b1. -Children (rest > task)** |  | |  |  |  |  |  |
| R Fusiform | 25620 | | 30 | -42 | -12 | 15.15 | < 0.001 |
| L Fusiform |  | | -25 | -48 | -10 | 12.30 | < 0.001 |
| R Postcentral | 473 | | 40 | -12 | 32 | 6.03 | < 0.001 |
| R Inferior frontal, triangular | 367 | | 52 | 40 | 10 | 4.64 | < 0.001 |
| R Middle frontal |  | | 48 | 50 | 12 | 4.22 | < 0.001 |
| R Inferior parietal | 120 | | 52 | -50 | 38 | 4.38 | < 0.001 |
| R Supramarginal | 71 | | 60 | -35 | 40 | 3.99 | < 0.001 |
| L Cerebellum crus 2 | 34 | | -12 | -80 | -45 | 3.92 | < 0.001 |
| R Middle frontal | 10 | | 38 | 15 | 42 | 3.47 | 0.001 |
| R Inferior frontal, opercular | 5 | | 40 | 12 | 30 | 3.39 | 0.001 |
| **b2. Adults (task > rest)** |  |  | |  |  |  |  |
| L Cerebellum 8 | 3078 | -18 | | -60 | -48 | 8.51 | < 0.001 |
| L Cerebellum 4-5 |  | -5 | | -58 | -15 | 7.70 | < 0.001 |
| R Cerebellum 6 |  | 18 | | -55 | -25 | 7.40 | < 0.001 |
| L SMA | 10936 | -2 | | 0 | 60 | 7.98 | < 0.001 |
| L Superior parietal |  | -28 | | -55 | 50 | 7.61 | < 0.001 |
| L Precentral |  | -15 | | -12 | 72 | 7.54 | < 0.001 |
| R Thalamus | 21 | 18 | | -10 | 2 | 4.00 | < 0.001 |

Table presents clusters at p_uncorr_ < 0.001. Voxels that were located outside of grey matter are not reported. k = cluster size, determined at p_uncorr_ < 0.001. Minimum cluster size = 5 voxels. x, y and z coordinates are specified in MNI space. SMA = Supplementary Motor Area. Results presented in this table are based on *n* = 21 children and 23 adults. In brief, the [adults – children] contrast remained similar, with significant age group differences in frontal, parietal and cerebellar regions as well as the putamen and fusiform gyrus. Accordingly, activation differences between children and adults in Training 1 do not appear to be driven by differences in performance speed. In contrast, the inclusion of speed as a covariate caused the disappearance of all regions in the [children – adults] contrast, including the left superior frontal left angular, left middle frontal and left posterior cingulate gyri. Accordingly, the default mode network (DMN), did not show a smaller deactivation in children than adults anymore when performance speed was included as a covariate.

#### Table S15. Regression analyses between brain responses during training 1 and micro-online performance changes [micro-online changes x training 1].

| **Area** | **k** | **x** | **y** | **z** | **T** | **p_uncorr_** |
| --- | --- | --- | --- | --- | --- | --- |
| **a. Children - adults** |  |  |  |  |  |  |
| No suprathreshold voxels |  |  |  |  |  |  |
| **a1. Children** |  |  |  |  |  |  |
| No suprathreshold voxels |  |  |  |  |  |  |
| **a2. -Adults** |  |  |  |  |  |  |
| No suprathreshold voxels |  |  |  |  |  |  |
| **b. Adults - children** |  |  |  |  |  |  |
| No suprathreshold voxels |  |  |  |  |  |  |
| **b1. -Children** |  |  |  |  |  |  |
| No suprathreshold voxels |  |  |  |  |  |  |
| **b2. Adults** |  |  |  |  |  |  |
| R Postcentral | 34 | 15 | -28 | 62 | 3.74 | < 0.001 |
| **c. ALL** |  |  |  |  |  |  |
| No suprathreshold voxels |  |  |  |  |  |  |
| **d. -ALL** |  |  |  |  |  |  |
| No suprathreshold voxels |  |  |  |  |  |  |

Table presents clusters at p_uncorr_ < 0.001. Voxels that were located outside of grey matter are not reported. k = cluster size. Minimum cluster size = 5 voxels. x, y and z coordinates are specified in MNI space. Results presented in this table are based on n = 21 children and 23 adults. ALL = across all participants, irrespective of age group.

#### Table S16. Regression analyses between brain responses during training 1 and micro-offline performance changes [micro-offline changes x training 1].

| **Area** | **k** | **x** | **y** | **z** | **T** | **p_uncorr_** |
| --- | --- | --- | --- | --- | --- | --- |
| **a. Children - adults** |  |  |  |  |  |  |
| No suprathreshold voxels |  |  |  |  |  |  |
| **a1. Children** |  |  |  |  |  |  |
| No suprathreshold voxels |  |  |  |  |  |  |
| **a2. -Adults** |  |  |  |  |  |  |
| R Postcentral | 14 | 18 | -30 | 62 | 3.57 | < 0.001 |
| **b. Adults - children** |  |  |  |  |  |  |
| No suprathreshold voxels |  |  |  |  |  |  |
| **b1. -Children** |  |  |  |  |  |  |
| No suprathreshold voxels |  |  |  |  |  |  |
| **b2. Adults** |  |  |  |  |  |  |
| No suprathreshold voxels |  |  |  |  |  |  |
| **c. ALL** |  |  |  |  |  |  |
| No suprathreshold voxels |  |  |  |  |  |  |
| **d. -ALL** |  |  |  |  |  |  |
| No suprathreshold voxels |  |  |  |  |  |  |

Table presents clusters at p_uncorr_ < 0.001. Voxels that were located outside of grey matter are not reported. k = cluster size. Minimum cluster size = 5 voxels. x, y and z coordinates are specified in MNI space. Results presented in this table are based on n = 21 children and 23 adults. ALL = across all participants, irrespective of age group.

#### Table S17. Additional results of the regression analysis of task activity during training 1 with the macro-offline performance changes.

| **Area** | **k** | **x** | **y** | **z** | **T** | **p_uncorr_** |
| --- | --- | --- | --- | --- | --- | --- |

| **a. Adults** |  |  |  |  |  |  |
| --- | --- | --- | --- | --- | --- | --- |
| No suprathreshold clusters |  |  |  |  |  |  |
| **b. Children** |  |  |  |  |  |  |
| L Middle cingulate | 15367 | -12 | -15 | 35 | 6.23 | < 0.001 |
| R Middle frontal | 250 | 52 | 45 | 5 | 6.00 | < 0.001 |
| R Inferior frontal, triangular |  | 40 | 28 | 8 | 3.36 | 0.001 |
| L Precentral | 2074 | -50 | 5 | 38 | 4.95 | < 0.001 |
| L Middle frontal | 305 | -48 | 42 | 22 | 4.95 | < 0.001 |
|  |  | -40 | 45 | 32 | 4.67 | < 0.001 |
|  |  | -30 | 55 | 30 | 4.52 | < 0.001 |
| R Middle frontal | 163 | 55 | 15 | 40 | 4.20 | < 0.001 |
|  |  | 38 | 28 | 50 | 3.78 | < 0.001 |
| R Postcentral | 328 | 45 | -42 | 60 | 4.05 | < 0.001 |
| R Inferior parietal |  | 48 | -50 | 40 | 3.92 | < 0.001 |
| R Supramarginal |  | 58 | -32 | 50 | 3.57 | < 0.001 |
| L SMA | 49 | -5 | 5 | 60 | 3.93 | < 0.001 |
| L Inferior parietal | 34 | -60 | -38 | 50 | 3.77 | < 0.001 |
|  |  | -58 | -52 | 45 | 3.69 | < 0.001 |
| R Cerebellum 8 | 9 | 35 | -52 | -52 | 3.72 | < 0.001 |
| R Superior frontal | 9 | 22 | 50 | 42 | 3.71 | < 0.001 |
| R Amygdala | 5 | 20 | -2 | -12 | 3.66 | < 0.001 |
| L Lateral orbitofrontal | 40 | -40 | 45 | -12 | 3.64 | < 0.001 |
| L Inferior frontal, orbital |  | -50 | 45 | -5 | 3.53 | 0.001 |
| R SMA | 6 | 5 | -8 | 72 | 3.58 | < 0.001 |
| R Middle frontal | 16 | 40 | 8 | 55 | 3.54 | 0.001 |
| L Cerebellum 7 | 12 | -30 | -70 | -48 | 3.51 | 0.001 |
| R Posterior orbital | 9 | 28 | 28 | -20 | 3.48 | 0.001 |
| R Superior parietal | 8 | 15 | -65 | 58 | 3.46 | 0.001 |
| L Inferior parietal | 28 | -48 | -45 | 38 | 3.45 | 0.001 |
|  |  | -42 | -35 | 35 | 3.39 | 0.001 |
| R Inferior frontal, orbital | 6 | 35 | 38 | -12 | 3.43 | 0.001 |
| **c. -Adults** |  |  |  |  |  |  |
| R Postcentral | 115 | 25 | -38 | 62 | 4.43 | < 0.001 |
| L Postcentral | 5 | -25 | -38 | 58 | 3.50 | 0.001 |
| **d. -Children** |  |  |  |  |  |  |
| No suprathreshold clusters |  |  |  |  |  |  |

| **e. ALL** |  |  |  |  |  |  |
| --- | --- | --- | --- | --- | --- | --- |
| L Precentral | 59 | -52 | 0 | 48 | 4.33 | < 0.001 |
| L Inferior frontal, opercular | 87 | -52 | 10 | 0 | 4.14 | < 0.001 |
| R SMA | 8 | 10 | 12 | 68 | 4.11 | < 0.001 |
| L Middle frontal | 13 | -48 | 42 | 22 | 4.01 | < 0.001 |
| R Posterior cingulate | 46 | 12 | -40 | 8 | 4.00 | < 0.001 |
| R Cerebellum 8 | 12 | 35 | -52 | -52 | 3.95 | < 0.001 |
| L Middle frontal | 7 | -30 | 55 | 30 | 3.94 | < 0.001 |
| R Angular | 155 | 45 | -58 | 40 | 3.91 | < 0.001 |
|  |  | 32 | -62 | 45 | 3.73 | < 0.001 |
| L Cerebellum 6 | 54 | -22 | -72 | -25 | 3.85 | < 0.001 |
|  |  | -8 | -80 | -18 | 3.67 | < 0.001 |
| L Paracentral | 5 | -12 | -15 | 75 | 3.84 | < 0.001 |
| L Paracentral | 30 | -10 | -25 | 62 | 3.70 | < 0.001 |
| R Cerebellum crus 1 | 26 | 40 | -70 | -22 | 3.63 | < 0.001 |
| R Cerebellum 6 | 17 | 25 | -62 | -20 | 3.41 | 0.001 |
| **f. -ALL** |  |  |  |  |  |  |
| No suprathreshold clusters | |  |  |  |  |  |

Table presents clusters at p_uncorr_ < 0.001. Voxels that were located outside of grey matter are not reported. k = cluster size, determined at p_uncorr_ < 0.001. Minimum cluster size = 5 voxels. x, y and z coordinates are specified in MNI space. Results presented in this table are based on n = 21 children and 23 adults. Results of the between-group contrasts are depicted in Table 2b. ALL = across all participants, irrespective of age group.

#### Table S18. Task-related activity during training 2 [task practice vs. interleaved rest].

| **Area** | **k** | **x** | **y** | **z** | **T** | **p_uncorr_** |
| --- | --- | --- | --- | --- | --- | --- |
| **a. Children - adults** |  |  |  |  |  |  |
| L Medial superior frontal | 403 | -10 | 52 | 30 | 4.40 | < 0.001 |
| L Superior frontal |  | -12 | 32 | 52 | 3.72 | < 0.001 |
| R Medial superior frontal |  | 8 | 52 | 32 | 3.61 | < 0.001 |
| L Posterior cingulate | 45 | -8 | -52 | 30 | 3.86 | < 0.001 |
| L Angular | 71 | -48 | -68 | 30 | 3.85 | < 0.001 |
| L Inferior frontal, triangular | 6 | -55 | 25 | 22 | 3.83 | < 0.001 |
| L Medial frontal, orbital | 6 | -2 | 60 | -10 | 3.43 | 0.001 |
| **a1. Children (task > retest)** |  |  |  |  |  |  |
| Vermis 4_5 | 1028 | -2 | -52 | -12 | 8.25 | < 0.001 |
| R Precentral | 540 | 38 | -20 | 62 | 7.69 | < 0.001 |
| L Precentral | 490 | -38 | -20 | 60 | 7.02 | < 0.001 |
| L Superior frontal |  | -22 | -8 | 50 | 4.62 | < 0.001 |
| L SMA | 217 | -5 | 5 | 55 | 5.82 | < 0.001 |
| R Cerebellum 8 | 7 | 30 | -42 | -48 | 4.04 | < 0.001 |
| R Cerebellum 6 | 11 | 38 | -42 | -35 | 3.82 | < 0.001 |
| L Cerebellum 8 | 5 | -12 | -62 | -45 | 3.64 | < 0.001 |
| L Inferior parietal | 21 | -28 | -48 | 45 | 3.58 | < 0.001 |
| **a2. -Adults (rest > task)** |  |  |  |  |  |  |
| L Inferior frontal, triangular | 6020 | -55 | 25 | 22 | 8.02 | < 0.001 |
| L Superior frontal |  | -20 | 22 | 60 | 6.89 | < 0.001 |
| L Medial superior frontal |  | -5 | 35 | 42 | 6.47 | < 0.001 |
| L Middle cingulate | 8620 | -8 | -40 | 42 | 7.65 | < 0.001 |
| L Fusiform |  | -28 | -40 | -12 | 7.19 | < 0.001 |
| R Inferior frontal, orbital | 81 | 28 | 32 | -12 | 4.74 | < 0.001 |
| R Inferior frontal, triangular | 12 | 52 | 30 | 20 | 3.53 | 0.001 |
| **b. Adults - children** |  |  |  |  |  |  |
| L Cerebellum 8 | 549 | -20 | -65 | -48 | 7.87 | < 0.001 |
| L Precentral | 14932 | -15 | -12 | 80 | 7.81 | < 0.001 |
| R SMA |  | 5 | 0 | 62 | 7.41 | < 0.001 |
| R Postcentral |  | 40 | -40 | 60 | 7.32 | < 0.001 |
| R Cerebellum 6 | 4518 | 25 | -55 | -22 | 7.44 | < 0.001 |
| L Cerebellum 6 |  | -28 | -55 | -25 | 6.34 | < 0.001 |
| R Cerebellum 8 | 432 | 22 | -55 | -50 | 6.41 | < 0.001 |
| **b1. -Children (rest > task)** |  |  |  |  |  |  |
| Fusiform | 15643 | 30 | -38 | -20 | 10.66 | < 0.001 |
| L Inferior parietal | 722 | -55 | -50 | 42 | 6.51 | < 0.001 |
|  |  | -60 | -58 | 50 | 5.21 | < 0.001 |
| R Superior frontal | 105 | 18 | 70 | -8 | 5.09 | < 0.001 |
| R Anterior orbital |  | 12 | 65 | -15 | 4.39 | < 0.001 |
| R Superior frontal |  | 25 | 68 | 12 | 3.89 | < 0.001 |
| R Supramarginal | 1034 | 62 | -35 | 45 | 5.03 | < 0.001 |
|  |  | 55 | -45 | 38 | 5.02 | < 0.001 |
|  |  | 65 | -25 | 42 | 4.80 | < 0.001 |
| R Inferior frontal, triangular | 297 | 50 | 40 | 12 | 4.62 | < 0.001 |
| R Middle frontal |  | 42 | 12 | 38 | 3.55 | 0.001 |
| R Inferior frontal, triangular |  | 45 | 18 | 25 | 3.45 | 0.001 |
| L Anterior orbital | 52 | -22 | 35 | -18 | 4.56 | < 0.001 |
| L Medial superior frontal | 286 | -2 | 28 | 48 | 4.30 | < 0.001 |
| L Anterior cingulate |  | 0 | 35 | 30 | 3.97 | < 0.001 |
| L Inferior frontal, triangular | 121 | -52 | 35 | 15 | 4.24 | < 0.001 |
| R Medial orbital | 38 | 20 | 35 | -18 | 4.23 | < 0.001 |
| L Anterior orbital | 8 | -28 | 55 | -15 | 3.92 | < 0.001 |
| **b2. Adults (task > rest)** |  |  |  |  |  |  |
| R Precentral | 18262 | 40 | -22 | 60 | 15.39 | < 0.001 |
| L Precentral |  | -38 | -18 | 60 | 15.14 | < 0.001 |
| R SMA |  | 2 | 0 | 60 | 14.62 | < 0.001 |
| Vermis 4_5 | 5366 | -2 | -55 | -12 | 13.94 | < 0.001 |
| R Cerebellum 6 |  | 20 | -52 | -25 | 14.50 | < 0.001 |
| R Cerebellum 4_5 |  | 10 | -50 | 018 | 14.12 | < 0.001 |

Table presents clusters at p_uncorr_ < 0.001. Voxels that were located outside of grey matter are not reported. k = cluster size. Minimum cluster size = 5 voxels. x, y and z coordinates are specified in MNI space. SMA = Supplementary Motor Area. Results presented in this table are based on n = 19 children and 23 adults.

#### Table S19. Regression analyses between brain responses during training 2 and micro-online performance changes [micro-online changes x training 2].

| **Area** | **k** | **x** | **y** | **z** | **T** | **p_uncorr_** |
| --- | --- | --- | --- | --- | --- | --- |
| **a. Children - adults** |  |  |  |  |  |  |
| No suprathreshold voxels |  |  |  |  |  |  |
| **a1. Children** |  |  |  |  |  |  |
| No suprathreshold voxels |  |  |  |  |  |  |
| **a2. -Adults** |  |  |  |  |  |  |
| No suprathreshold voxels |  |  |  |  |  |  |
| **b. Adults - children** |  |  |  |  |  |  |
| L Precentral | 11 | -25 | -5 | 48 | 3.54 | 0.001 |
| **b1. -Children** |  |  |  |  |  |  |
| No suprathreshold voxels |  |  |  |  |  |  |
| **b2. Adults** |  |  |  |  |  |  |
| L Precentral | 21 | -28 | -5 | 48 | 3.73 | < 0.001 |
| **c. ALL** |  |  |  |  |  |  |
| No suprathreshold voxels |  |  |  |  |  |  |
| **d. -ALL** |  |  |  |  |  |  |
| No suprathreshold voxels |  |  |  |  |  |  |

Table presents clusters at p_uncorr_ < 0.001. Voxels that were located outside of grey matter are not reported. k = cluster size. Minimum cluster size = 5 voxels. x, y and z coordinates are specified in MNI space. Results presented in this table are based on n = 19 children and 23 adults. ALL = across all participants, irrespective of age group.

#### Table S20. Regression analyses between brain responses during training 2 and micro-offline performance changes [micro-offline changes x training 2].

| **Area** | **k** | **x** | **y** | **z** | **T** | **p_uncorr_** |
| --- | --- | --- | --- | --- | --- | --- |
| **a. Children - adults** |  |  |  |  |  |  |
| L Precentral | 12 | -25 | -5 | 48 | 3.58 | < 0.001 |
| **a1. Children** |  |  |  |  |  |  |
| No suprathreshold voxels |  |  |  |  |  |  |
| **a2. -Adults** |  |  |  |  |  |  |
| L Precentral | 18 | -28 | -5 | 48 | 3.70 | < 0.001 |
| **b. Adults - children** |  |  |  |  |  |  |
| No suprathreshold voxels |  |  |  |  |  |  |
| **b1. -Children** |  |  |  |  |  |  |
| No suprathreshold voxels |  |  |  |  |  |  |
| **b2. Adults** |  |  |  |  |  |  |
| No suprathreshold voxels |  |  |  |  |  |  |
| **c. ALL** |  |  |  |  |  |  |
| No suprathreshold voxels |  |  |  |  |  |  |
| **d. -ALL** |  |  |  |  |  |  |
| No suprathreshold voxels |  |  |  |  |  |  |

Table presents clusters at p_uncorr_ < 0.001. Voxels that were located outside of grey matter are not reported. k = cluster size. Minimum cluster size = 5 voxels. x, y and z coordinates are specified in MNI space. Results presented in this table are based on n = 19 children and 23 adults. ALL = across all participants, irrespective of age group.

#### Table S21. Regression analyses between brain responses during training 2 and macro-offline performance changes [macro-offline changes x training 2].

| **Area** | **k** | **x** | **y** | **z** | **T** | **p_uncorr_** |
| --- | --- | --- | --- | --- | --- | --- |
| **a. Children - adults** |  |  |  |  |  |  |
| R Superior frontal | 124 | 30 | 48 | 18 | 3.90 | < 0.001 |
| L Middle frontal | 18 | -38 | 35 | 22 | 3.70 | < 0.001 |
| R Middle cingulate | 44 | 15 | 22 | 30 | 3.68 | < 0.001 |
|  |  | 12 | 15 | 42 | 3.61 | < 0.001 |
| R Cerebellum 6 | 22 | 8 | -72 | -25 | 3.64 | < 0.001 |
| R Inferior frontal, triangular | 10 | 50 | 40 | 5 | 3.57 | < 0.001 |
| **a1. Children** |  |  |  |  |  |  |
| L Inferior frontal, triangular | 22 | -35 | 20 | 18 | 3.55 | 0.001 |
| L Angular | 5 | -48 | -62 | 52 | 3.50 | 0.001 |
| **a2. -Adults** |  |  |  |  |  |  |
| R Postcentral | 62 | 18 | -40 | 62 | 3.74 | < 0.001 |
| L Cerebellum 8 | 32 | -28 | -60 | -42 | 3.70 | < 0.001 |
| L Postcentral | 17 | -22 | -38 | 52 | 3.69 | < 0.001 |
| R Precuneus | 13 | 15 | -55 | 55 | 3.62 | < 0.001 |
| **b. Adults - children** |  |  |  |  |  |  |
| No suprathreshold clusters |  |  |  |  |  |  |
| **b1. -Children** |  |  |  |  |  |  |
| No suprathreshold clusters |  |  |  |  |  |  |
| **b2. Adults** |  |  |  |  |  |  |
| No suprathreshold clusters |  |  |  |  |  |  |
| **c. ALL** |  |  |  |  |  |  |
| No suprathreshold clusters |  |  |  |  |  |  |
| **d. -ALL** |  |  |  |  |  |  |
| L Cerebellum 8 | 5 | -25 | -60 | -45 | 3.50 | 0.001 |

Table presents clusters at p_uncorr_ < 0.001. Voxels that were located outside of grey matter are not reported. k = cluster size. Minimum cluster size = 5 voxels. x, y and z coordinates are specified in MNI space. Results presented in this table are based on n = 19 children and 23 adults. ALL = across all participants, irrespective of age group.

#### Table S22. Changes in activity from training 1 to training 2 [Training 2 – Training 1].

| **Area** | **k** | **x** | **y** | **z** | **T** | **p_uncorr_** |
| --- | --- | --- | --- | --- | --- | --- |
| **a. Children - adults** |  |  |  |  |  |  |
| No suprathreshold clusters |  |  |  |  |  |  |
| **a1. Children** |  |  |  |  |  |  |
| No suprathreshold clusters |  |  |  |  |  |  |
| **a2. -Adults** |  |  |  |  |  |  |
| R Superior frontal | 19 | 30 | 12 | 62 | 3.72 | < 0.001 |
| L Precentral | 11 | -40 | 8 | 32 | 3.61 | < 0.001 |
| L Inferior frontal, triangular | 15 | -35 | 18 | 30 | 3.58 | < 0.001 |
| R Superior parietal | 12 | -22 | -78 | 52 | 3.58 | < 0.001 |
| **b. Adults - children** |  |  |  |  |  |  |
| R Anterior orbital | 16 | 30 | 48 | -18 | 3.86 | < 0.001 |
| L Superior frontal | 24 | -28 | 55 | -8 | 3.81 | < 0.001 |
| **b1. -Children** |  |  |  |  |  |  |
| R Anterior orbital | 35 | 30 | 50 | -18 | 3.88 | < 0.001 |
|  |  | 22 | 45 | -12 | 3.54 | 0.001 |
| L Middle frontal | 17 | -30 | 52 | -8 | 3.74 | < 0.001 |
| **b2. Adults** |  |  |  |  |  |  |
| L Paracentral | 201 | -12 | -20 | 72 | 4.61 | < 0.001 |
| L SMA |  | -10 | -10 | 72 | 4.28 | < 0.001 |
| R SMA |  | 5 | -8 | 72 | 4.24 | < 0.001 |
| L Precentral | 117 | -35 | -20 | 65 | 3.96 | < 0.001 |
| R Putamen | 5 | 20 | 0 | 10 | 3.45 | 0.001 |
| **c. ALL** |  |  |  |  |  |  |
| No suprathreshold clusters |  |  |  |  |  |  |
| **d. -ALL** |  |  |  |  |  |  |
| L Superior parietal | 64 | -25 | -75 | 50 | 4.29 | < 0.001 |
| L Precentral | 12 | -38 | 5 | 32 | 3.57 | < 0.001 |
| R Inferior parietal | 74 | 40 | -52 | 48 | 3.56 | < 0.001 |
| R Superior parietal | 5 | 35 | -72 | 52 | 3.39 | 0.001 |

Table presents clusters at p_uncorr_ < 0.001. Voxels that were located outside of grey matter are not reported. k = cluster size. Minimum cluster size = 5 voxels. x, y and z coordinates are specified in MNI space. SMA = Supplementary Motor Area. Results presented in this table are based on n = 19 children and 23 adults. ALL = across all participants, irrespective of age group.

#### Table S23. Regression analyses between activity changes from training 1 to training 2 and macro-offline performance changes [macro-offline changes x (training 2 – training 1)].

| **Area** | **k** | **x** | **y** | **z** | **T** | **p_uncorr_** |
| --- | --- | --- | --- | --- | --- | --- |
| **a. Children - adults** |  |  |  |  |  |  |
| No suprathreshold clusters |  |  |  |  |  |  |
| **a1. Children** |  |  |  |  |  |  |
| No suprathreshold clusters |  |  |  |  |  |  |
| **a2. -Adults** |  |  |  |  |  |  |
| R Precentral | 5 | 35 | 0 | 42 | 3.45 | 0.001 |
| **b. Adults - children** |  |  |  |  |  |  |
| L Inferior frontal, triangular | 58 | -55 | 30 | 0 | 4.14 | < 0.001 |
| L Medial superior frontal | 11 | -10 | 32 | 60 | 3.78 | < 0.001 |
|  |  | -10 | 40 | 55 | 3.50 | 0.001 |
| L Medial superior frontal | 49 | -5 | 52 | 45 | 3.62 | < 0.001 |
| R Medial superior frontal |  | 2 | 55 | 40 | 3.54 | 0.001 |
| L Superior frontal |  | -12 | 58 | 35 | 3.50 | 0.001 |
| L Medial superior frontal | 13 | 2 | 68 | 20 | 3.61 | < 0.001 |
| L Postcentral | 5 | -22 | -30 | 75 | 3.57 | < 0.001 |
| L Medial frontal, orbital | 8 | -2 | 58 | -8 | 3.43 | 0.001 |
| L Medial superior frontal | 21 | 0 | 60 | 10 | 3.42 | 0.001 |
| **b1. -Children** |  |  |  |  |  |  |
| R Cerebellum 8 | 75 | 18 | -52 | -55 | 4.33 | < 0.001 |
|  |  | 35 | -55 | -52 | 3.64 | < 0.001 |
| L Middle cingulate | 68 | -15 | 20 | 38 | 4.26 | < 0.001 |
| R SMA | 310 | 2 | -20 | 60 | 4.24 | < 0.001 |
| R Paracentral |  | 8 | -28 | 65 | 4.17 | < 0.001 |
| L Postcentral |  | -22 | -32 | 75 | 3.77 | < 0.001 |
| L Precentral | 28 | -55 | 2 | 45 | 4.04 | < 0.001 |
| R Inferior frontal, opercular | 30 | 50 | 8 | 25 | 3.74 | < 0.001 |
| R Precentral |  | 42 | 0 | 28 | 3.36 | 0.001 |
| L Superior parietal | 22 | -28 | -68 | 60 | 3.72 | < 0.001 |
| R Thalamus | 9 | 5 | -10 | 18 | 3.54 | 0.001 |
| Vermis 3 | 16 | -2 | -45 | -10 | 3.53 | 0.001 |
| L Inferior parietal | 67 | -38 | -32 | 35 | 3.50 | 0.001 |
|  |  | -48 | -30 | 58 | 3.98 | 0.001 |
| L Cerebellum 8 | 13 | -25 | -62 | -48 | 3.50 | 0.001 |
| **b2. Adults** |  |  |  |  |  |  |
| L Medial superior frontal | 5 | -10 | 32 | 60 | 3.83 | < 0.001 |
| **c. ALL** |  |  |  |  |  |  |
| No suprathreshold clusters |  |  |  |  |  |  |
| **d. -ALL** |  |  |  |  |  |  |
| L Inferior parietal | 1332 | -38 | -32 | 30 | 4.78 | < 0.001 |
| L Postcentral |  | -42 | -25 | 42 | 4.17 | < 0.001 |
| R SMA | 180 | 8 | -18 | 62 | 4.53 | < 0.001 |
| L Paracentral |  | -10 | -28 | 65 | 4.24 | < 0.001 |
| L Cerebellum 8 | 201 | -28 | -62 | -45 | 4.52 | < 0.001 |
| R Cerebellum 8 | 70 | 32 | -55 | -52 | 4.24 | < 0.001 |
|  |  | 20 | -52 | -52 | 3.80 | < 0.001 |
|  |  | 20 | -62 | -48 | 3.42 | 0.001 |
| R Precentral | 76 | 38 | 0 | 45 | 3.96 | < 0.001 |
| R Cerebellum 6 | 105 | 15 | -62 | -25 | 3.93 | < 0.001 |
| R Cerebellum 4_5 |  | 18 | -45 | -28 | 3.65 | < 0.001 |
| R Inferior parietal | 157 | 28 | -48 | 50 | 3.92 | < 0.001 |
| R Inferior frontal, opercular | 41 | 52 | 8 | 22 | 3.72 | < 0.001 |
| R Precentral |  | 40 | 0 | 28 | 3.59 | < 0.001 |
| R Inferior frontal |  |  |  |  |  |  |

Table presents clusters at p_uncorr_ < 0.001. Voxels that were located outside of grey matter are not reported. k = cluster size. Minimum cluster size = 5 voxels. x, y and z coordinates are specified in MNI space. SMA = Supplementary Motor Area. Results presented in this table are based on n = 19 children and 23 adults. ALL = across all participants, irrespective of age group.

#### Table S24. Task-related activity during Retest [task practice vs. interleaved rest].

| **Area** | **k** | **x** | **y** | **z** | **T** | **p_uncorr_** |
| --- | --- | --- | --- | --- | --- | --- |
| **a. Children - adults** |  |  |  |  |  |  |
| R Medial superior frontal | 82 | 5 | 65 | 30 | 3.85 | < 0.001 |
| L Medial superior frontal |  | -10 | 55 | 30 | 3.81 | < 0.001 |
| L Angular | 31 | -45 | -75 | 40 | 3.59 | < 0.001 |
| **a1. Children (task > retest)** |  |  |  |  |  |  |
| R Precentral | 2670 | 38 | -22 | 58 | 9.04 | < 0.001 |
| L SMA |  | -8 | 8 | 50 | 7.44 | < 0.001 |
| L Precentral |  | -35 | -20 | 58 | 7.21 | < 0.001 |
| Vermis 4_5 | 1031 | 2 | -60 | -15 | 6.39 | < 0.001 |
| R Cerebellum 4_5 |  | 10 | -52 | -20 | 5.70 | < 0.001 |
| R Cerebellum 6 |  | 28 | -55 | -28 | 5.59 | < 0.001 |
| L Inferior parietal | 42 | -28 | -48 | 45 | 3.79 | < 0.001 |
| L Superior parietal |  | -22 | -58 | 50 | 3.59 | < 0.001 |
| L Postcentral | 11 | -55 | -20 | 28 | 3.79 | < 0.001 |
| **a2. -Adults (rest > task)** |  |  |  |  |  |  |
| L Inferior frontal, triangular | 5877 | -55 | 28 | 18 | 7.98 | < 0.001 |
| L Superior frontal |  | -18 | 65 | 12 | 6.75 | < 0.001 |
|  |  | -15 | 55 | 30 | 6.63 | < 0.001 |
| L Middle cingulate | 1197 | -5 | -40 | 42 | 7.20 | < 0.001 |
| L Precuneus |  | -5 | -58 | 22 | 4.36 | < 0.001 |
| R Precuneus |  | 12 | -52 | 12 | 3.96 | < 0.001 |
| L Angular | 2206 | -50 | -62 | 48 | 7.17 | < 0.001 |
|  |  | -52 | -70 | 28 | 6.97 | < 0.001 |
| R Parahippocampus | 617 | 30 | -38 | -12 | 6.99 | < 0.001 |
| R Hippocampus |  | 28 | -15 | -20 | 5.33 | < 0.001 |
| R Fusiform |  | 25 | -72 | -8 | 3.92 | < 0.001 |
| R Inferior frontal, triangular | 114 | 58 | 30 | 20 | 5.98 | < 0.001 |
| R Inferior frontal, orbital | 124 | 32 | 35 | -10 | 5.28 | < 0.001 |
| L Hippocampus | 253 | -25 | -18 | -18 | 5.04 | < 0.001 |
| L Fusiform |  | -28 | -45 | -10 | 4.97 | < 0.001 |
| **b. Adults - children** |  |  |  |  |  |  |
| R Superior parietal | 22199 | 35 | -48 | 55 | 10.42 | < 0.001 |
| L Inferior parietal |  | -40 | -38 | 50 | 10.32 | < 0.001 |
| R SMA |  | 15 | -25 | 50 | 8.53 | < 0.001 |
| R Cerebellum 4_5 | 8144 | 20 | -50 | -22 | 7.06 | < 0.001 |
| L Cerebellum 6 |  | -28 | -60 | -22 | 6.68 | < 0.001 |
| L Middle frontal | 77 | -32 | 40 | 18 | 4.06 | < 0.001 |
| R Anterior orbital | 89 | 22 | 52 | -15 | 3.74 | < 0.001 |
| R Middle frontal |  | 35 | 52 | -8 | 3.71 | < 0.001 |
| R Medial orbital |  | 20 | 35 | -18 | 3.64 | < 0.001 |
| L Anterior orbital | 25 | -25 | 48 | -15 | 3.68 | < 0.001 |
| **b1. -Children (rest > task)** |  |  |  |  |  |  |
| L Middle cingulate | 16952 | -12 | -38 | 45 | 8.25 | < 0.001 |
|  |  | 10 | -30 | 48 | 8.18 | < 0.001 |
| L Inferior parietal | 572 | -52 | -55 | 48 | 6.75 | < 0.001 |
| R Supramarginal | 547 | 55 | -45 | 38 | 4.89 | < 0.001 |
| R Inferior parietal |  | 55 | -48 | 50 | 4.59 | < 0.001 |
| R Supramarginal |  | 65 | -25 | 45 | 4.50 | < 0.001 |
| L Middle frontal | 150 | -40 | 18 | 45 | 4.08 | < 0.001 |
|  |  | -25 | 22 | 38 | 3.95 | < 0.001 |
| L Superior frontal | 11 | -15 | 15 | 62 | 3.67 | < 0.001 |
| **b2. Adults (task > rest)** |  |  |  |  |  |  |
| R Precentral | 32667 | 38 | -22 | 58 | 17.72 | < 0.001 |
| L SMA |  | 0 | 0 | 60 | 17.15 | < 0.001 |
| L Postcentral |  | -40 | -35 | 48 | 16.99 | < 0.001 |
| R Middle frontal | 188 | 35 | 40 | 25 | 4.58 | < 0.001 |

Table presents clusters at p_uncorr_ < 0.001. Voxels that were located outside of grey matter are not reported. k = cluster size. Minimum cluster size = 5 voxels. x, y and z coordinates are specified in MNI space. SMA = Supplementary Motor Area. Results presented in this table are based on n = 21 children and 23 adults.

#### Table S25. Regression analyses between brain responses during retest and macro-offline performance changes [macro-offline changes x retest].

| **Area** | **k** | **x** | **y** | **z** | **T** | **p_uncorr_** |
| --- | --- | --- | --- | --- | --- | --- |
| **a. Children - adults** |  |  |  |  |  |  |
| No suprathreshold clusters |  |  |  |  |  |  |
| **a1. Children** |  |  |  |  |  |  |
| No suprathreshold clusters |  |  |  |  |  |  |
| **a2. -Adults** |  |  |  |  |  |  |
| L Postcentral | 25 | -28 | -38 | 50 | 4.18 | < 0.001 |
| R Postcentral | 36 | 20 | -40 | 60 | 3.86 | < 0.001 |
| **b. Adults - children** |  |  |  |  |  |  |
| No suprathreshold clusters |  |  |  |  |  |  |
| **b1. -Children** |  |  |  |  |  |  |
| No suprathreshold clusters |  |  |  |  |  |  |
| **b2. Adults** |  |  |  |  |  |  |
| No suprathreshold clusters |  |  |  |  |  |  |
| **c. ALL** |  |  |  |  |  |  |
| No suprathreshold clusters |  |  |  |  |  |  |
| **d. -ALL** |  |  |  |  |  |  |
| L Postcentral | 53 | -25 | -38 | 50 | 4.60 | < 0.001 |
| R Postcentral | 41 | 18 | -42 | 60 | 3.94 | < 0.001 |

Table presents clusters at p_uncorr_ < 0.001. Voxels that were located outside of grey matter are not reported. k = cluster size. Minimum cluster size = 5 voxels. x, y and z coordinates are specified in MNI space. Results presented in this table are based on n = 21 children and 23 adults. ALL = across all participants, irrespective of age group.

#### Table S26. Additional results of the inter-session changes in brain activity [Retest – Training 2].

| **Area** | **k** | **x** | **y** | **z** | **T** | **p_uncorr_** |
| --- | --- | --- | --- | --- | --- | --- |

| **a. Adults** |  |  |  |  |  |  |
| --- | --- | --- | --- | --- | --- | --- |
| L SMA | 1636 | 0 | -12 | 65 | 5.77 | < 0.001 |
| L Precentral |  | -28 | -30 | 62 | 5.65 | < 0.001 |
| R Postcentral |  | 35 | -38 | 68 | 5.40 | < 0.001 |
| Vermis 6 | 2209 | 2 | -68 | -8 | 5.53 | < 0.001 |
| Vermis 4-5 |  | 0 | -58 | -2 | 5.28 | < 0.001 |
| L Cerebellum 6 |  | -35 | -65 | -25 | 4.99 | < 0.001 |
| L Supramarginal | 578 | -55 | -25 | 25 | 4.79 | < 0.001 |
|  |  | -62 | -38 | 25 | 4.52 | < 0.001 |
|  |  | -60 | -28 | 45 | 3.93 | < 0.001 |
| L Superior parietal | 64 | -22 | -45 | 72 | 4.28 | < 0.001 |
| L Postcentral |  | -45 | -40 | 60 | 3.63 | < 0.001 |
| R Postcentral | 193 | 58 | -22 | 32 | 3.90 | < 0.001 |
| R Cerebellum 9 | 14 | 8 | -50 | -52 | 3.72 | < 0.001 |
| L Precentral | 17 | -58 | 0 | 30 | 3.65 | < 0.001 |
| L Cerebellum crus 1 | 6 | -45 | -48 | -32 | 3.51 | 0.001 |
| L Anterior cingulate | 8 | 0 | 15 | 30 | 3.48 | 0.001 |
| R Supramarginal | 5 | 52 | -35 | 25 | 3.40 | 0.001 |
| **b. Children** |  |  |  |  |  |  |
| R Fusiform | 37 | 32 | -12 | -32 | 5.83 | < 0.001 |
| L Cerebellum 6 | 53 | -5 | -72 | -15 | 3.98 | < 0.001 |
| R SMA | 53 | 2 | -10 | 58 | 3.84 | < 0.001 |
| R Cerebellum 6 | 32 | 30 | -50 | -22 | 3.75 | < 0.001 |
| R Fusiform |  | 28 | -40 | -20 | 3.43 | 0.001 |
| R Supramarginal | 27 | 58 | -38 | 28 | 3.70 | < 0.001 |
| R Supramarginal | 28 | 62 | -18 | 28 | 3.52 | 0.001 |
| L Supramarginal | 56 | -58 | -25 | 28 | 3.43 | 0.001 |
| **c. -Adults** |  |  |  |  |  |  |
| No suprathreshold clusters |  |  |  |  |  |  |
| **d. -Children** |  |  |  |  |  |  |
| L Amygdala | 18 | -22 | -2 | -20 | 4.48 | < 0.001 |
| L Medial superior frontal | 9 | -2 | 50 | 45 | 3.64 | < 0.001 |
| L Hippocampus | 11 | -22 | -25 | -12 | 3.63 | < 0.001 |
| L Middle frontal | 8 | -20 | 15 | 32 | 3.44 | 0.001 |

| **e. ALL** |  |  |  |  |  |  |
| --- | --- | --- | --- | --- | --- | --- |
| Vermis 6 | 1441 | -2 | -70 | -15 | 6.38 | < 0.001 |
|  |  | 2 | -68 | -8 | 5.88 | < 0.001 |
| Vermis 4-5 |  | 0 | -58 | -2 | 5.63 | < 0.001 |
| L SMA | 1244 | 0 | -12 | 60 | 6.29 | < 0.001 |
| L Precentral |  | -28 | -30 | 62 | 5.98 | < 0.001 |
| R Middle cingulate |  | 8 | -28 | 50 | 4.79 | < 0.001 |
| R Postcentral | 643 | 28 | -30 | 65 | 5.80 | < 0.001 |
|  |  | 35 | -35 | 68 | 5.46 | < 0.001 |
| R Superior parietal |  | 30 | -52 | 70 | 4.13 | < 0.001 |
| L Supramarginal | 792 | -58 | -25 | 28 | 5.71 | < 0.001 |
|  |  | -65 | -38 | 28 | 5.06 | < 0.001 |
| L Inferior parietal |  | -60 | -28 | 48 | 4.76 | < 0.001 |
| R Cerebellum 6 | 782 | 25 | -52 | -22 | 5.57 | < 0.001 |
| R Cerebellum crus I |  | 40 | -75 | -20 | 3.91 | < 0.001 |
| R Supramarginal | 705 | 60 | -20 | 28 | 5.09 | < 0.001 |
|  |  | 58 | -35 | 28 | 4.73 | < 0.001 |
| R Fusiform | 25 | 38 | -12 | -30 | 4.92 | < 0.001 |
| L Cerebellum 7 | 106 | -20 | -78 | -50 | 3.88 | < 0.001 |
| L Cerebellum 8 |  | -32 | -62 | -48 | 3.80 | < 0.001 |
| L Cerebellum 7 |  | -8 | -75 | -45 | 3.56 | < 0.001 |
| R Cerebellum 8 | 29 | 28 | -55 | -48 | 3.72 | < 0.001 |
| R Cerebellum crus 2 |  | 30 | -70 | -42 | 3.40 | 0.001 |
| **f. -ALL** |  |  |  |  |  |  |
| No suprathreshold clusters | |  |  |  |  |  |

Table presents clusters at p_uncorr_ < 0.001. Voxels that were located outside of grey matter are not reported. k = cluster size, determined at p_uncorr_ < 0.001. Minimum cluster size = 5 voxels. x, y and z coordinates are specified in MNI space. SMA = Supplementary Motor Area. Results presented in this table are based on n = 18 children and 23 adults. Results of the between-group contrasts are depicted in Table 3a. ALL = across all participants, irrespective of age group.

#### Table S27. Additional results of the regression analysis inter-session changes with the macro-offline performance changes [(retest – training 2) x macro-offline performance changes].

| **Area** | **k** | **x** | **y** | **z** | **T** | **p_uncorr_** | **p_uncorr_** |
| --- | --- | --- | --- | --- | --- | --- | --- |

| **a. Adults** |  |  |  |  |  |  |
| --- | --- | --- | --- | --- | --- | --- |
| No suprathreshold clusters |  |  |  |  |  |  |
| **b. Children** |  |  |  |  |  |  |
| No suprathreshold clusters |  |  |  |  |  |  |
| **c. -Adults** |  |  |  |  |  |  |
| R Medial superior frontal | 46 | 2 | 70 | 10 | 4.07 | < 0.001 |
|  |  | 5 | 68 | 22 | 3.81 | < 0.001 |
| L Medial superior frontal | 11 | -10 | 62 | 32 | 3.85 | < 0.001 |
| **d. -Children** |  |  |  |  |  |  |
| R Precuneus | 924 | 12 | -68 | 28 | 4.94 | < 0.001 |
|  |  | 18 | -55 | 15 | 4.22 | < 0.001 |
| R Medial frontal, orbital | 75 | 10 | 40 | -12 | 4.36 | < 0.001 |
| L Anterior cingulate | 83 | -8 | 35 | 22 | 4.23 | < 0.001 |
| R Caudate | 51 | 10 | 15 | -5 | 4.21 | < 0.001 |
| L Middle cingulate | 246 | -10 | -35 | 48 | 4.12 | < 0.001 |
| L Paracentral lobule |  | -8 | -28 | 75 | 3.73 | < 0.001 |
|  |  | 10 | -30 | 35 | 3.65 | < 0.001 |
| R Postcentral | 87 | 10 | -40 | 70 | 3.88 | < 0.001 |
| R Precuneus |  | 8 | -48 | 65 | 3.85 | < 0.001 |
| L Inferior frontal, triangular | 76 | -52 | 22 | 10 | 3.80 | < 0.001 |
| L Caudate | -14 | -8 | 18 | -5 | 3.79 | < 0.001 |
| R Middle frontal | 30 | 28 | 28 | 42 | 3.78 | < 0.001 |
| L Thalamus | 20 | -12 | -18 | 18 | 3.76 | < 0.001 |
| L Inferior frontal, opercular | 132 | -38 | 20 | 32 | 3.73 | < 0.001 |
|  |  | -35 | 2 | 28 | 3.69 | < 0.001 |
| L Precentral |  | -42 | 8 | 40 | 3.53 | 0.001 |
| L Inferior frontal, triangular | 40 | -35 | 25 | 15 | 3.64 | < 0.001 |
| R Precuneus | 14 | 15 | -45 | 45 | 3.61 | < 0.001 |
| L Fusiform | 12 | -38 | -12 | -20 | 3.52 | 0.001 |
| L Inferior frontal, orbital | 5 | -48 | 40 | -10 | 3.49 | 0.001 |
| L Posterior cingulate | 14 | -8 | -40 | 22 | 3.47 | 0.001 |
| R Caudate | 10 | 15 | 10 | 15 | 3.42 | 0.001 |

| **e. ALL** |
| --- |
| No suprathreshold clusters |
| **f. -ALL** |

| R Medial superior frontal | 110 | 2 | 68 | 12 | 4.27 | < 0.001 |
| --- | --- | --- | --- | --- | --- | --- |
|  |  | 8 | 65 | 22 | 3.77 | < 0.001 |
| R Medial superior frontal | 8 | 5 | 40 | 55 | 3.82 | < 0.001 |
| L Superior frontal | 6 | -20 | 55 | 38 | 3.54 | 0.001 |
| L Superior frontal | 5 | -12 | 52 | 42 | 3.48 | 0.001 |
| R Medial frontal, orbital | 5 | 8 | 38 | -10 | 3.43 | 0.001 |

Table presents clusters at p_uncorr_ < 0.001. Voxels that were located outside of grey matter are not reported. k = cluster size, determined at p_uncorr_ < 0.001. Minimum cluster size = 5 voxels. x, y and z coordinates are specified in MNI space. Results presented in this table are based on n = 18 children and 23 adults. ALL = across all participants, irrespective of age group. Results of the between- and within-group contrasts are depicted in Table 3b.

#### Table S28. Age-related changes in brain responses in the child sample [task practice vs. interleaved rest].

| **Area** | **k** | **x** | **y** | **z** | **T** | **p_uncorr_** |
| --- | --- | --- | --- | --- | --- | --- |
| **a. Training 1** |  |  |  |  |  |  |
| Children |  |  |  |  |  |  |
| No suprathreshold voxels |  |  |  |  |  |  |
| -Children |  |  |  |  |  |  |
| R Precentral | 37 | 42 | -5 | 38 | 3.70 | < 0.001 |
| L Postcentral | 9 | -62 | -10 | 30 | 3.56 | < 0.001 |
| **b. Training 2** |  |  |  |  |  |  |
| Children |  |  |  |  |  |  |
| No suprathreshold voxels |  |  |  |  |  |  |
| -Children |  |  |  |  |  |  |
| L Inferior frontal, opercular | 33 | -48 | 5 | 5 | 3.72 | < 0.001 |
| L Postcentral | 14 | -62 | -10 | 28 | 3.71 | < 0001 |
| L Middle cingulate | 30 | -10 | 22 | 32 | 3.68 | < 0.001 |
| **c. Retest** |  |  |  |  |  |  |
| Children |  |  |  |  |  |  |
| No suprathreshold voxels |  |  |  |  |  |  |
| -Children |  |  |  |  |  |  |
| R Superior frontal | 35 | 20 | 62 | 0 | 4.24 | < 0.001 |
| **d. Training 2 – Training 1** |  |  |  |  |  |  |
| Children |  |  |  |  |  |  |
| R Medial superior frontal | 8 | 12 | 60 | 0 | 3.51 | 0.001 |
| -Children |  |  |  |  |  |  |
| L Hippocampus | 13 | -32 | -15 | -20 | 3.80 | < 0.001 |
| **e. Retest – Training 2** |  |  |  |  |  |  |
| Children |  |  |  |  |  |  |
| No suprathreshold voxels |  |  |  |  |  |  |
| -Children |  |  |  |  |  |  |
| No suprathreshold voxels |  |  |  |  |  |  |

Table presents clusters at p_uncorr_ < 0.001. Voxels that were located outside of grey matter are not reported. k = cluster size. Minimum cluster size = 5 voxels. x, y and z coordinates are specified in MNI space.

### APPENDIX 6: Supplementary discussion

#### 6.1 Behavioral initial learning dynamics

Although both children and adults learned the novel motor sequence, there was evidence demonstrating that performance differed between the two age groups. Specifically, and in contrast with our previous research (Van Roy et al., 2024), significant group differences were observed in normalized performance speed during both training runs, with slower performance in children. This indicates that in children, the median performance of each block showed less improvement relative to the first training block (i.e., the block used for normalization) as compared to adults. Importantly, the omnibus test revealed no significant group x block interactions, demonstrating that block-to-block changes were similar between age groups. Visual inspection of Figure 3 in the main text suggests that performance improvements from the first to second practice block in training 1 were larger in adults than children. This created a gap in performance that then was maintained for the remaining blocks of the training runs. We speculate that this performance difference may be attributed to the task design. Specifically, the current study employed a bimanual FTT during which participants pressed the keys in the order of a sequence of numbers that was displayed on the screen. It is possible that this explicit mapping of the numeric sequence shown on screen (i.e., 4-2-1-3-4) to the specific keys/fingers to be pressed was more difficult for children and thus minimized performance improvements at the beginning of training. It is worth noting that our experimental procedures aimed to facilitate this sequence-key mapping by implementing several familiarization tasks prior to the motor sequence learning session. Future research should consider extending these familiarization runs to minimize the potential impact of age-related differences in acquiring this mapping on motor sequence learning performance. Importantly, while the gap in performance between age groups remained during training run 2, it was no longer significant in the post-learning test. This suggests that children were able to “catch up” to the performance level of adults over the course of the initial learning session.

#### 6.2 Micro-offline performance changes

In line with previous literature (Du et al., 2017; Van Roy et al., 2024) and our hypothesis, children exhibited a larger magnitude of micro-offline performance changes relative to adults. However, the group effect within the group x training run ANOVA was not significant (p = 0.061) and the corresponding Bayes Factor (BF_10_ = 0.869) provided anecdotal evidence in support of no group differences (Wagenmakers et al., 2011). In our previous study (Van Roy et al., 2024), we also found a non-significant trend toward larger micro-offline performance changes in children, with a Bayes Factor that was consistent with no group differences. Taken together, results from both of our studies did not provide strong evidence for a childhood advantage in micro-offline performance improvements. Our previous research (Van Roy et al., 2024) includes an expanded discussion of potential explanations for this difference. In brief, this discussion focused on the specific ages of the participants (i.e., Du and colleagues acquired data from children younger than in our research) and differences in experimental design (i.e., duration of the interleaved rest periods). Nevertheless, both children and adults demonstrated significantly negative micro-online and positive micro-offline performance changes (see Appendix 3), suggesting that performance improvements during training were largely due to micro-offline performance changes in both age groups.

#### 6.3 Potential explanations for the smaller modulations in brain activity between task and rest epochs in children

Altogether, results suggest that children exhibit smaller modulations in brain activity between task and rest epochs. That is, the magnitude of activations in task-relevant regions (e.g., M1, SMA, putamen, globus pallidus and cerebellum) and the magnitude of *de*activations in regions commonly associated with the default mode network are smaller in children (i.e., betas closer to 0). We thus speculate that the child’s brain stays more engaged during the short rest breaks relative to adults. Several potential explanations could contribute to this continued engagement of the developing brain during interleaved rest. One possibility is that the activation of brain regions during interleaved rest reflects active consolidation processes similar to the previously demonstrated spontaneous reactivation of task-related activity patterns during the short rest periods in adults (Buch et al., 2021; Gann et al., 2023). Although this explanation is certainly possible, it likely cannot explain the full set of results presented above. Specifically, the reactivation framework posits that the patterns of activity observed during learning are spontaneously reactivated during subsequent rest intervals in *specific* brain regions known to be vital for learning that particular task. Given that the current results suggest continued engagement during interleaved rest intervals across such a widespread network of regions, many of which would not be considered task-relevant (e.g., DMN), it does not appear that reactivation would be the sole contributing process. Alternatively, Camacho et al. (2020) proposed that the “inability to rest” of the developing brain could be explained by two potential factors. On one hand, they suggested that children are focused on laying still during the rest periods while they are in the scanner. As executive functions (e.g., shifting, updating, inhibiting) continue to develop across childhood (Anderson, 2002; Luna et al., 2004), the task to remain still may induce an increased demand for inhibitory control. On the other hand, Camacho et al. (2020) hypothesized that children utilize the short rest breaks to cognitively prepare themselves for the upcoming practice block. Such an explanation would also be consistent with recent research in adults by Das and colleagues (2024), which suggested that interleaved rest breaks serve as an opportunity to plan movements in the upcoming practice block. In the current study, relatively comparable activation levels across task and rest blocks in task-relevant regions were only observed in children, potentially suggesting that they demonstrate greater preparation/planning during interleaved rest intervals than adults. Similar to the reactivation explanation above, it is not clear how this notion of pre-planning could explain the continued engagement during rest in children across such a widespread network of regions. Although the current experimental design and analytic approach does not allow us to distinguish between these various possible explanations, our results do suggest that brain activity during the interleaved rest epochs – as opposed to solely during task practice - may contribute to the observed age-related differences.

#### 6.4 The relationship between brain activity during initial learning and micro-offline performance changes

As the current study aimed to identify the functional neural correlates underlying the childhood advantage in offline processing, we conducted regression analyses that assessed the relationship between brain activation during initial training (i.e., training 1) and the micro-offline performance changes. Prior research in adults has shown a link between micro-offline learning and neural activity in subcortical regions during the interleaved rest periods (Buch et al., 2021; Gann et al., 2023; Jacobacci et al., 2020). In contrast to our hypothesis, we did not find any significant relationships between activity during training 1 and the micro-offline performance changes across all participants, nor did we observe any differences between age groups. The reasons for the lack of such a brain-behavior relationship are not entirely clear, but it is worth noting that our behavioral results revealed only non-significantly greater micro-offline performance changes in children, with a relatively small Bayes Factor that is indicative of no differences between children and adults. Thus, the absence of any group differences in the relationships between neural activity and micro-offline performance changes may mirror the lack of a robust difference at the behavioral level.

#### 6.5 Inter-session changes in brain activity in young adults

The assessment of changes in brain activity across the 5-hour offline period (i.e., from training 2 to retest) revealed that adults showed significant increases in activation of M1 and the cerebellum. While the cerebellum is traditionally seen as a key player in motor coordination, research has indicated that the cerebellar cortex is also involved in cognitive control (Saadon-Grosman et al., 2024; Stoodley et al., 2012). Furthermore, the cerebellar cortex has also been linked to the early phase of MSL processes that takes place under high cognitive control and typically demonstrates decreases in activity across a consolidation interval in healthy adults (Doyon et al., 2002). The observed increase in activity in the cerebellar cortex (i.e., cerebellum lobule 4-5, lobule 9 and crus I) from training 2 to retest in the adults was thus unexpected. Given the role of the cerebellar cortex in early learning, one could speculate that higher cerebellar recruitment during retest in adults was necessary to re-learn the sequence after their offline deterioration in performance, reflecting poor consolidation of the motor memory. However, the observed inter-session increases in M1 activity in adults would suggest the opposite. Specifically, prior research in adults found that sleep-dependent enhancements in performance were paralleled by increases in M1 activity (Debas et al., 2010). While adults exhibited an offline deterioration in performance, the observed inter-session increase in activity may thus suggest at least a partial stabilization of the memory representation of the sequence across the consolidation interval.
